# Zinc availability in the tumor microenvironment dictates anti-PD1 response in *CDKN2A*^Low^ tumors via increased macrophage phagocytosis

**DOI:** 10.1101/2025.02.08.637227

**Authors:** Raquel Buj, Aidan R. Cole, Jeff Danielson, Jimmy Xu, Drew Hurd, Akash Kishore, Katarzyna M. Kedziora, Jie Chen, Baixue Yang, David Barras, Apoorva Uboveja, Amandine Amalric, Juan J. Apiz Saab, Jayamanna Wickramasinghe, Naveen Kumar Tangudu, Evan Levasseur, Hui Wang, Aspram Minasyan, Rebekah E. Dadey, Allison C. Sharrow, Sheryl Kunning, Frank P. Vendetti, Dayana B. Rivadeneira, Christopher J. Bakkenist, Tullia C. Bruno, Greg M. Delgoffe, Nadine Hempel, Nathaniel W. Snyder, Riyue Bao, Adam C. Soloff, John M. Kirk-wood, Denarda Dangaj Laniti, Andrew V. Kossenkov, Alexander Muir, Jishnu Das, Diwakar Davar, Clementina Mesaros, Katherine M. Aird

## Abstract

Anti-PD1 therapies are primarily thought to rely on functional T cell responses; yet tumors with limited T cell infiltration can still benefit, suggesting alternative mechanisms contribute to therapeutic efficacy. Indeed, we found that myeloid-rich, T cell-poor tumor models respond to anti-Pd1, and this is dependent on a cancer cell-macrophage crosstalk mediated by cancer cell *Cdkn2a* expression. Mechanistically, we found that cancer cells with decreased *Cdkn2a* expression (C*dkn2a*^Low^), which occurs in ∼50% of all human cancers, reorganize zinc compartmentalization by upregulating the zinc importer Slc39a9 at the plasma membrane. Increased cancer cell plasma membrane Slc39a9 leads to intracellular zinc accumulation in cancer cells and depletion of zinc in the tumor microenvironment (TME), resulting in zinc-starved tumor-associated macrophages (TAMs) with reduced phagocytic activity. Restoring zinc availability in TAMs—via dietary supplementation or Slc39a9 knockdown in cancer cells—reprograms TAMs to a pro-phagocytic state and sensitizes *Cdkn2a*^Low^ tumors to anti-Pd1 therapy. Remarkably, Slc39a9 knockdown tumors respond to anti-Pd1 in Rag1^−/−^ mice, and co-injection of zinc-replete macrophages is sufficient to drive an anti-Pd1 response in immunodeficient mice, demonstrating the T cell-independent nature of this response. Clinically, TAMs from *CDKN2A*^Low^ cancer patients show reduced zinc and phagocytosis gene signatures. Moreover, patients with lower circulating zinc levels have significantly worse time-to-event outcomes than those with higher levels. Together, these findings uncover a previously unrecognized mechanism by which *Cdkn2a*^Low^ cancer cells outcompete TAMs for zinc, impairing their function and limiting anti-Pd1 efficacy. They also provide evidence that macrophages alone, without T cells, can enhance anti-PD1 response through zinc-mediated reprogramming of phagocytosis.

## INTRODUCTION

A number of data mining studies have highlighted a significant correlation between Cyclin-Dependent Kinase Inhibitor 2A gene (*CDKN2A*) genetic alterations and a pro-tumorigenic tumor microenvironment (TME), including TMEs with decreased T cell infiltration^1–6^. Prior studies have also shown a correlation between loss of *CDKN2A* function and poor response to immune checkpoint blockade (ICB) therapies^1–18^. ICB therapies, particularly monoclonal antibodies targeting the PD-1/PD-L1 axis, have become a cornerstone in cancer treatment, demonstrating significant clinical success across several cancers^19–23^. Their effectiveness is primarily attributed to enhancing T cell activity and T cell-mediated tumor killing^24^. However, published studies have described specific contexts in which tumors with low T cell infiltration but high myeloid cell content show responsiveness to immunotherapies, including ICB^25–28^. Supporting this notion, recent evidence suggests that at least a subset of tumor-associated macrophages (TAMs) play a role in the response to ICB^28–35^. Anti-PD-1/PD-L1 therapies can directly affect monocytes and macrophages to improve therapeutic responses at least in part by increasing their phagocytic activity^28,29,36–38^. Indeed, studies have shown that increased TAM phagocytic activity inhibits tumor growth in mouse models^39–41^. However, the mechanisms underlying this enhancement of TAM phagocytosis remain unclear.

The *CDKN2A* gene, located on chromosome 9p21, is a critical tumor suppressor and one of the most frequently altered loci in human cancers^42–45^. Disruption of *CDKN2A* typically occurs through promoter hypermethylation and genetic deletions, resulting in the loss of *CDKN2A* mRNA expression^46–48^. These alterations are observed in 12% to 80% of tumors, depending on cancer type and stage^49^. Additionally, loss-of-function mutations in the *CDKN2A* coding sequence represent another common mechanism of disruption, occurring in approximately 6% to 16% of cases, with frequencies also depending on tumor context^48,49^. *CDKN2A* encodes two distinct proteins through alternative reading frames: p14^ARF^ (p19^ARF^ in mice) and p16^INK4A^ (hereafter p16), both of which are essential regulators of the cell cycle^50^. While the canonical roles of *CDKN2A* in cell cycle regulation and senescence induction are well established^51,52^, evidence from our laboratory and others suggests that its functions extend beyond this paradigm^13,18,53–63^. We have demonstrated that loss of *CDKN2A*/p16 expression reprograms intracellular metabolism, enhancing key biosynthetic metabolites that contribute to cellular proliferation and tumorigenesis, including nucleotides and cholesterol^54,56^. However, whether this metabolic disruption downstream of *CDKN2A* loss affects the metabolic landscape of the TME remains unexplored. Additionally, while metabolic reprogramming is known to influence macrophage phagocytic activity^39,64,65^, the role of cancer cell-TAM nutrient competition in driving this process remains poorly understood.

Zinc is an essential trace element obtained through the diet^66^ and is the second most abundant metal in the human body^67^. Over 2,500 human proteins depend on zinc for structural or functional roles (e.g. zinc fingers and cofactors)^67^, reflecting its broader importance in numerous cellular processes, including immune responses^68–75^. Therefore, tight regulation of zinc homeostasis is essential. The SLC39A family members, also known as ZRT-and IRT-like proteins (ZIP), increase cytosolic zinc by importing it from the extracellular environment or releasing it from intracellular organelles^76^. In contrast, the SLC30A family members or zinc transporter (ZNT) proteins, reduce cytosolic zinc by transporting it into organelles or exporting it to the extracellular space^77^. Indeed, dysregulated zinc homeostasis, due to altered transport or systemic deficiency, has been linked to multiple malignancies^78–90^. Moreover, *in vivo* studies have shown that zinc supplementation can reduce tumor burden, enhance resistance to tumor challenges, and decrease the incidence of spontaneous tumors in mice^91–94^, suggesting a potential therapeutic role for zinc in cancer treatment. Additionally, recent reports have found that patients who respond to ICB have TAMs with metal/zinc signatures^30^, and an elevated serum zinc level has recently been associated with improved ICB responses in cancer patients^95^. While some studies have explored the link between zinc and immunotherapy response^96–100^, the results remain contradictory, and the precise molecular mechanisms underlying zinc in the TME are unclear.

Here, we found that in myeloid-rich, T cell-poor tumors, the expression of *Cdkn2a* in cancer cells determines the response to anti-Pd1 by regulating zinc compartmentalization, which in turn modulates TAM phagocytic activity. By cross-comparing differentially regulated metabolites and metals in the tumor interstitial fluid (TIF) and negatively enriched plasma membrane solute carrier transporters (SLCs) from a metabolism-focused *in vivo* CRISPR knockout screen, we identified zinc and the zinc importer Slc39a9 as critical regulators of anti-Pd1 resistance in tumors with low *Cdkn2a* mRNA expression (*Cdkn2a*^Low^). *Cdkn2a*^Low^ cells have increased cholesterol-driven plasma membrane (PM) localization of Slc39a9, leading to elevated intracellular zinc and a zinc-depleted TME. This competition for zinc within the TME leads to decreased zinc in TAMs, reducing their phagocytic activity and impairing anti-Pd1 response. Restoring TME zinc levels through Slc39a9 knockdown in cancer cells or dietary zinc supplementation rescues TAM function and enhances anti-Pd1 response. Excitingly, this was independent of T cells. Using publicly available single-cell RNA-Sequencing (scRNA-Seq), we also found that zinc and phagocytosis signatures were impaired in TAMs from patients with *CDKN2A*^Low^ tumors. These findings highlight zinc homeostasis as essential for anti-PD1 efficacy in *CDKN2A*^Low^ tumors, emphasize T cell-independent mechanisms of anti-PD1 response, and have clinical implications for the use of zinc supplements to improve patient outcomes.

## RESULTS

### Expression of *CDKN2A*/p16 in cancer cells modulates anti-PD1 response in myeloid-rich, CD8 T cell-poor tumors

Loss of *CDKN2A* is a common event in cancer cells^49^, including melanoma, where ∼40% of patients show reduced *CDKN2A* expression^101,102^. Published studies show that tumors with loss of *CDKN2A* correlate with decreased response to ICB, including anti-PD1^1,7,9,12,14–17^. Consistently, using publicly available data, we found that melanoma patients with low *CDKN2A* mRNA expression (*CDKN2A*^Low^) have shorter overall survival compared to those with high *CDKN2A* mRNA expression (*CDKN2A*^High^) following anti-PD1 treatment (**Fig. 1A**). Therefore, we sought to determine the mechanisms by which *CDKN2A* loss in cancer cells contributes to anti-PD1 resistance. Towards this goal, we developed two complementary syngeneic models. In the first model, *Cdkn2a* expression was knocked down in the *Cdkn2a*^High^ murine melanoma cell line Yumm5.2 (**Fig. S1A**). In the second model, p16 was expressed at physiological levels in the *Cdkn2a*^Null^ melanoma cell line Yumm1.7 (**Fig. S1B**). Note that we did not observe differences in cell proliferation or senescence upon physiological expression of p16 (**Fig. S1C-D**). Additionally, p19 remained unexpressed in the Yumm1.7 cells with p16 expression (**Fig. S1B**); thus, results with this model specifically relate to p16 function. *CDKN2A* is located on chromosome 9p21 in humans and 4qC4 in mice. In both species, this region also contains *MTAP* and the interferon (IFN) gene cluster, which are frequently co-deleted with *CDKN2A* in cancer^1,103,104^. Importantly, both of our models have intact *Mtap* and *Ifn* gene cluster (**Fig. S1E-F**); therefore, all subsequent observations with these models are related only to *Cdkn2a*. Herein, for simplicity, Yumm5.2 and Yumm1.7 with p16 expression are referred to as *Cdkn2a*^High,^ while Yumm5.2 with knockdown of *Cdkn2a* and Yumm1.7 cells are referred to as *Cdkn2a*^Low^ cancer cells (**Fig. S1G**). Importantly, both syngeneic tumors mimic the ICB response seen in cancer patients, where decreased mRNA expression of *Cdkn2a* in both models leads to anti-Pd1 resistance (**Fig. 1B-C**).

**Figure 1.**
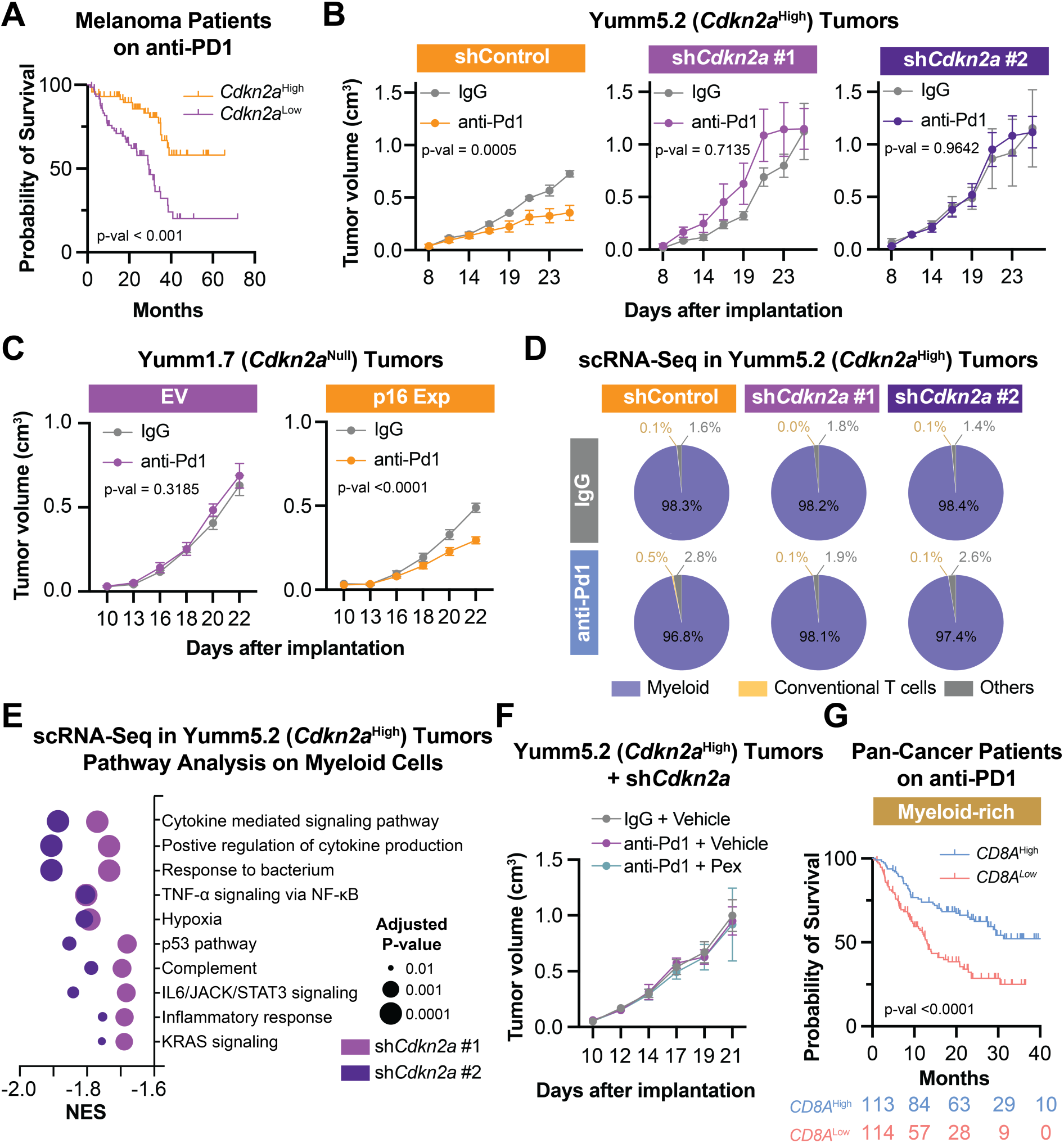
Loss of *CDKN2A* expression abrogates anti-PD1 efficacy in myeloid-rich, CD8 T cell-poor tumors. **(A)** Overall survival of melanoma patients treated with anti-PD1 stratified by *CDKN2A* mRNA expression. Melanoma Patient Cohort; Log-rank (Mantel-Cox) test. **(B)** Longitudinal tumor growth in mice implanted with Yumm5.2 (*Cdkn2a*^High^) cells with knockdown of GFP (shControl) or *Cdkn2a* (sh*Cdkn2a,* hairpins #1 and #2) and treated with IgG or anti-Pd1 (200 µg, i.p., 3×/week). Mean ± SEM (n=4/group); mixed-effects model. **(C)** Longitudinal tumor growth in mice implanted with Yumm1.7 (*Cdkn2a*^Null^) cells expressing empty vector (EV) or physiological levels of p16 (p16 Exp) and treated with IgG or anti-Pd1 (200 µg, i.p., 3×/week). Mean ± SEM (n=8/group); mixed-effects model. **(D-E)** Cd45^+^ cells from the indicated Yumm5.2 (*Cdkn2a*^High^) tumors treated with IgG or anti-Pd1 (200 µg, i.p., 3×/week) were sorted and used for scRNA-Seq. **(D)** Pie charts showing percentages of immune cell types. **(E)** Dot plot showing the top 10 significant gene signatures from gene set enrichment analysis (GSEA) performed in the myeloid subset. NES = normalized enrichment score. **(F)** Longitudinal tumor growth in mice implanted with Yumm5.2 (*Cdkn2a*^High^) cells with knockdown of *Cdkn2a* and treated with pexidartinib (Pex, 45 mg/kg, o.g., 3×/week) or vehicle in combination with IgG or anti-Pd1 (200 µg, i.p., 3×/week). Mean ± SEM (n=6/group); mixed-effects model. **(G)** Overall survival of anti-PD1-treated, myeloid-rich (*ITGAM* mRNA expression above the group average) cancer patients stratified by *CD8A* mRNA expression. KMP Patient Cohort; Log-rank (Mantel-Cox) test.

Given the central role of the tumor immune microenvironment in dictating ICB outcomes^105^, we next investigated how *Cdkn2a* expression in cancer cells shapes the immune compartment. Towards this goal, we performed scRNA-Seq on Cd45⁺ immune cells isolated from the Yumm5.2 isogenic model in mice treated with and without anti-Pd1. We found that these tumors are heavily enriched in myeloid cells (>95% of all Cd45^+^ cells) and poor in conventional T cells (<1% of all Cd45^+^ cells), regardless of *Cdkn2a* status or anti-Pd1 treatment (**Fig. 1D** and **S1H**). This myeloid-rich, T cell–poor composition aligns with prior reports showing that *Cdkn2a*^Low^ tumors show reduced T cell infiltration^1–6^ and reflects the immune landscape observed in many human tumors^106^. Intriguingly, despite the paucity of T cells, shControl (*Cdkn2a*^High^) tumors still respond to anti-Pd1 (**Fig. 1B-C**). Thus, these data point to a potential T cell-independent mechanism of anti-Pd1 response in these tumors, which we aimed to explore. Interestingly, Gene Set Enrichment Analysis (GSEA) of scRNA-seq data from the myeloid compartment showed broadly reduced inflammatory signals in Yumm5.2 sh*Cdkn2a* compared to shControl tumors (**Fig. 1E**). This is consistent with prior research showing that anti-inflammatory myeloid cells have protumorigenic functions that suppress ICB responses^107–109^. Therefore, we aimed to determine whether depletion of myeloid cells in *Cdkn2a*^Low^ tumors would phenocopy the improved response to anti-Pd1 observed in *Cdkn2a*^High^ tumors. To test this, we treated mice bearing Yumm5.2 sh*Cdkn2a* tumors with pexidartinib, a CSF1R inhibitor known to effectively reduce tumor myeloid cells^110,111^. Unexpectedly, myeloid depletion (**Fig. S1I**) failed to sensitize sh*Cdkn2a* tumors to anti-Pd1 (**Fig. 1F**). These findings indicate that the overall anti-inflammatory bulk myeloid compartment is not the primary driver of anti-Pd1 resistance in *Cdkn2a*^Low^ tumors.

CD8 T cells are commonly considered the main mediators of anti-PD1 responses^112,113^. However, both *Cdkn2a*^High^ Yumm tumor models respond to anti-Pd1 treatment (**Fig. 1B-C**) despite being Cd8 T cell-poor – particularly Yumm5.2 tumors, which are severely depleted (**Fig. 1D, S1H, and S1J**). Interestingly, there is literature to demonstrate that specific myeloid subsets also contribute to the therapeutic efficacy of ICB^28,30,35,114^. In this regard, analysis of publicly available bulk RNA-seq data from anti-PD1-treated patients showed that 21% of tumors with high *ITGAM* (encoding for CD11B) and low *CD8A* expression (reflecting a myeloid-rich, T cell-poor TME) had similar survival at 20 months to tumors with both high *ITGAM* and high *CD8A* (reflecting a myeloid-rich, T cell-rich TME; **Fig. 1G**). Surprisingly, 7.9% of the myeloid-rich, T cell-poor group maintained similar survival even at 30 months (**Fig. 1G**). Together, these data suggest that a subset of myeloid cells may actively contribute to the anti-PD1 response, even in a T cell-poor environment. In this study, we took advantage of the Yumm models as they provide a unique platform to investigate how myeloid cells, in the context of high *Cdkn2a,* may promote anti-Pd1 responses in a T cell-poor TME.

### *Cdkn2a*^Low^ tumors have impaired anti-Pd1-mediated TAM phagocytosis

We found that depletion of the bulk myeloid compartment does not sensitize T cell-poor sh*Cdkn2a* Yumm5.2 tumors to anti-Pd1 (**Fig. 1F** and **S1I**) and that some patients with myeloid-riched, T cell-poor tumors show sustained benefit from anti-PD1 (**Fig. 1G**). Together, these data suggest that a specific subset of myeloid cells may unexpectedly promote, not inhibit, an anti-PD1 response. To explore this idea, we further investigated the different types of myeloid cells in our scRNA-Seq. We found that the large majority of myeloid cells in these tumors are tumor-associated macrophages (TAMs) (**Fig. 2A**). Interestingly, a growing body of recent literature has reported that TAMs alone drive response to ICB by promoting phagocytosis^28,29,36–38^. Consistently, analysis of publicly available patient data shows that when phagocytosis is high, overall survival following anti-PD1 treatment is not significantly different between T cell-rich and T cell-poor tumors (**Fig. 2B**). Therefore, we sought to determine the role of anti-Pd1-mediated TAM phagocytosis in our models. Further analysis of our scRNA-seq revealed a significant increase in phagocytosis-related gene signatures in TAMs following anti-Pd1 treatment in Yumm5.2 shControl (*Cdkn2a*^High^) but not in sh*Cdkn2a* (*Cdkn2a*^Low^) tumors (**Fig. 2C** and **Table S1**). Notably, even in IgG treated mice, Yumm5.2 shControl tumors showed modestly higher phagocytic activity scores in TAMs compared to Yumm5.2 sh*Cdkn2a* tumors (**Fig. 2C** and **Table S1**), suggesting that TAMs in *Cdkn2a*^Low^ tumors impair basal TAM phagocytosis that is further exacerbated by anti-Pd1. To assess whether this is relevant in human tumors, we analyzed publicly available scRNA-seq data from patients with melanoma and head and neck squamous cell carcinoma (HNSCC), cancers where *CDKN2A* loss is common and clinically relevant^49^. Consistent with our mouse tumor scRNA-Seq data, we observed reduced TAM phagocytosis signatures in *CDKN2A*^Low^ vs. *CDKN2A*^High^ patient tumors (**Fig. 2D**), underscoring the clinical relevance of our findings. To functionally validate these observations, we generated fluorescently-labeled cancer cells and assessed *in vivo* and *in vitro* phagocytosis by measuring the percentage of Cd11b⁺ cells that have engulfed fluorescently-labeled cancer cells (**Fig. 2E** and **Fig. S2**), similar to previous reports^29,115^. We acknowledge the limitation that Cd11b is not a specific TAM marker, as other myeloid cells can also express Cd11b. However, given that the overwhelming majority (60-90%) of phagocytic myeloid cells in these tumors are TAMs (**Fig. 2A**), the Cd11b⁺ population primarily reflects TAMs in these tumors. Consistent with our scRNA-Seq gene signatures, we observed that TAMs in both *Cdkn2a*^High^ tumor models were more phagocytic than those in *Cdkn2a*^Low^ tumors (**Fig. 2F-G**), although we note this was more pronounced in the Yumm5.2 model. Moreover, anti-Pd1 treatment enhanced TAM phagocytosis only in *Cdkn2a*^High^ tumors in both models (**Fig. 2F-G**). This increase in phagocytic activity in response to anti-Pd1 aligns with the tumor growth reduction observed upon anti-Pd1 treatment in both models (**Fig. 1B-C**). Together with patient data, these *in vivo* data demonstrate that TAM phagocytosis is modulated by *CDKN2A* status in cancer cells. Finally, we aimed to confirm that this is a direct effect on macrophages by performing *in vitro* coculture assays with cancer cells and IFNγ-activated RAW264.7 macrophages (**Fig. 2E**). Consistent with our *in vivo* results, macrophages *in vitro* had higher phagocytic activity towards *Cdkn2a*^High^ cancer cells, which was further enhanced by anti-Pd1 (**Fig. 2H**). This effect was not observed when these macrophages were co-cultured with *in vitro Cdkn2a*^Low^ cancer cells (**Fig. 2H**). Collectively, these data demonstrate that *Cdkn2a* expression in cancer cells directly regulates macrophage phagocytic activity and that anti-Pd1 enhances macrophage phagocytosis, which corresponds to decreased tumor volume. These findings add to emerging evidence challenging the prevailing view that T cells are always required to mediate responses to anti-PD1, instead highlighting a critical role for TAMs.

**Figure 2.**
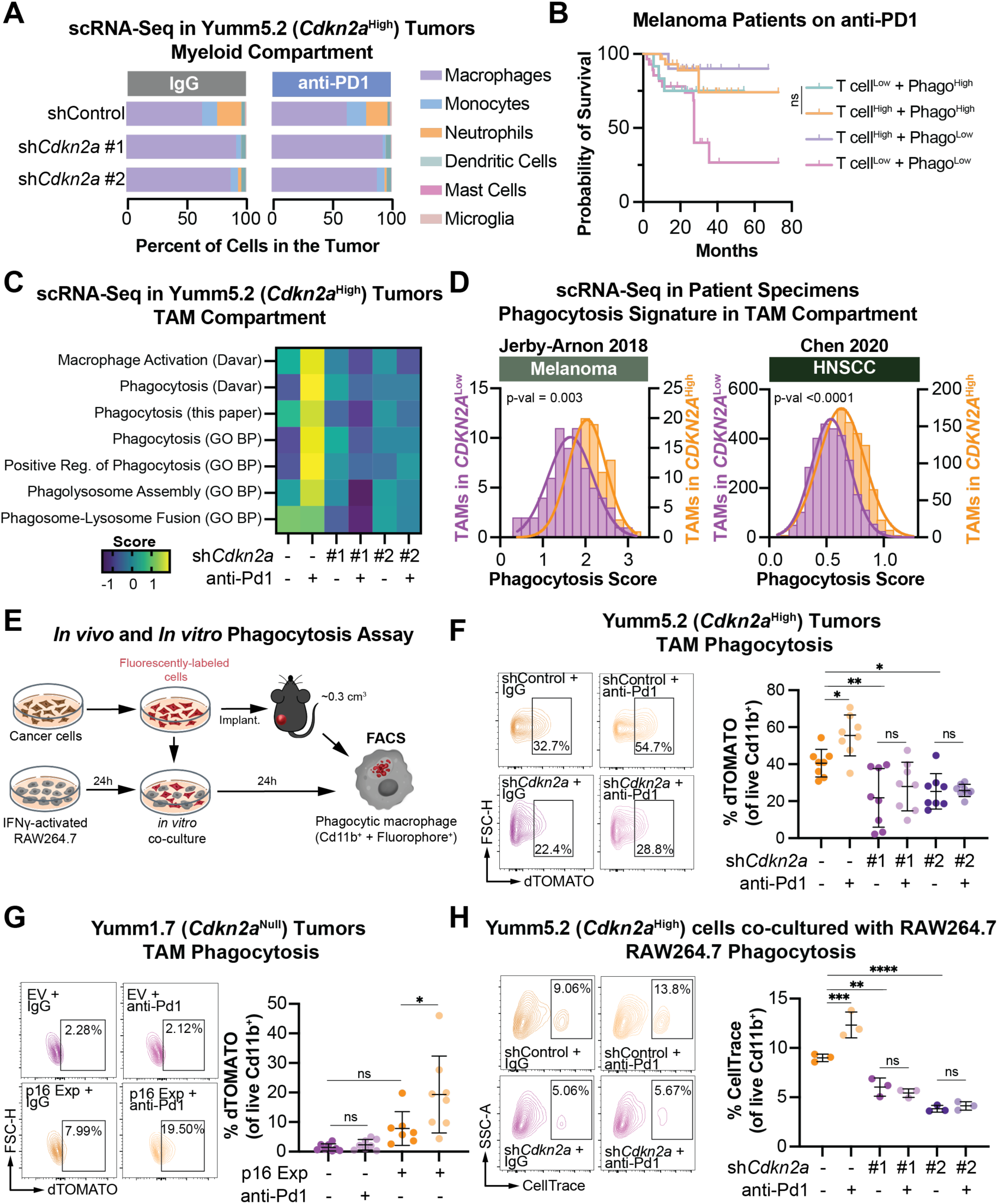
Loss of *CDKN2A* expression in cancer cells impairs tumor-associated macrophage (TAM) phagocytic activity. **(A)** Cd45^+^ cells from the indicated Yumm5.2 (*Cdkn2a*^High^) tumors treated with IgG or anti-Pd1 (200 µg, i.p., 3×/week) were sorted and used for scRNA-Seq. Bar plots showing percentages of myeloid cell types. **(B)** Overall survival of anti-PD1-treated patients stratified by T cell inflamed score and phagocytic score (see **Table S1**). Melanoma Patient Cohort; Log-rank (Mantel-Cox) test. **(C)** Heatmap of phagocytosis signature scores in the TAM compartment from scRNA-seq of Yumm5.2 (*Cdkn2a*^High^) tumors from (A)**. (D)** Histogram showing the phagocytosis score in TAMs from the indicated scRNA-Seq datasets from melanoma and head and neck squamous cell carcinoma (HNSCC) patients stratified by *CDKN2A* expression in cancer cells. Linear mixed model. **(E)** Schematic of *in vivo* and *in vitro* phagocytosis assays. **(F)** dTOMATO-labeled Yumm5.2 (*Cdkn2a*^High^) cells with knockdown of GFP (shControl) or *Cdkn2a* (sh*Cdkn2a,* hairpins #1 and #2) were injected into mice (see schematic in panel E), and mice were treated with IgG or anti-Pd1 (200 µg, i.p., 3×/week). Representative flow cytometry plots (left) and quantification (right) of the percentage of TAMs with dTOMATO signal (phagocytic TAMs). Mean ± SD; one-way ANOVA. **(G)** dTOMATO-labeled Yumm1.7 (*Cdkn2a*^Null^) cells expressing empty vector (EV) or physiological levels of p16 (p16 Exp) were injected into mice (see schematic in panel E), and mice were treated with IgG or anti-Pd1 (200 µg, i.p., 3×/week). Representative flow cytometry plots (left) and quantification (right) of phagocytic TAMs. Mean ± SD; one-way ANOVA. **(H)** CellTrace™ Violet-labeled Yumm5.2 (*Cdkn2a*^High^) cells with shControl or sh*Cdkn2a* knockdown were co-cultured with RAW264.7 macrophages. Cells in co-culture were treated with IgG or anti-Pd1 (1 µg/mL, 24 h). Representative flow cytometry plots (left) and quantification (right) of the percentage of RAW264.7 with CellTrace™ Violet signal (phagocytic RAW264.7 macrophages). Mean ± SD; one-way ANOVA. * p<0.05, ** p<0.01, *** p<0.001, **** p<0.0001, ns = not significant.

### The zinc importer Slc39a9 in cancer cells abrogates anti-Pd1 response in *Cdkn2a*^Low^ tumors via a TAM-dependent mechanism

We found that *Cdkn2a*^High^ cancer cells promote increased TAM phagocytosis compared to *Cdkn2a*^Low^ cancer cells both *in vitro* and *in vivo* and that anti-Pd1 treatment further enhances this effect only in the context of *Cdkn2a*^High^ cells (**Fig. 2C-D and F-H**). TAMs are highly plastic cells and responsive to the tumor’s metabolic environment^116^. We and others have published that suppression of *CDKN2A*/p16 alters cell-intrinsic metabolism^54,55,58,59,117–119^, although no study to date has assessed TME metabolites in the context of *CDKN2A*^Low^ tumors. Analyzing the tumor interstitial fluid (TIF) by unbiased metabolomics and metalomics (**Fig. S3A**), we found that multiple analytes are also differentially regulated in the TME of Yumm5.2 sh*Cdkn2a* tumors compared to controls (**Fig. 3A**). Therefore, we hypothesized that these metabolic changes could extend beyond cancer cells to influence the surrounding immune microenvironment, including TAMs. To identify metabolic drivers of the altered TME in sh*Cdkn2a* tumors and evaluate their role in anti-Pd1 resistance, we performed an *in vivo* CRISPR screen using a metabolism-focused sgRNA knockout library in the Yumm5.2 isogenic model, with and without anti-Pd1 treatment (**Fig. 3B**). We identified 137 genes with significant negative enrichment in sh*Cdkn2a* tumors treated with anti-Pd1 vs. IgG compared to shControl tumors treated with anti-Pd1 vs. IgG, indicating their requirement for inducing ICB resistance in the context of low *Cdkn2a* expression (**Table S2**). As we observed differences in TME metabolites in *Cdkn2a*^Low^ tumors (**Fig. 3A**), we focused on plasma membrane solute carrier transporters (**Fig. 3C and Table S3**), which play critical roles in metabolite compartmentalization between the intracellular and extracellular space^120^. We cross-compared these genes with depleted analytes in sh*Cdkn2a* TIF (**Fig. 3A**). Using this comparison, we identified zinc and the plasma membrane zinc importer *Slc39a9* as potential drivers of anti-Pd1 resistance in sh*Cdkn2a* tumors (**Fig. 3A and Fig. 3C**). Notably, we found that the decrease of zinc in the TIF in sh*Cdkn2a* tumors correspond to increased zinc in cancer cells (**Fig. 3D**). Consistently, data mining of scRNA-Seq from melanoma patient specimens showed that *CDKN2A*^Low^ cancer cells have higher zinc scores compared to *CDKN2A*^High^ cancer cells (**Fig. 3E**). Altogether, these data demonstrate that loss of *CDKN2A* in cancer cells drives a differential zinc compartmentalization within the tumor, leading to zinc accumulation in cancer cells and concurrent depletion of zinc in the TME. Interestingly, a recent publication demonstrates that cancer patients with elevated serum zinc levels show improved overall survival when undergoing ICB treatment, including anti-PD1^95^. Consistently, we also found that anti-PD1-treated patients with low circulating zinc levels have reduced time-to-event outcomes compared to those with higher zinc levels (**Fig. 3F**), providing support for the role of zinc in ICB response. We validated that the knockdown of Slc39a9 (**Fig. S3B-D**) sensitizes *Cdkn2a*^Low^ tumors to anti-Pd1 treatment in our two models (**Fig. 3G-H**). Importantly, TAM depletion abrogated the anti-Pd1 efficacy following Slc39a9 knockdown (**Fig. 3I**), while there was no difference in anti-Pd1 response in tumors implanted into Rag-1-/- mice (**Fig. 3J**), suggesting that this restored response was dependent on TAMs and not T cells. Altogether, these data demonstrate that Slc39a9 is essential for anti-Pd1 resistance in *Cdkn2a*^Low^ tumors and that TAMs, not T cells, are the primary immune effectors driving anti-Pd1 responses in these tumors. Moreover, these data indicate that zinc compartmentalization between cancer cells and the TME plays a critical role in regulating the anti-Pd1 response.

**Figure 3.**
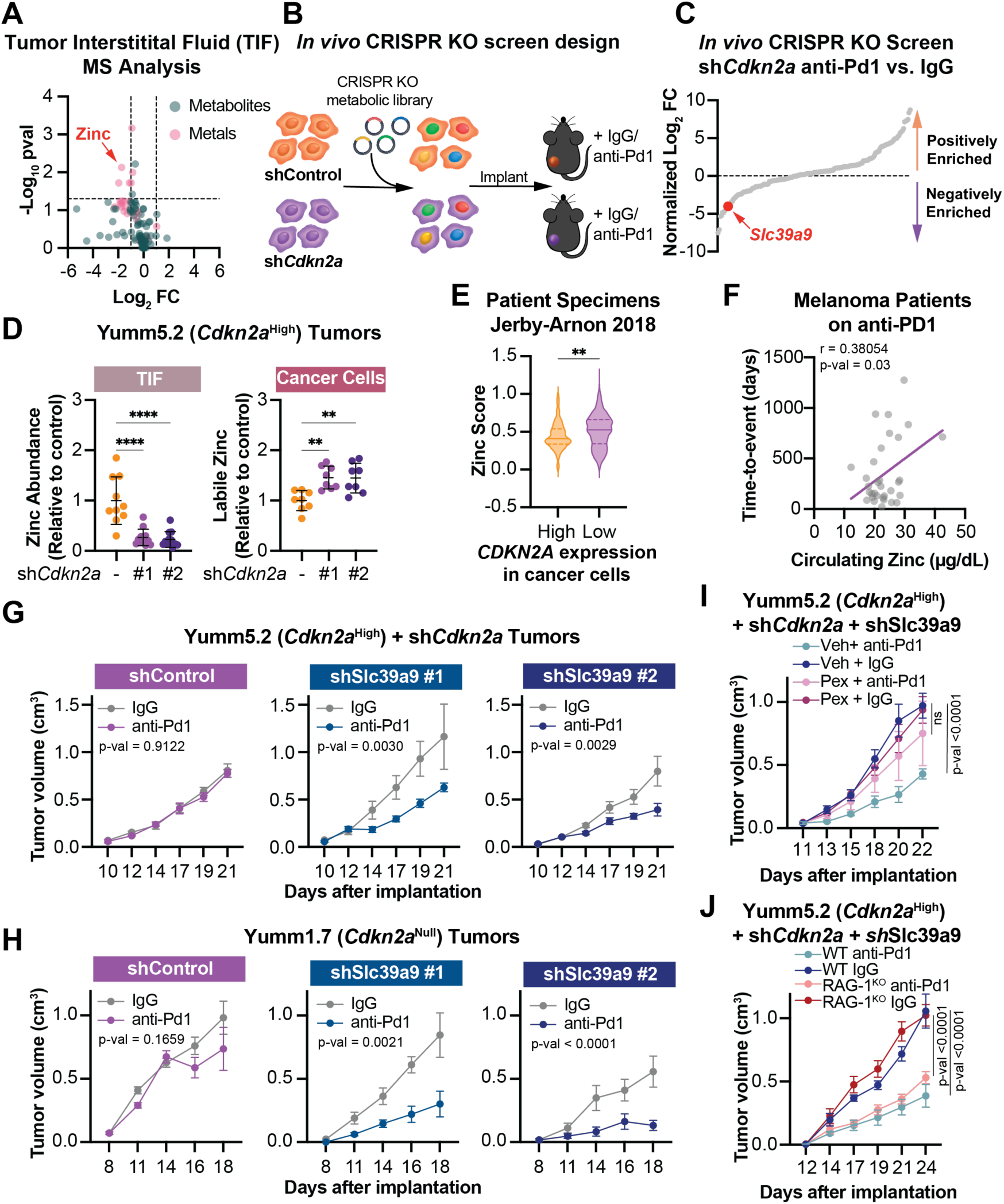
Zinc and Slc39a9 are key determinants of anti-Pd1 response in *Cdkn2a*^Low^ tumors. **(A)** Volcano plot of metabolites and metals in the tumor interstitial fluid (TIF) of tumors derived from Yumm5.2 (*Cdkn2a*^High^) cells with knockdown of GFP (shControl) or *Cdkn2a* (sh*Cdkn2a*). Metabolites were quantified by liquid chromatography–mass spectrometry (LC-MS) and metals by inductively coupled plasma mass spectrometry (ICP-MS). Data show log2 fold change (FC) of analytes in sh*Cdkn2a* vs. shControl and corresponding –log10 p-value (FDR-adjusted). **(B)** Schematic of *in vivo* metabolic CRISPR knockout (KO) screen design. Yumm5.2 (*Cdkn2a*^High^) cells with knockdown of GFP (shControl) or *Cdkn2a* (sh*Cdkn2a*) were implanted into mice. Mice were treated with IgG or anti-Pd1 (200 µg, i.p., 3×/week). **(C)** Ranked plot of CRISPR KO screen results in tumors (see Schematic in B). Graph is log2 Fold change (FC) of CRISPR negative scores of plasma membrane solute carrier transporters in sh*Cdkn2a* tumors treated with anti-Pd1 vs. IgG compared to shControl tumors treated with anti-Pd1 vs. IgG (see **Table S1**). **(D)** ICP-MS quantification of total zinc levels in TIF (left) and labile zinc levels in cancer cells (right) from tumors derived from Yumm5.2 (*Cdkn2a*^High^) cells with knockdown of GFP (shControl) or *Cdkn2a* (sh*Cdkn2a,* hairpins #1 and #2). Data show mean ± SD; one-way ANOVA. **(E)** Violin plot showing the cancer cell zinc score from scRNA-seq of melanoma patient tumors stratified by *CDKN2A* expression in cancer cells. Student’s t-test. **(F)** Correlation plot between total zinc levels in plasma (quantified by ICP-MS) and time-to-event outcomes in patients treated with anti-PD1. Pearson correlation. **(G)** Mice were implanted with Yumm5.2 (*Cdkn2a*^High^) cells with knockdown of *Cdkn2a* in combination with knockdown of GFP (shControl) or Slc39a9 (shSlc39a9, hairpins #1 and #2). Longitudinal tumor growth of mice treated with IgG or anti-Pd1 (200 µg, i.p., 3×/week). Mean ± SEM (n=5/group); mixed-effects model. **(H)** Mice were implanted with Yumm1.7 (*Cdkn2a*^Null^) cells with knockdown of GFP (shControl) or Slc39a9 (shSlc39a9, hairpins #1 and #2). Longitudinal tumor growth of mice treated with IgG or anti-Pd1 (200 µg, i.p., 3×/week). Mean ± SEM (n=5/group); mixed-effects model. **(I)** Mice were implanted with Yumm5.2 (*Cdkn2a*^High^) cells with knockdown of *Cdkn2a* in combination with knockdown of GFP (shControl) or Slc39a9 (shSlc39a9). Longitudinal tumor growth of mice treated with pexidartinib (Pex, 45 mg/kg, o.g., 3×/week) or vehicle, and IgG or anti-Pd1 (200 µg, i.p., 3×/week). Data show mean ± SEM (n=6/group); mixed-effects model. **(J)** Wildtype (WT) or B6.129S7-Rag1^tm1Mom^/J (RAG-1^KO^) mice were implanted with Yumm5.2 (*Cdkn2a*^High^) cells with knockdown of *Cdkn2a* in combination with knockdown of GFP (shControl) or Slc39a9 (shSlc39a9). Longitudinal tumor growth of mice treated with IgG or anti-Pd1 (200 µg, i.p., 3×/week). Mean ± SEM (n=5/group); mixed-effects model. * p<0.05, ** p<0.01, *** p<0.001, **** p<0.0001, ns = not significant.

### *CDKN2A*^Low^ cancer cells reprogram zinc compartmentalization via plasma membrane SLC39A9 in a mechanism mediated by cholesterol

Next, we sought to determine how *CDKN2A* regulates zinc homeostasis via SLC39A9. Prior work found that SLC39A9 is localized in the plasma membrane and imports zinc ions from the extracellular space into the cytosol^121–125^. We found that knockdown of *CDKN2A* in multiple human and mouse cancer cell models increased plasma membrane SLC39A9 (**Fig. 4A** and **Fig. S4A-B**), whereas expression of p16 in *Cdkn2a*^Low^ cancer cells decreased plasma membrane Slc39a9 (**Fig. 4B**). Cells with increased plasma membrane Slc39a9 had elevated intracellular zinc levels and reduced extracellular zinc levels, an effect that was reversed by knockdown of Slc39a9 in *Cdkn2a*^Low^ cancer cells (**Fig. 4C-E** and **Fig. S4C-F**). Importantly, knockdown of Slc39a9 decreased both labile zinc (**Fig. 4C-E**) and total zinc (**Fig. S4C-F**), where total zinc includes both labile zinc and zinc bound to proteins and other molecules. These findings indicate that Slc39a9 is a key regulator of zinc homeostasis, and that labile zinc quantification is a reliable indicator of zinc content in our model. Consistently, overexpression of wildtype Slc39a9 rescued the intracellular labile zinc defect in Slc39a9 knockdown cells, whereas the putative hypomorphic mutant Slc39a9^H156R 126–128^ was less effective in both Yumm5.2 (**Fig. 4F-H**) and Yumm1.7 (**Fig. S4G-I**) cells. Importantly, the decreased zinc uptake in cells expressing the Slc39a9^H156R^ mutant was not due to the inability of the mutant to localize to the plasma membrane (**Fig. 4F-G** and **Fig. S4G-H**). We did not observe a compensatory upregulation of other *Slc39* or *Slc30* genes known to transport zinc upon knockdown of Slc39a9 (**Fig. S4J** and **Table S4**), further confirming Slc39a9 as the key zinc transporter in these cells. Additionally, none of the metal ions commonly transported by SLC39 family members were altered upon knockdown of Slc39a9 (**Fig. S4K**), indicating that the observed changes are not due to differential uptake of other metals. These findings demonstrate that plasma membrane SLC39A9 is a major and specific regulator of zinc compartmentalization between intracellular and extracellular spaces in *CDKN2A*^Low^ cancer cells (**Fig. 4I**).

**Figure 4.**
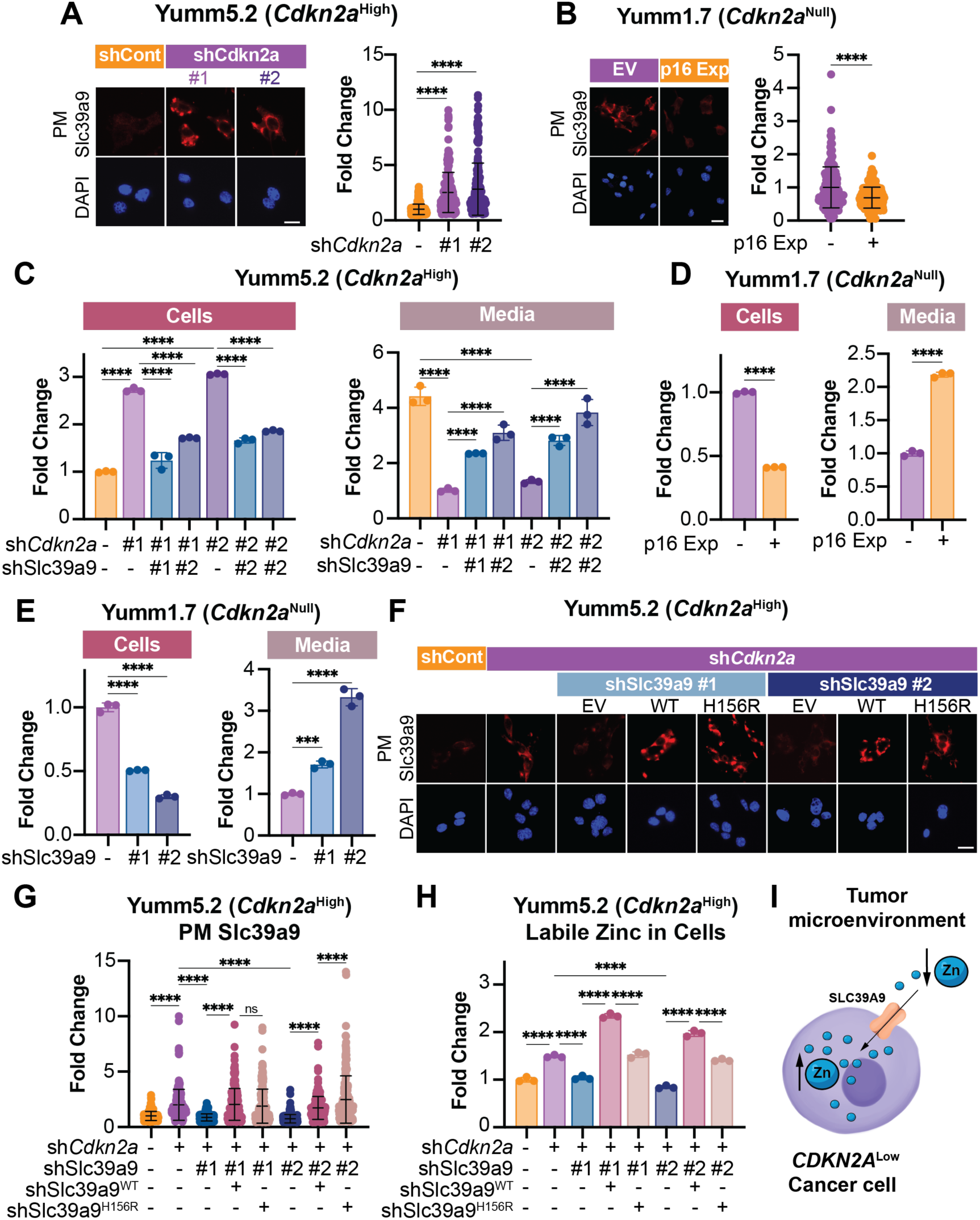
Increased plasma membrane Slc39a9 drives zinc uptake in *Cdkn2a*^Low^ cancer cells. **(A-B)** Representative images (left) and quantification (right) of Slc39a9 immunofluorescence on non-permeabilized cells to visualize plasma membrane (PM) Slc39a9 in **(A)** Yumm5.2 (*Cdkn2a*^High^) cells with knockdown of GFP (shControl) or *Cdkn2a* (sh*Cdkn2a,* hairpins #1 and #2) and **(B)** Yumm1.7 (*Cdkn2a*^Null^) cells expressing empty vector (EV) or physiological levels of p16 (p16 Exp). Mean ± SD (n=200/group); one-way ANOVA. **(C-E)** Bar graph showing labile zinc quantification in cells (left) and media (right) in **(C)** Yumm5.2 (*Cdkn2a*^High^) cells with knockdown of GFP (shControl) or *Cdkn2a* (sh*Cdkn2a,* hairpins #1 and #2) in combination with knockdown of GFP (shControl) or Slc39a9 (shSlc39a9, hairpins #1 and #2), **(D)** Yumm1.7 (*Cdkn2a*^Null^) cells expressing empty vector (EV) or physiological levels of p16 (p16 Exp), and **(E)** Yumm1.7 (*Cdkn2a*^Null^) cells with knockdown of GFP (shControl) or Slc39a9 (shSlc39a9, hairpins #1 and #2). Mean ± SD; Student’s t-test and one-way ANOVA. **(F-H)** Yumm5.2 (*Cdkn2a*^High^) cells with knockdown of GFP (shControl) or *Cdkn2a* (sh*Cdkn2a,* hairpins #1 and #2) alone or in combination with empty vector (EV) control, wildtype (WT), or hypomorphic mutant (H156R) Slc39a9. **(F)** Representative images of Slc39a9 immunofluorescence on non-permeabilized cells to visualize plasma membrane (PM) Slc39a9. **(G)** Quantification of (F). Mean ± SD (n=200/group); one-way ANOVA. **(H)** Labile zinc quantification in cells. Mean ± SD; one-way ANOVA. **(I)** Schematic of how plasma membrane Slc39a9 compartmentalizes zinc between the cells and the tumor microenvironment. Scale bars = 25μm.* p<0.05, ** p<0.01, *** p<0.001, **** p<0.0001, ns = not significant.

Finally, we sought to determine the mechanism whereby *Cdkn2a*^Low^ cancer cells have increased plasma membrane Slc39a9. We did not observe differential mRNA expression of *Slc39a9* upon *Cdkn2a* knockdown or p16 expression (**Fig. S5A**). SLC39A9 is a transmembrane protein, which are often anchored to the plasma membrane through cholesterol^129^. Our prior study found that human cells with knockdown of *CDKN2A*/p16 have increased cholesterol^56^, a finding we validated in the complementary mouse cell line models used in this study (**Fig. S5B-C**). Thus, we hypothesized that increased cholesterol levels in *CDKN2A*^Low^ cancer cells may enhance plasma membrane SLC39A9 accumulation. Consistent with this hypothesis, depleting plasma membrane cholesterol with methyl-β-cyclodextrin (MβCD)^130,131^ or inhibiting *de novo* cholesterol synthesis using statins reduced plasma membrane Slc39a9 levels and decreased intracellular labile zinc in *Cdkn2a*^Low^ cancer cells (**Fig. 5A-I**). Altogether, these data demonstrate that *Cdkn2a*^Low^ cancer cells increase intracellular zinc via increased cholesterol-mediated plasma membrane Slc39a9.

**Figure 5.**
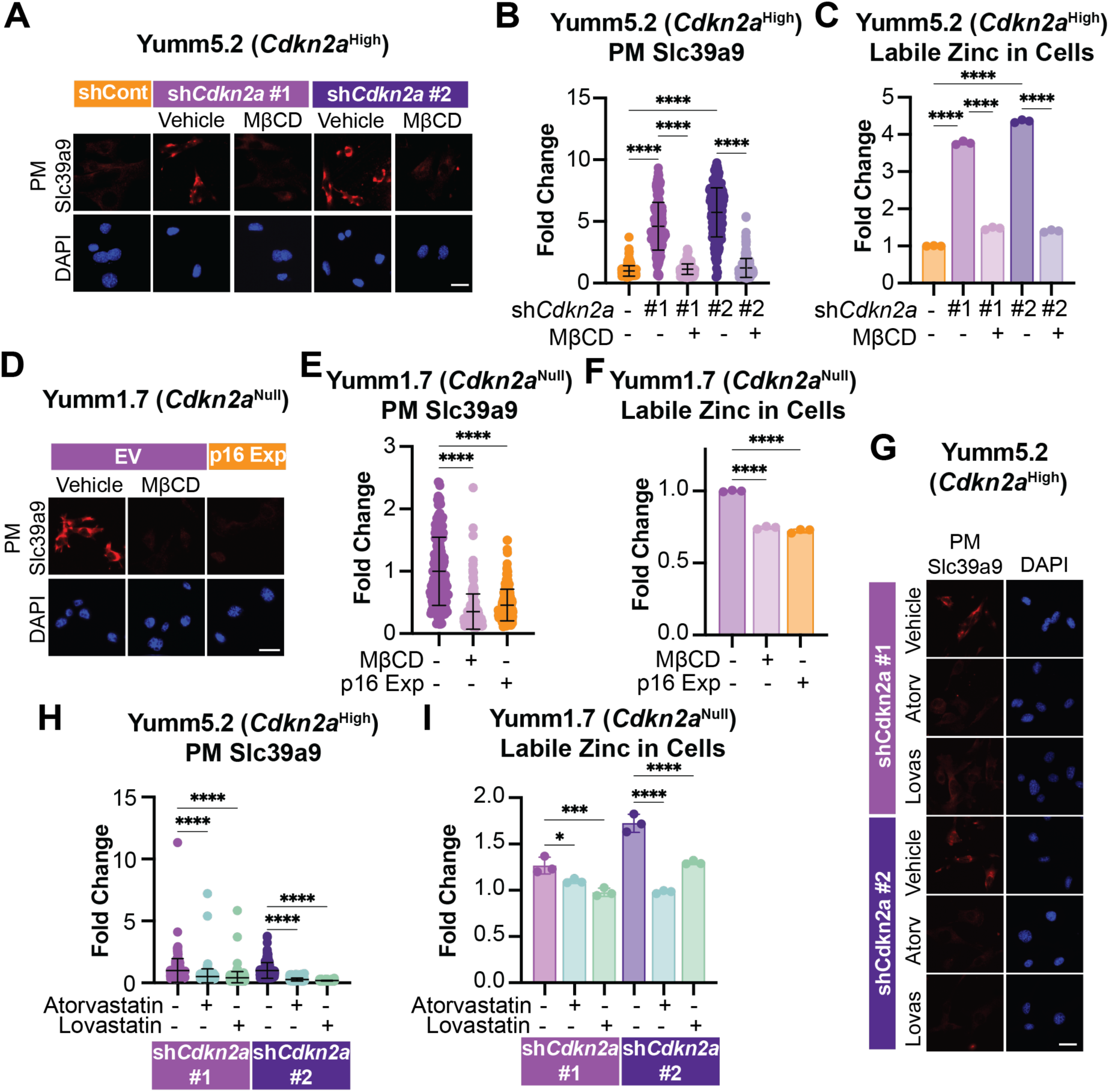
Increased plasma membrane Slc39a9 in *Cdkn2a*^Low^ cancer cells is mediated by cholesterol. **(A-C)** Yumm5.2 (*Cdkn2a*^High^) cells with knockdown of GFP (shControl) or *Cdkn2a* (sh*Cdkn2a,* hairpins #1 and #2) were treated with vehicle or Methyl-β-cyclodextrin (MβCD; 10mM, 1h). **(A)** Representative images of Slc39a9 immunofluorescence on non-permeabilized cells to visualize plasma membrane (PM) Slc39a9. **(B)** Quantification of (A). Mean ± SD (n=200/group); one-way ANOVA. **(C)** Labile zinc quantification in cells. Mean ± SD; one-way ANOVA. **(D-F)** Yumm1.7 (*Cdkn2a*^Null^) cells expressing empty vector (EV) or physiological levels of p16 (p16 Exp) were treated with vehicle or Methyl-β-cyclodextrin (MβCD; 10mM, 1h). **(D)** Representative images of Slc39a9 immunofluorescence on non-permeabilized cells to visualize plasma membrane (PM) Slc39a9. **(E)** Quantification of (D). Mean ± SD (n=200/group); one-way ANOVA. **(F)** Labile zinc quantification in cells. Mean ± SD; one-way ANOVA. **(G-I)** Yumm5.2 (*Cdkn2a*^High^) cells with knockdown of *Cdkn2a* (sh*Cdkn2a,* hairpins #1 and #2) were treated with vehicle or the statins atorvastatin (Atov, 1µM, 4 days) or lovastatin (Lovas,1µM, 4 days). **(G)** Representative images of Slc39a9 immunofluorescence on non-permeabilized cells to visualize plasma membrane (PM) Slc39a9, **(H)** Quantification of (G). Mean ± SD (n=200/group); one-way ANOVA. **(I)** Labile zinc quantification in cells. Mean ± SD; one-way ANOVA. Scale bars = 25μm.* p<0.05, ** p<0.01, *** p<0.001, **** p<0.0001, ns = not significant.

### Competition for zinc between *Cdkn2a*^Low^ cancer cells and TAMs regulates phagocytosis and anti-Pd1 response

We found that *CDKN2A*^Low^ tumors do not respond to anti-PD1 treatment (**Fig. 1A–C**), and that macrophages are less phagocytic when exposed to *Cdkn2a*^Low^ cancer cells, both *in vitro* (**Fig. 2H**) and *in viv*o (**Fig. 2C–D** and **F-G**). We also found that *Cdkn2a*^Low^ cancer cells deplete microenvironmental zinc through zinc uptake via increased Slc39a9 at the plasma membrane (**Fig. 4C-E and Fig. S4C-F**) and that knockdown of Slc39a9 in *Cdkn2a*^Low^ cells restores anti-Pd1 response (**Fig. 3G–H**). Thus, we hypothesized that the availability of zinc in the TME may play a direct role in TAM phagocytosis. Analysis of publicly available scRNA-seq data from melanoma and HNSCC patients showed that TAMs in *CDKN2A*^Low^ tumors have lower zinc scores and that zinc scores positively correlate with phagocytosis scores (**Fig. 6A** and **Fig. S6A**). Moreover, further analysis of our scRNA-Seq data from Yumm5.2 tumors demonstrated that TAMs in *Cdkn2a*^Low^ tumors, which have low phagocytic activity (**Fig. 2C**), also have decreased zinc scores (**Fig. S6B**). Consistently, *in vivo* validation demonstrated that TAMs in tumors derived from *Cdkn2a*^Low^ cancer cells have reduced zinc levels (**Fig. 6B** and **Fig. S6C**), and zinc levels positively correlate with phagocytosis (**Fig. 6B** and **Fig. S6D**). Moreover, we found that RAW264.7 macrophages with higher intracellular zinc have greater phagocytic activity *in vitro* (**Fig. S6E**). Strikingly, while anti-Pd1 treatment increased the expression of phagocytosis-related genes in RAW264.7 macrophages *in vitro*, this effect was further amplified by zinc supplementation and suppressed by zinc chelation (**Fig. S6F**). These data demonstrate that zinc availability indeed modulates anti-Pd1-mediated macrophage phagocytosis.

**Figure 6.**
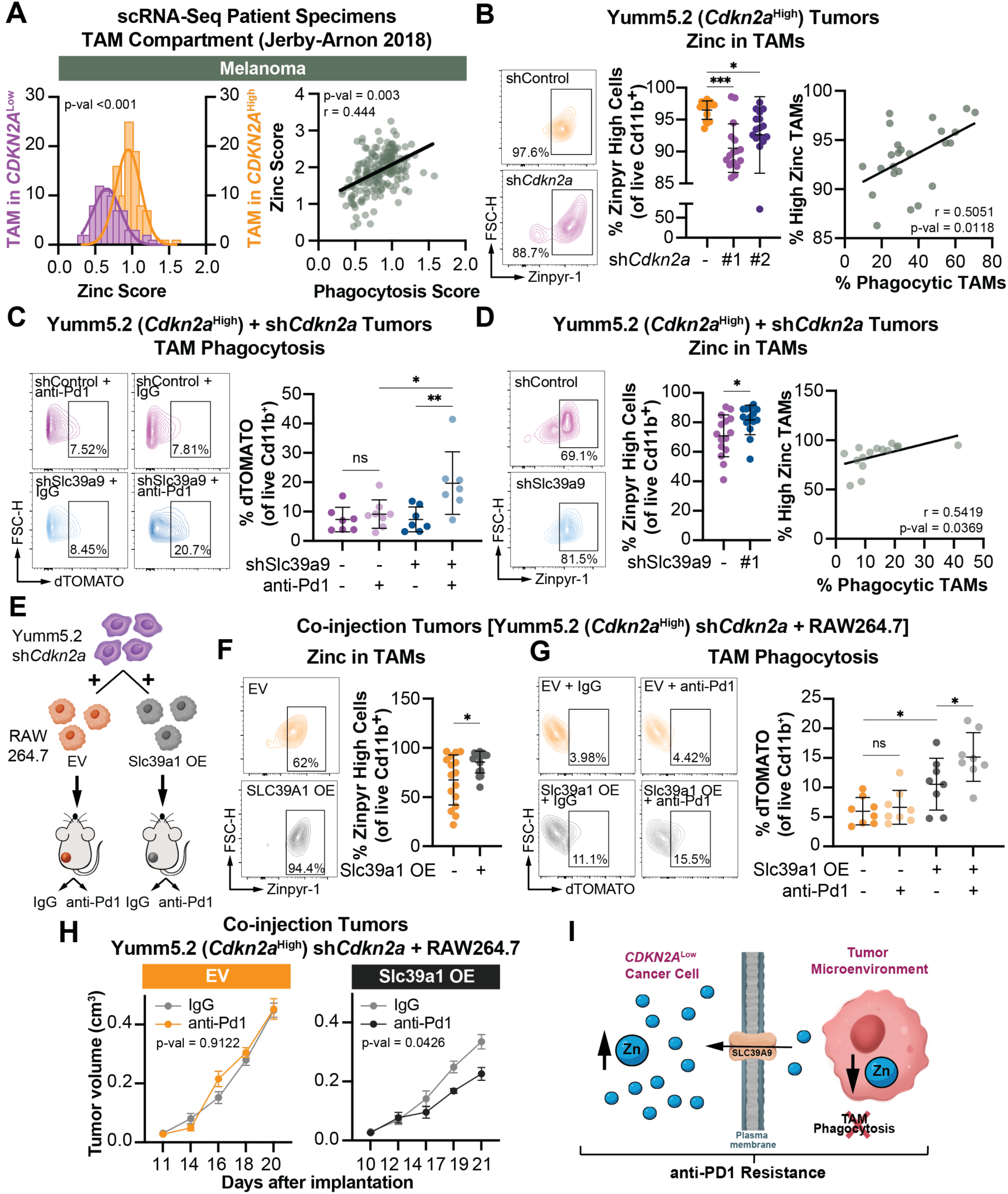
Reduced zinc availability to TAMs in *Cdkn2a*^Low^ tumors suppresses anti-Pd1-driven phagocytosis. **(A)** scRNA-Seq dataset from melanoma patients stratified by *CDKN2A* expression in cancer cells. Histogram showing the zinc score in TAMs from the indicated scRNA-Seq datasets from melanoma patients stratified by *CDKN2A* expression in cancer cells (left) and zinc score correlation with phagocytic score (right). Linear mixed models and Pearson correlation. **(B)** Mice were implanted with dTOMATO-labeled Yumm5.2 (*Cdkn2a*^High^) cells with knockdown of GFP (shControl) or *Cdkn2a* (sh*Cdkn2a,* hairpins #1 and #2). Representative labile zinc flow cytometry plots in TAMs (left), quantification of labile zinc in TAMs (center), and correlation between labile zinc and phagocytosis in TAMs (right). Mean ± SD; one-way ANOVA and Pearson correlation. **(C-D)** Mice were implanted with dTOMATO-labeled Yumm5.2 (*Cdkn2a*^High^) cells with knockdown of *Cdkn2a* (sh*Cdkn2a*) alone or in combination with knockdown of GFP (shControl) or Sl39a9 (shSlc39a9). **(C)** Representative flow cytometry plots of TAM phagocytosis (left) and quantification of TAM phagocytosis (right). Mean ± SD; one-way ANOVA. **(D)** Representative labile zinc flow cytometry plots in TAMs (left), quantification of labile zinc in TAMs (center), and correlation between labile zinc and phagocytosis in TAMs (right). Mean ± SD; one-way ANOVA and Pearson correlation. **(E)** Schematic of macrophage coinjection model. **(F-H)** RAW264.7 macrophages expressing empty vector (EV) control or Slc39a1 (Slc39a1 OE) were co-injected in immunocompromised NSG mice with Yumm5.2 (*Cdkn2a*^High^) cells with knockdown of *Cdkn2a* (shCdkn2a) at a ratio of 1:10. Mice were treated with IgG or anti-Pd1 (200 µg, i.p., 3×/week). **(F)** Representative flow cytometry plots (left) and quantification (right) of labile zinc in TAMs. Mean ± SD; Student’s t-test. **(G)** Representative flow cytometry plots (left) and quantification (right) of the percentage of TAMs with dTOMATO signal (phagocytic TAMs). Mean ± SD; one-way ANOVA. **(H)** Longitudinal tumor growth. Mean ± SEM (n=8/group); mixed-effects model. **(I)** Schematic of zinc competition between *CDKN2A*^Low^ cells and TAMs, leading to decreased phagocytosis and anti-PD1 resistance. * p<0.05, ** p<0.01, *** p<0.001, **** p<0.0001, ns = not significant.

We found that knockout or knockdown of the zinc importer Slc39a9 restores anti-Pd1 response in *Cdkn2a*^Low^ tumors (**Fig. 3C, 3G-H**) and that increased intracellular zinc in macrophages correlates with enhanced phagocytosis in human melanoma and HNSCC patients (**Fig. 6A** and **Fig. S6A**). A recent publication found that increased metal signatures in TAMs, mainly genes related to zinc function, correlate with improved ICB responses in patients^30^. Additionally, both published data^95^ and our data (**Fig. 3F**) show that patients with higher circulating zinc levels show improved outcomes on anti-PD1 therapy. Thus, we aimed to determine whether zinc depletion in the TME, driven by Slc39a9 in *Cdkn2a*^Low^ cancer cells, impairs TAM zinc content and phagocytosis. To test this, we knocked down Slc39a9 in fluorophore-labeled Yumm5.2 *Cdkn2a*^Low^ cancer cells and implanted them into mice (**Fig. 2E**). Slc39a9 knockdown restored TAM phagocytosis upon anti-Pd1 (**Fig. 6C**), corresponding with increased zinc in TAMs (**Fig. 6D**). Altogether, these data indicate that Slc39a9-mediated zinc sequestration by cancer cells directly impairs TAM function in response to anti-Pd1. We did not observe consistent expression differences in “eat me” or “don’t eat me” signals in *Cdkn2a*^Low^ cells with or without Slc39a9 knockdown (**Fig. S6G** and **Table S5**), nor did we detect changes in Pd-l1 expression on cancer cells (**Fig. S6H-I**), indicating that the observed changes in TAM phagocytosis and anti-Pd1 response is macrophage-intrinsic. Finally, to demonstrate that increased zinc in macrophages directly increases phagocytosis *in vivo*, we engineered RAW264.7 macrophages that have the ability to compete for TME zinc by overexpressing the zinc importer Slc39a1 (**Fig. S6J**). Indeed, these macrophages are able to increase intracellular zinc *in vitro* (**Fig. S6K**). We then co-implanted these zinc-replete macrophages with Yumm5.2 sh*Cdkn2a* cells in immunodeficient NSG mice (**Fig. 6E**). TAMs in tumors co-implanted with Slc39a1-overexpressing macrophages had higher intracellular zinc (**Fig. 6F**). Excitingly, upon anti-Pd1 treatment, these TAMs had increased phagocytic activity (**Fig. 6G**), which was associated with reduced tumor volume (**Fig. 6H**). Together, our data demonstrate that zinc is critical for TAM phagocytosis and that zinc-replete TAMs respond to anti-Pd1 by increasing their phagocytic gene expression and activity. Thus, increased zinc uptake by *Cdkn2a*^Low^ cancer cells impairs TAM phagocytosis by outcompeting macrophages for zinc, corresponding to anti-Pd1 resistance (**Fig. 6I**).

### High zinc diet combined with anti-Pd1 decreases tumor burden and enhances TAM phagocytosis in *Cdkn2a*^Low^ tumors

Finally, we aimed to find a manner to exploit our discoveries to improve anti-Pd1 treatment in *Cdkn2a*^Low^ tumors. We found that *Cdkn2a*^Low^ cancer cells deplete the TME of zinc by upregulating plasma membrane Slc39a9 (**Fig. 4A-E** and **Fig. S4C-F**), therefore out-competing TAMs for zinc (**Fig. 6B-D**). We also found that engineering TAMs that have an enhanced ability to compete for zinc uptake have increased phagocytosis (**Fig. 6E-H**). Since there are no available inhibitors for SLC39A9, we hypothesized that increasing dietary zinc would raise zinc levels in the TME to restore zinc within TAMs and promote an anti-Pd1 response. Excitingly, a high zinc diet (∼3X higher than normal chow) significantly elevated TME zinc levels (**Fig. 7A-B**), which corresponded to decreased tumor volume in *Cdkn2a*^Low^ tumors upon anti-Pd1 treatment (**Fig. 7C-D and Fig. S7A**). The dietary intervention did not significantly affect food intake or body weight (**Fig. S7B-C**) nor did it markedly alter tumor growth in the IgG control treatment group (**Fig. S7D**). Similar to *Cdkn2a*^Low^ tumors with Slc39a9 knockdown (**Fig. 3I**), TAM depletion abrogated this anti-Pd1 response (**Fig. 7E**), indicating that TAMs are essential for the anti-Pd1 response of *Cdkn2a*^Low^ cancer cells on a high zinc diet. Excitingly, we confirmed that the high zinc diet increased TAM zinc levels (**Fig. 7F** and **Fig. S7E**), and these zinc-replete TAMs had enhanced phagocytic activity following anti-Pd1 treatment (**Fig. 7G** and **Fig. S7F**). Similar to our findings with knockdown of Slc39a9 (**Fig. S6I**), the observed increase in TAM phagocytosis was independent of changes in Pd-l1 expression on cancer cells (**Fig. S7G**), further confirming that TAM phagocytosis and decreased tumor burden upon anti-Pd1 treatment is macrophage-intrinsic. Taken together, these findings support the notion that replenishing the TME with zinc via a dietary intervention can reverse anti-Pd1 resistance in *Cdkn2a*^Low^ tumors, which is associated with an enhanced TAM phagocytotic response (**Fig. 7H**).

**Figure 7.**
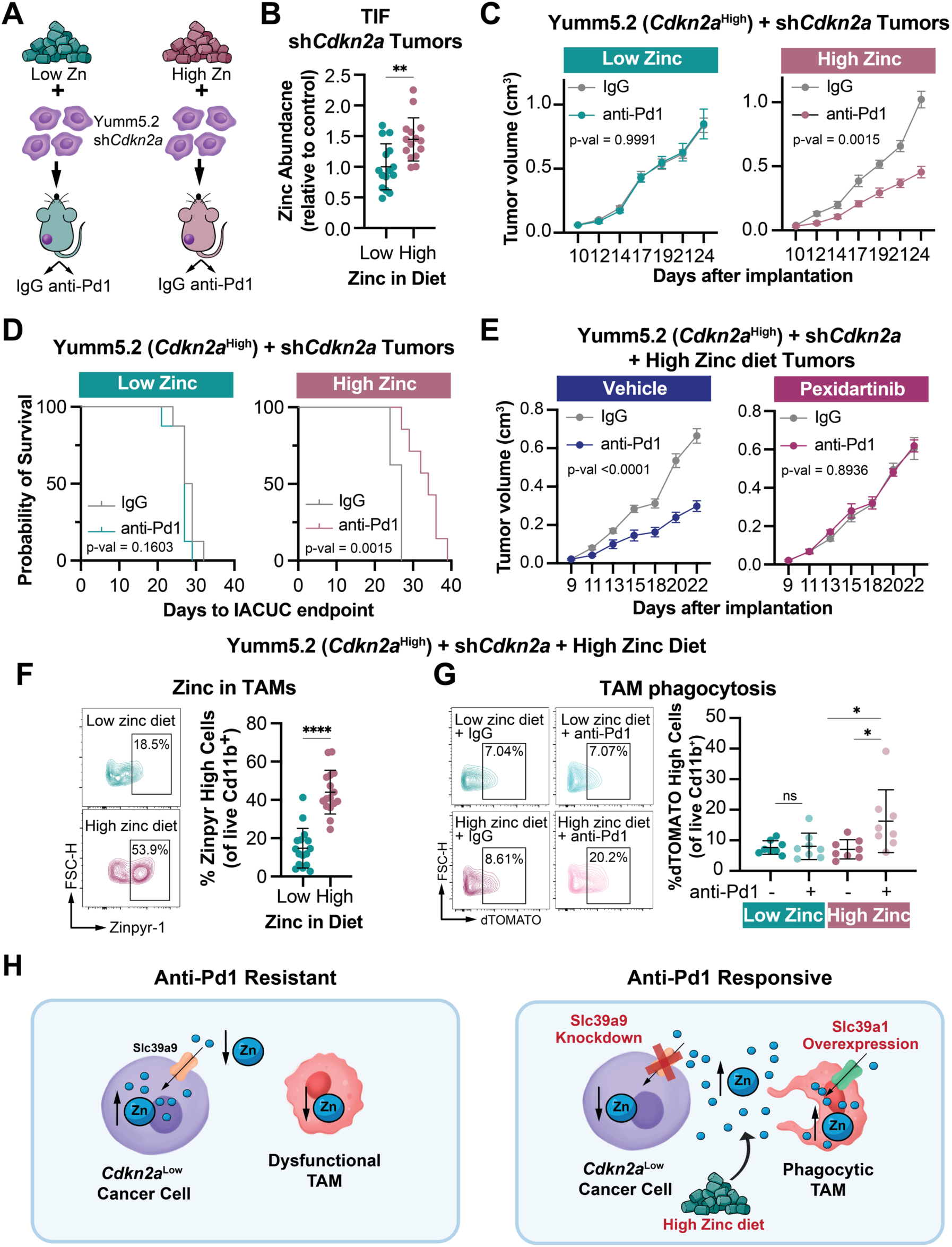
Replenishing TME zinc via the diet in *Cdkn2a*^Low^ tumors re-educates TAMs and promotes anti-Pd1-mediated phagocytosis. **(A)** Schematic of diet study. **(B-D)** Mice were assigned to low (55 mg/kg) or high (300 mg/kg) zinc diets, implanted with Yumm5.2 (*Cdkn2a*^High^) cells with knockdown of *Cdkn2a* (sh*Cdkn2a*), and treated with IgG or anti-Pd1 (200 µg, i.p., 3×/week). **(B)** Dot plot showing total levels of zinc in the tumor interstitial fluid (TIF) quantified by ICP-MS. Mean ± SD; student’s t-test. **(C)** Longitudinal tumor growth. Mean ± SEM (n=8/group); mixed-effects model. **(D)** Kaplan-Meier plot depicting survival analysis. Log-rank (Mantel-Cox) test. **(E)** Mice fed a high (300 mg/kg) zinc diet were implanted with Yumm5.2 (*Cdkn2a*^High^) cells with knockdown of *Cdkn2a* (sh*Cdkn2a*). Mice were treated with Pexidartinib (45 mg/kg, o.g., 3×/week) or vehicle in combination with IgG or anti-Pd1 (200 µg, i.p., 3×/week). Longitudinal tumor growth. Mean ± SEM (n=8/group); mixed-effects model. **(F-G)** Mice were assigned to a high (300 mg/kg) zinc diet, implanted with Yumm5.2 (*Cdkn2a*^High^) cells with knockdown of *Cdkn2a* (shCdkn2a), and treated with IgG or anti-Pd1 (200 µg, i.p., 3×/week). **(F)** Representative flow cytometry plots (left) and quantification (right) of labile zinc in TAMs. Mean ± SD; student’s t-test. **(G)** Representative flow cytometry plots (left) and quantification (right) of the percentage of TAMs with dTOMATO signal (phagocytic TAMs). Mean ± SD; one-way ANOVA. (H) *Cdkn2a*^Low^ cancer cells sequester TME zinc via increased plasma membrane Slc39a9, leading to macrophage dysfunction and anti-Pd1 resistance. Replenishment of TME zinc via knockdown of Slc39a9 in cancer cells, overexpression of Slc39a1 in co-implanted macrophages, or high zinc diet increases intracellular zinc in macrophages priming their phagocytic activity to respond to anti-Pd1. * p<0.05, ** p<0.01, *** p<0.001, **** p<0.0001, ns = not significant.

## DISCUSSION

The role of *CDKN2A* as a critical regulator of the cell cycle has been well established for >30 years^51,132^. However, accumulating evidence from our lab and others has revealed that *CDKN2A*/p16 loss has non-canonical pro-tumorigenic functions that extend beyond its role in cell cycle control, including in metabolism^133^. Moreover, there is recent evidence that nutrient competition within the TME can affect immune cell activity and function^134–137^, although how the micronutrient zinc is compartmentalized within cancer and immune cells has never been explored. Here, we demonstrate that loss of *CDKN2A* in cancer cells enhances zinc uptake through upregulation of the SLC39A9 transporter at the plasma membrane, a process that is mediated by cholesterol. The increased zinc uptake by *Cdkn2a*^Low^ cancer cells leads to a zinc-depleted TME, impairing TAM phagocytic activity and diminishing the efficacy of anti-Pd1 treatment. Our findings underscore the critical role of *CDKN2A*/p16 expression in modulating the immune response through zinc, demonstrate an anti-tumorigenic role for zinc-replete TAMs, and open up new therapeutic possibilities through dietary interventions. Moreover, given that the effects we observed were T cell-independent, and that many human tumors are myeloid-rich and T cell-poor^106^, our data provide compelling evidence that elevating TME zinc could sensitize tumors that have generally been considered anti-PD1 refractory.

Our studies and others have shown that suppression of *CDKN2A*/p16 correlates with reduced expression of pro-inflammatory factors across various diseases, including cancer^18,63,138–142^. Consistently, *CDKN2A*^Low^ tumors exhibit ‘cold’ immune phenotypes with reduced immune infiltration, which has been linked to the observed resistance to ICB^1–5,11,143–146^. Excitingly, our findings reveal that TAMs, rather than T cells, are the critical immune cell promoting the response to anti-PD1 in *CDKN2A*^Low^ tumors (**Fig. 3I-J**). In this regard, a small but growing body of literature suggests that patients with T cell-poor tumors can still benefit from ICB, potentially through effects on other immune cell populations^25–27^. Consistent with this, our analysis shows that around 20% of patients with myeloid-rich, T cell-poor tumors exhibit clinical benefit from anti-PD1 therapy beyond 20 months, and >7% continue to benefit beyond 30 months (**Fig. 1G**). Recent studies have shown that certain myeloid cells, including TAMs, can directly respond to anti-PD1 and contribute to therapeutic efficacy^28–35^. Using myeloid-rich, T cell-poor models, we found that *Cdkn2a*^Low^ murine tumors have increased TAMs (**Fig. 3A**), consistent with prior reports in human patients^1,5^. However, while other research focused on the macrophage polarization status^1,3,6^, here we found that *CDKN2A*^Low^ cancer cells modulate TAM phagocytic activity (**Fig. 2F-H**). This paradigm shift underscores the critical importance of studying TAM activity, not simply polarization status or macrophage markers, in the context of ICB treatment to improve anti-tumor efficacy. Our results also give rise to the intriguing possibility that myeloid-rich, T cell-poor tumors, which have typically been considered refractory to ICB, can be sensitized by reprogramming TAMs to a more phagocytic phenotype.

Previous research indicates that TAMs can directly respond to ICB in part by enhancing phagocytosis^28,29,35^. Additionally, a recent pan-cancer study in patient samples identified a subgroup of TAMs that upregulate zinc-binding metallothioneins (named *metalloMac*) that are enriched in ICB-responsive patients^30^. This is consistent with our data demonstrating that elevated zinc in TAMs boosts the response to anti-Pd1 (**Fig. 6G-H**). Indeed, zinc has been previously shown to enhance phagocytosis capacity and/or pathogen clearance^68–70,147–150^, although none of this work is in cancer and the mechanism is not well understood. Here we found that there is a correlation between the amount of intracellular zinc in TAMs and their phagocytic activity in both *in vivo* tumor models and human tumors (**Fig. 6A-B**, **6D** and **Fig. S6A, S6D**). Moreover, zinc supplementation enhanced the expression of phagocytosis-related genes in macrophages treated with anti-Pd1 *in vitro*, an effect that was abolished by zinc chelation (**Fig. S6F**). The human genome encodes over 700 zinc-finger proteins, the largest class of putative transcription factors^151^, and prior research has explored whether intracellular zinc levels directly influence gene transcription^152^. Future studies will investigate the specific transcription factors that drive macrophage phagocytic activity in response to zinc replenishment. Collectively, our findings underscore the critical role of TAM zinc metabolism in modulating the efficacy of anti-PD1 therapy, particularly in the context of *CDKN2A*^Low^ tumors where zinc homeostasis within the TME is disrupted. This study also highlights the potential for boosting zinc metabolism in TAMs as a strategy to enhance the therapeutic response to anti-PD1 in cancer patients, regardless of T cell infiltration.

Our study demonstrates that *CDKN2A*/p16 itself drives inherent resistance to anti-PD1. There are a handful of prior reports linking loss of *CDKN2A* expression, or 9p21 deletion, with changes in anti-tumor immunity and ICB response^1–18^. Here, we found that *Cdk2na*, and more specifically p16 alone, can drive differential anti-PD1 responses (**Fig. 1B-C**). Consistent with our findings, an integrated analysis of immunogenomic and clinical data from over 1000 patients across 8 solid tumors demonstrated that 9p21 deletions, specifically encompassing *CDKN2A* and *MTAP*, were associated with decreased tumor immune responses and resistance to anti-PD1 or anti-PD-L1 monotherapy^1^. In contrast, a prior study using cells with different 9p21 deletions showed deletions involving only the *CDKN2A* gene did not affect the response to anti-CTLA4^104^. This discrepancy may arise from differences in the models and immunotherapies used. Despite these insights, routine assessment of *CDKN2A*/p16 expression remains uncommon in clinical practice, except in specific cases such as HPV-positive tumors or when germline alterations are suspected, particularly in individuals with a family history of melanoma and/or pancreatic cancer^153–155^. Together with the prior research, our work provides compelling evidence for the need to routinely assess *CDKN2A* status in cancer diagnostics, particularly as our research suggests that treatment outcomes of patients harboring *CDKN2A*^Low^ tumors may be improved by combining ICB therapy with dietary zinc supplementation.

*CDKN2A* has non-canonical roles in intracellular metabolism^54,55,58,59,117–119^, and we emphasize here a role for *CKDN2A* in influencing metabolites within the TME. Specifically, we found that *CDKN2A*^Low^ cancer cells sequester zinc from the TME (**Fig. 4**), reducing macrophage phagocytic activity during anti-Pd1 treatment (**Fig. 6C-D** and **Fig. S6C-D**), thereby diminishing the therapeutic response (**Fig. 3G-H**). Zinc is an essential micronutrient critical for the proper functioning of cells. For example, zinc deficiency impairs immune cell activity, while zinc supplementation can either restore normal function in deficient cells or enhance immune responses in healthy cells^156^. Despite decades of research on zinc’s role in cancer, its biological effects remain poorly understood and are often contradictory^157^. These inconsistencies stem in part from a historical focus on systemic zinc levels, which overlook the intricate regulation and compartmentalized distribution of zinc within the TME. In this study, we found that increased intracellular zinc in *CDKN2A*^Low^ cancer cells was mediated by a significant upregulation of the zinc transporter SLC39A9 at the plasma membrane (**Fig. 4**). Upregulation of plasma membrane Slc39a9 was regulated by *de novo* cholesterol synthesis (**Fig. 5**), which we have previously found to be elevated in sh*CDKN2A*/shp16 cells^117^ (**Fig. S5B-C**). Interestingly, several epidemiological studies have shown a strong correlation between statin use—drugs that lower cholesterol—and improved response to ICB in cancer patients^158–163^. Our data suggest that this effect may be mediated, at least in part, by reduced plasma membrane Slc39a9 expression in cancer cells (**Fig. 4-5**), thereby increasing zinc availability for immune cells. Future studies will be needed to determine the extent to which the ICB benefit observed in statin-treated patients is driven by altered zinc compartmentalization via SLC39A9. Moreover, these results contribute to the growing body of research highlighting the critical role of metabolite compartmentalization within the TME^134,136,164,165^ and underscores the limitations of systemic zinc-level assessments in evaluating zinc’s role in cancer.

Zinc can be modulated by the diet, and prior work has associated high dietary zinc intake with improved cancer outcomes or decreased risk^166–171^. However, others report adverse^172–175^ or neutral effects^176–179^. Here, we found that high dietary zinc effectively increases the zinc content in the TME and improves ICB response of *Cdkn2a*^Low^ tumors (**Fig. 7**). This increase in zinc in the diet is ∼3X more than the normal chow. The recommended daily allowance (RDA) of zinc is 8 mg for adult women and 11 mg for adult men^180^. Assuming similar zinc metabolism between humans and mice, this increase in dietary zinc is equivalent to a 30 mg zinc supplement, which is available over-the-counter. Future studies are required to explore the potential benefits of combining ICB with this highly accessible and cost-effective zinc supplementation for patients with *CDKN2A*^Low^ tumors. In contrast to our study, a recent article found that high dietary zinc promoted tumor progression in a B16F10 melanoma model by increasing regulatory T cells^98^. These differences are likely due to the amount of zinc in the diet (300mg/kg in our study vs. 80mg/kg in^98^), delivery method, or other context-dependent differences. Indeed, 80 mg/kg is below our normal chow levels, suggesting that this study did not achieve high enough intratumoral zinc levels to restore zinc in TAMs and enhance their phagocytosis. Nonetheless, these discrepancies underscore the importance of considering tumor-specific factors, zinc delivery methods, and bioavailability when exploring zinc-based therapies. In this regard, our data show that the high zinc diet used in this study effectively increases zinc levels not only in the TME (**Fig. 7B**), but also in intratumoral macrophages (**Fig. 7F**). Although we found that replenishing the TME with zinc enhances TAM zinc content and phagocytic activity in C*dkn2a*^Low^ tumors, the role of increased intracellular zinc in *CDKN2A*^Low^ cancer cells remains unclear. The literature reports both pro-tumorigenic^181–183^ and cytotoxic^184–187^ effects of high intracellular zinc in cancer cells. Thus, it is interesting to speculate that zinc may have pleiotropic roles within cancer cells similar to that observed for ROS^188^, where excessively high levels are detrimental, but moderate to high levels promote tumorigenesis. Further research is needed to determine the intracellular mechanisms of zinc in C*DKN2A*^Low^ cancer cells.

In conclusion, the TME represents a competitive environment where access to nutrients, such as zinc, critically shapes the dynamics between tumor and immune cells. Our findings demonstrate that suppression of *Cdkn2a* confers the ability to deplete zinc from the TME by upregulating the Slc39a9 importer, impairing TAM function, and reducing the efficacy of anti-Pd1 therapy irrespective of T cells. This study highlights a previously unrecognized role of *CDKN2A* in maintaining a balanced TME metabolism that supports immune activity and therapeutic responses. Our results also emphasize that, in certain contexts, T cells are dispensable for an anti-PD1 response. In summary, our results identify an unexpected role for *CDKN2A* as a master regulator of zinc homeostasis, which impacts the TME and anti-PD1 therapy through a T cell-independent mechanism. These results have major implications not only for understanding novel roles for *CDKN2A* outside of the cell cycle, but also in underscoring the anti-tumorigenic potential of TAMs that can be harnessed for improved ICB response.

## SUPPLEMENTAL FIGURES

**Figure S1.**
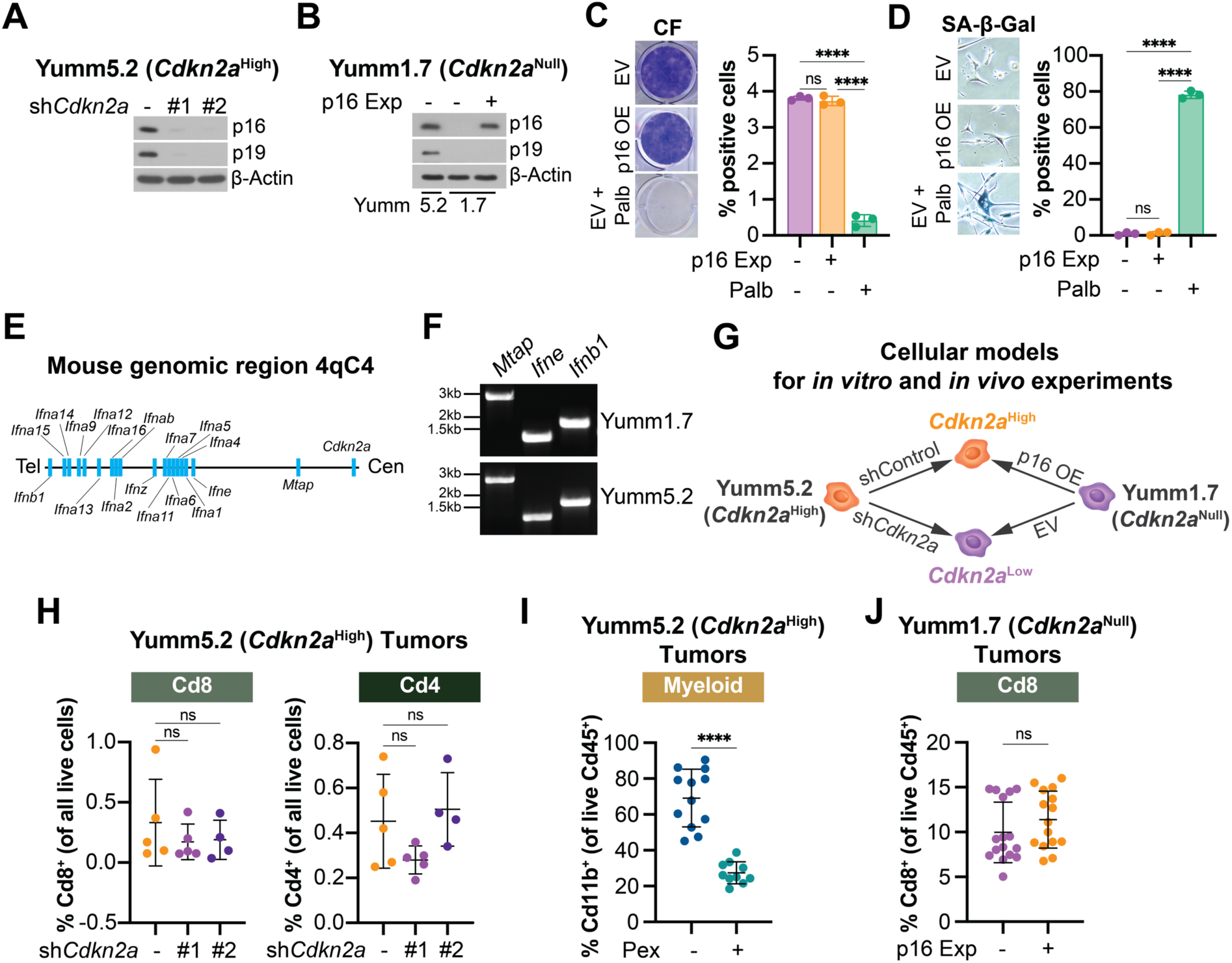
Cellular models used in this study. Related to Figure 1. **(A-B)** Immunoblot analysis of p16 and p19 in **(A)** Yumm5.2 (*Cdkn2a*^High^) with knockdown of GFP (shControl) or *Cdkn2a* (sh*Cdkn2a* hairpins #1 and #2) cells and **(B)** Yumm1.7 (*Cdkn2a*^Null^) cells expressing empty vector (EV) control of physiological levels of p16 (p16 Exp). β-Actin was used as a loading control. **(C-D)** Yumm1.7 (*Cdkn2a*^Null^) cells expressing empty vector (EV) control of physiological levels of p16 (p16 Exp). Induction of cellular senescence with palbociclib (Palb; 3μM, 10 days) was used as a positive control. **(C)** Representative images of colony formation (CF) assay (left) and quantification (right). Mean ± SD; one-way ANOVA. **(D)** Representative images of senescence-associated β-Galactosidase (SA-β-Gal) activity (left) and quantification (right). Mean ± SD; one-way ANOVA. **(E)** Schematic representation of the murine 4qC4 genomic region containing the *Cdkn2a* gene and gene neighbors. **(F)** Genomic DNA PCR amplification of indicated genes in Yumm1.7 and Yumm5.2 cell lines. **(G)** Schematic of cellular models. **(H)** Quantification of Cd8^+^ (left) and Cd4^+^ (right) T cells in tumors derived from Yumm5.2 (*Cdkn2a*^High^) cells with knockdown of GFP (shControl) or *Cdkn2a* (sh*Cdkn2a*, hairpins #1 and #2). Mean ± SD; one-way ANOVA. **(I)** Quantification of Cd11b^+^ cells in tumors derived from Yumm5.2 (*Cdkn2a*^High^) treated with pexidartinib (Pex, 45 mg/kg, o.g., 3×/week) or vehicle. Mean ± SD; student’s t-test**. (J)** Quantification of Cd8^+^ cells in tumors derived from Yumm1.7 (*Cdkn2a*^Null^) cells expressing empty vector (EV) control or physiological levels of p16 (p16 Exp). Mean ± SD; student’s t-test. * p<0.05, ** p<0.01, *** p<0.001, **** p<0.0001, ns = not significant.

**Figure S2.**
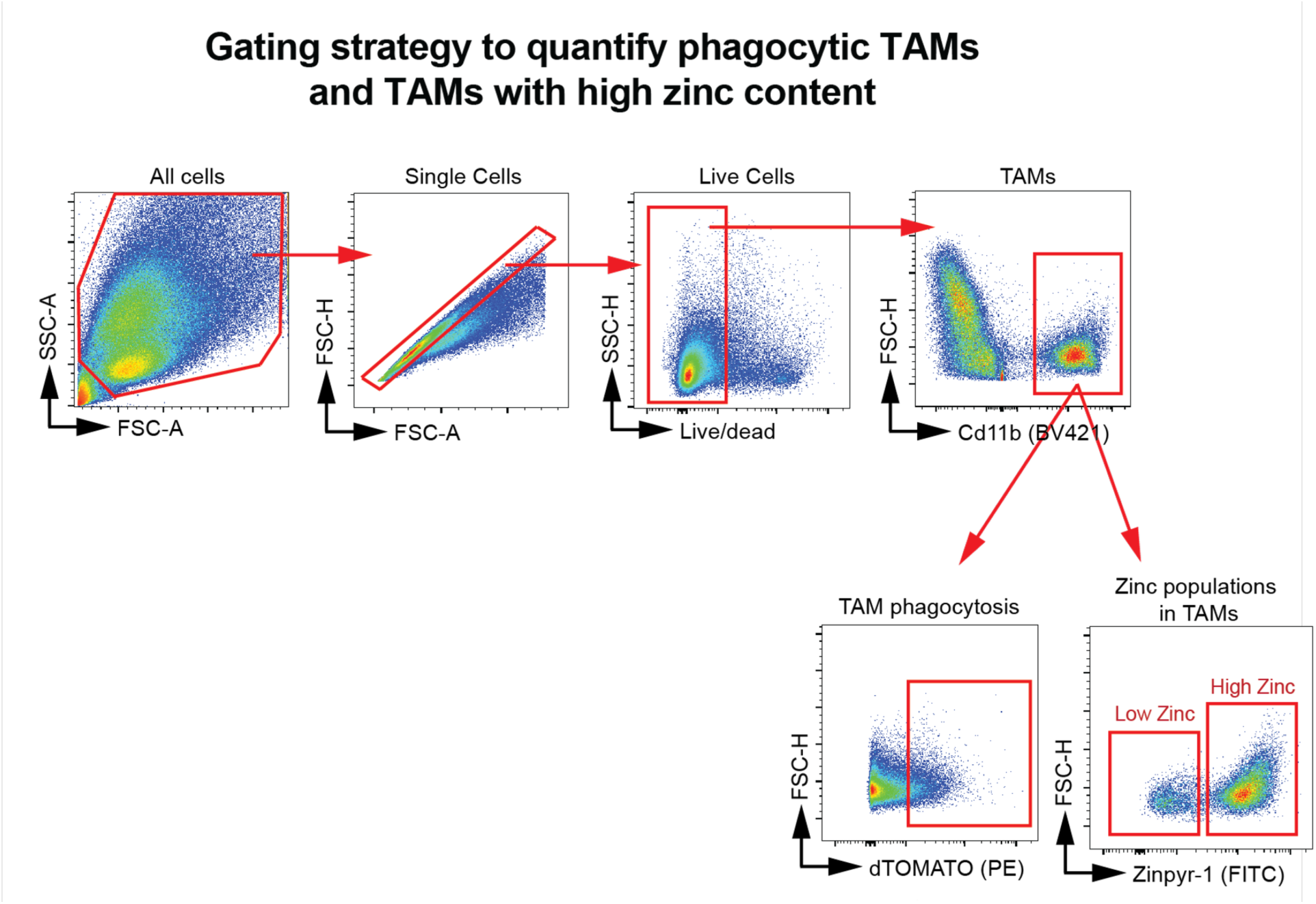
TAM gating strategy. Related to Figures 2, 6, 7, and S6. Figure shows the strategy used to identify live tumor-associated macrophages (TAMs), defined as the Cd11b⁺ population, and the subsequent gating to quantify: (1) High Zinc TAMs, defined as Cd11b⁺ TAMs with high Zinpyr-1 fluorescence, and (2) Phagocytic TAMs, defined as Cd11b⁺ TAMs positive for dTOMATO, the fluorophore used to label the cancer cells implanted into mice (see **Fig. 2E**).

**Figure S3.**
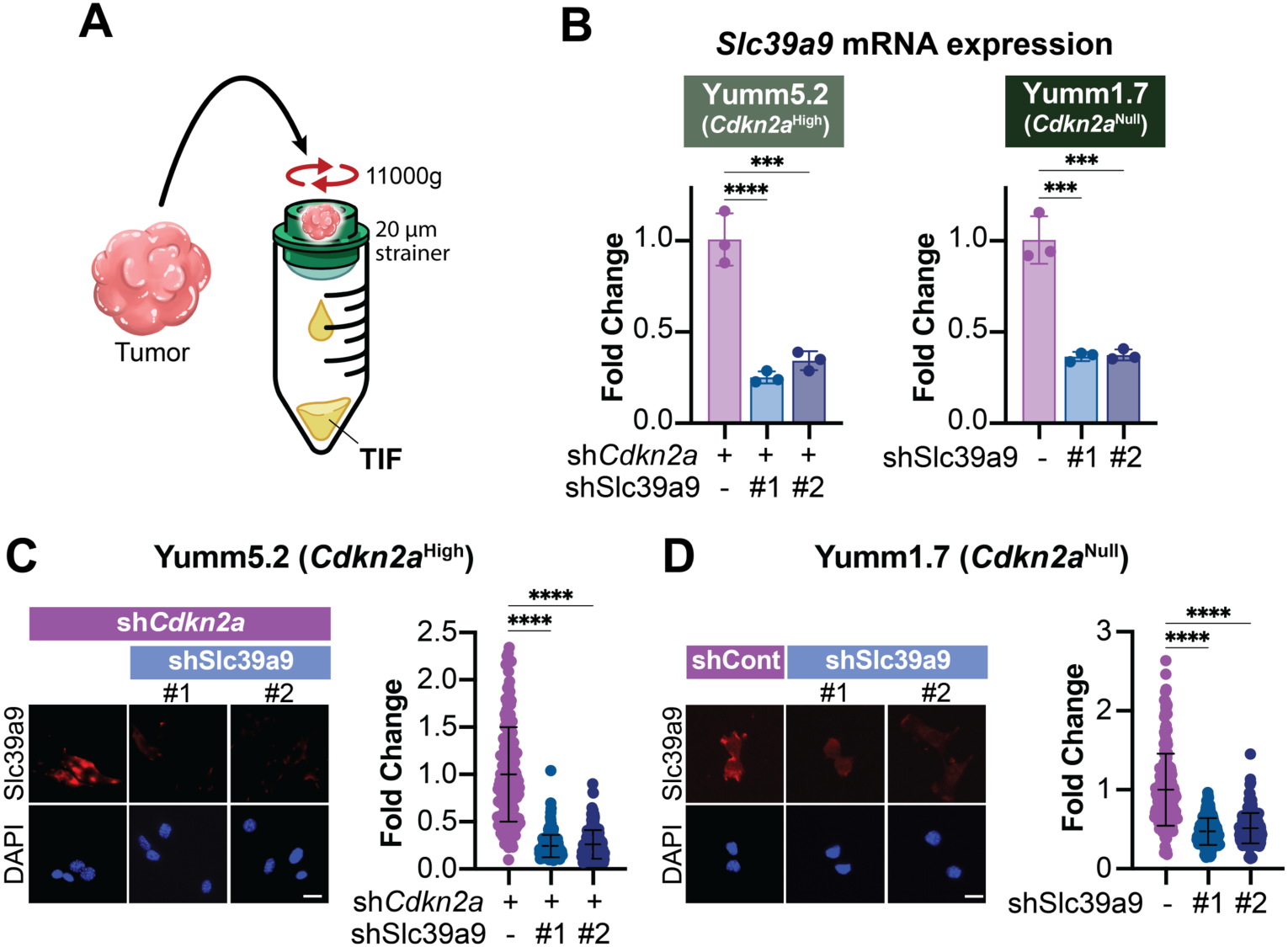
Tumor Interstitial Fluid (TIF) isolation; knockdown of Slc39a9 in cancer cells. Related to Figure 3. **(A)** Schematic of procedure to collect tumor interstitial fluid (TIF). **(B)** Slc39a9 mRNA expression in the indicated cell lines. Fold change to shControl group. Mean ± SD; one-way ANOVA. **(C-D)** Representative images (left) and quantification (right) of Slc39a9 immunofluorescence in **(C)** Yumm5.2 (*Cdkn2a*^High^) cells with knockdown of GFP (shControl) or *Cdkn2a* (sh*Cdkn2a*, hairpins #1 and #2). Mean ± SD (n=200/group); one-way ANOVA. **(D)** Yumm1.7 (*Cdkn2a*^Null^) cells expressing empty vector (EV) or physiological levels of p16 (p16 Exp). Mean ± SD (n=200/group); one-way ANOVA. Scale bars = 25μm.* p<0.05, ** p<0.01, *** p<0.001, **** p<0.0001, ns = not significant.

**Figure S4.**
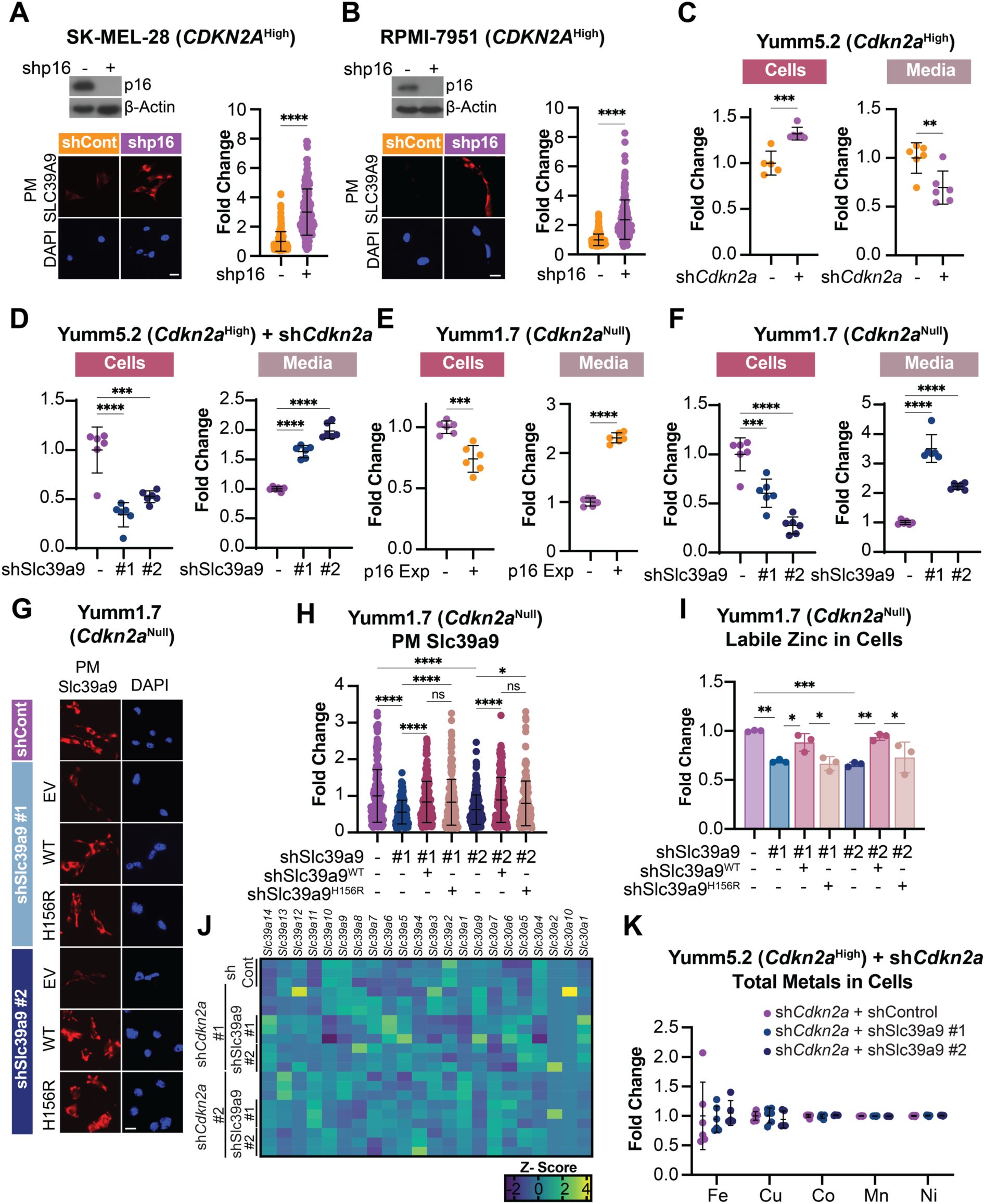
Knockdown of p16 in human melanoma cells increases plasma membrane SLC39A9; Slc39a9 regulates zinc uptake in *Cdkn2a*^Low^ cells. Related to Figure 4. **(A-B)** Top left-Immunoblot analysis of p16. β-Actin was used as a loading control. Bottom-representative images (bottom left), and quantification (bottom right) of the human melanoma cell lines **(A)** SK-MEL-28 (*Cdkn2a*^High^) and **(B)** RPMI-7951 (*Cdkn2a*^High^) with knockdown of GFP (shControl) or p16 (shp16). Mean ± SD (n=200/group); Student’s t-test. **(C-F)** Dot plot showing total zinc quantification by ICP-MS in cells (left) and media (right) in **(C)** Yumm5.2 (*Cdkn2a*^High^) cells with knockdown of GFP (shControl) or *Cdkn2a* (sh*Cdkn2a*), **(D)** Yumm5.2 (*Cdkn2a*^High^) cells with knockdown of *Cdkn2a* (sh*Cdkn2a*) alone or in combination with knockdown of GFP (shControl) or Slc39a9 (shSlc39a9, hairpins #1 and #2), **(E)** Yumm1.7 (*Cdkn2a*^Null^) cells expressing empty vector (EV) or physiological levels of p16 (p16 Exp), and **(F)** Yumm1.7 (*Cdkn2a*^Null^) cells with knockdown of GFP (shControl) or Slc39a9 (shSlc39a9, hairpins #1 and #2). Mean ± SD; Student’s t-test and one-way ANOVA. **(G-I)** Yumm1.7 (*Cdkn2a*^Null^) cells with knockdown of GFP (shControl) or Slc39a9 (shSlc39a9, hairpins #1 and #2) alone or in combination with empty vector (EV) control, wildtype (WT), or hypomorphic mutant (H156R) Slc39a9. **(G)** Representative images of Slc39a9 immuno-fluorescence on non-permeabilized cells to visualize plasma membrane (PM) Slc39a9. **(H)** Quantification of (G). Mean ± SD (n=200/group); one-way ANOVA. **(I)** Labile zinc quantification in cells. Mean ± SD; one-way ANOVA. **(J)** Heatmap of expression values for the indicated genes in the indicated Yumm5.2 isogenic cells. **(K)** Quantification of indicated metals by ICP-MS in Yumm5.2 (*Cdkn2a*^High^) cells with knockdown of *Cdkn2a* (sh*Cdkn2a*) alone or in combination with knockdown of GFP (shControl) or Slc39a9 (shSlc39a9, hairpins #1 and #2). Mean ± SD. Scale bars = 25μm.* p<0.05, ** p<0.01, *** p<0.001, **** p<0.0001, ns = not significant.

**Figure S5.**
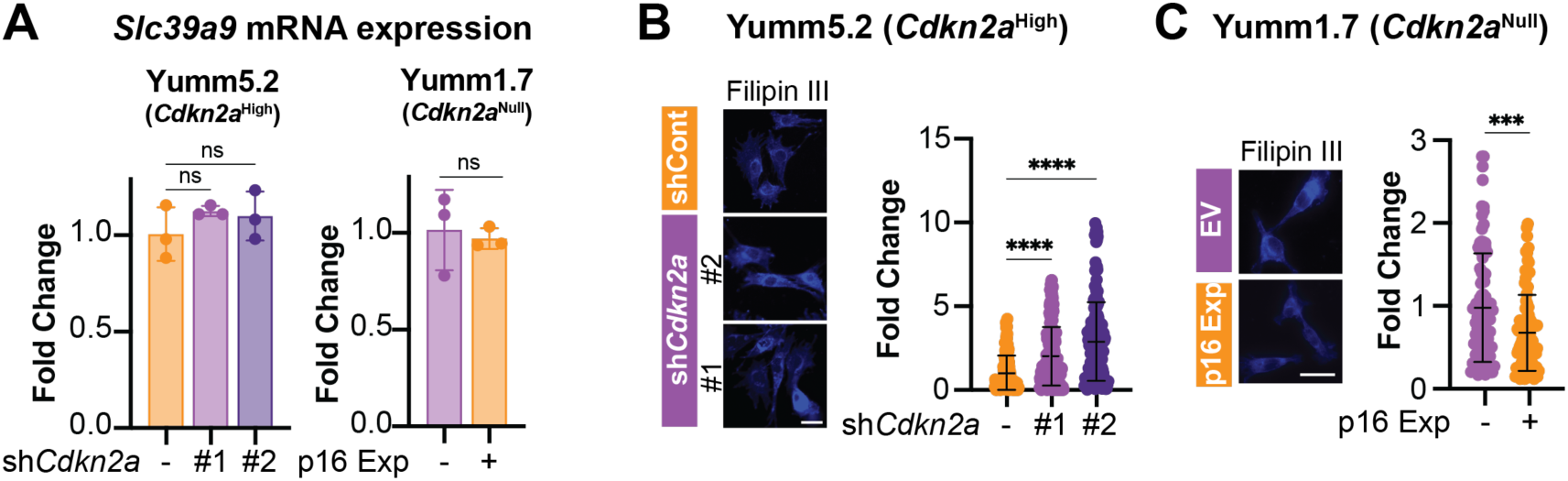
Modulation of *Cdkn2a*/p16 expression does not affect *Slc39a9* mRNA expression; *Cdkn2a*^Low^ cancer cells have decreased cholesterol. Related to Figure 5. **(A)** *Slc39a9* mRNA expression in the indicated cell lines. **(B-C)** Representative images of Filipin III staining to detect cholesterol (left) and quantification (right) in **(B)** Yumm5.2 (*Cdkn2a*^High^) cells with knockdown of GFP (shControl) or *Cdkn2a* (sh*Cdkn2a*, hairpins #1 and #2) and **(C)** Yumm1.7 (*Cdkn2a*^Null^) cells expressing empty vector (EV) control of physiological levels of p16 (p16 Exp). Mean ± SD (n=200/group); one-way ANOVA, student’s t-test. Scale bars = 25μm.* p<0.05, ** p<0.01, *** p<0.001, **** p<0.0001, ns = not significant.

**Figure S6.**
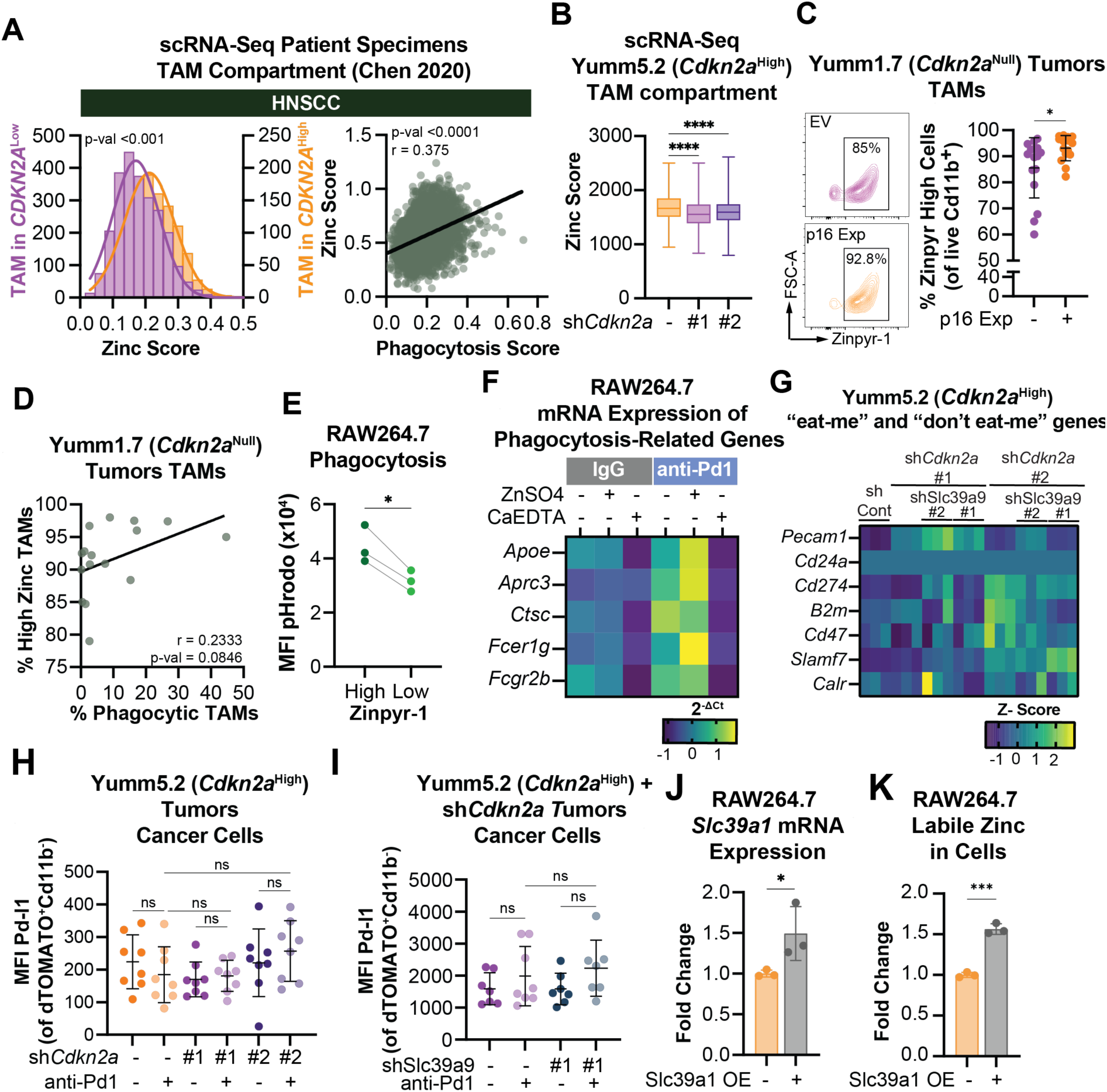
TAMs in *CDKN2A*^Low^ tumors have decreased zinc, correlating with phagocytosis activity; Zinc regulates the expression of phagocytic-related genes; Knockdown of *Cdkn2a* alone or in combination with Slc39a9 knockdown does not markedly affect the mRNA expression of ‘eat me’ and ‘don’t eat me’ signals; *Cdkn2a*^Low^ cancer cells alone or with knock-down of Slc39a9 do not have altered Pd-l1 expression *in vivo*. Related to Figure 6. **(A)** scRNA-Seq dataset from head and neck squamous cell carcinoma (HNSCC) patients stratified by *CDKN2A* expression in cancer cells. Histogram showing the zinc score in TAMs from the indicated scRNA-Seq datasets from melanoma patients stratified by *CDKN2A* expression in cancer cells (left) and zinc score correlation with phagocytic score (right). Linear mixed models and Pearson correlation **(B)** Box plot showing zinc signatures scores in the TAM compartment of the scRNA-Seq from Yumm5.2 (*Cdkn2a*^High^) cells with knockdown of *Cdkn2a* (sh*Cdkn2a*, hairpins #1 and #2). One-way ANOVA. **(C-D)** dTOMATO-labeled Yumm1.7 (*Cdkn2a*^Null^) cells expressing empty vector (EV) control or physiological levels of p16 (p16 Exp) were implanted into mice. **(C)** Representative flow cytometry plots of labile zinc in TAMs (left) and quantification (right). Mean ± SD; Student’s t-test. **(D)** Correlation between labile zinc in TAMs and TAM phagocytosis. Person’s correlation. **(E)** Mean fluorescence intensity (MFI) of pHrodoRed in RAW264.7 macrophages stratified based on high or low Zinpyr-1 staining of intracellular labile zinc. Mean ± SD; Student’s t-test. **(F)** Heatmap of phagocytosis-related genes measured by qPCR in RAW264.7 macrophages treated with ZnSO4 (30μg, 16h) or CaEDTA (3mM, 16h) in combination with IgG or anti-Pd1 (1 μg/mL,16h) in FBS-free media. **(G)** Heatmap of expression values for the indicated ‘eat me’ and ‘don’t eat me’ genes in Yumm5.2 (*Cdkn2a*^High^) cells with knockdown of *Cdkn2a* (sh*Cdkn2a*, hairpins #1 and #2) alone or in combination with knockdown of GFP (shControl) or Slc39a9 (shSlc39a9, hairpins #1 and #2). **(H)** Mice were implanted with dTOMATO-labeled Yumm5.2 (*Cdkn2a*^High^) cells with knockdown of GFP (shControl) or *Cdkn2a* (sh*Cdkn2a* #1 or #2). Mice were treated with IgG or anti-PD1 (200 µg, i.p., 3×/week). Dot plot of mean fluorescence intensity (MFI) of Pd-l1 in cancer cells (dTOMATO^+^ and Cd11b^−^). Mean ± SD; one-way ANOVA. **(I)** Mice were implanted with dTOMATO-labeled Yumm5.2 (*Cdkn2a*^High^) cells with knockdown of *Cdkn2a* (sh*Cdkn2a*) alone or in combination with knockdown of GFP (shControl) or Slc39a9 (shSlc39a9). Mice were treated with IgG or anti-PD1 (200 µg, i.p., 3×/week). Dot plot of mean fluorescence intensity (MFI) of Pd-l1 in cancer cells (dTOMATO^+^ and Cd11b^−^). Mean ± SD; one-way ANOVA. **(J)** Bar plot of *Slc39a1* mRNA expression in RAW264.7 macrophages expressing empty vector control or Slc39a1 overexpression plasmid (Slc39a1 OE). Mean ± SD as the fold change relative to shControl; Student’s t-test. **(K)** Bar plot of mean fluorescence intensity (MFI) of labile zinc in RAW264.7 macrophages expressing empty vector (EV) control or Slc39a1 overexpression plasmid (Slc39a1 OE). Mean ± SD as the fold change relative to shControl; Student’s t-test. * p<0.05, ** p<0.01, *** p<0.001, **** p<0.0001, ns = not significant.

**Figure S7.**
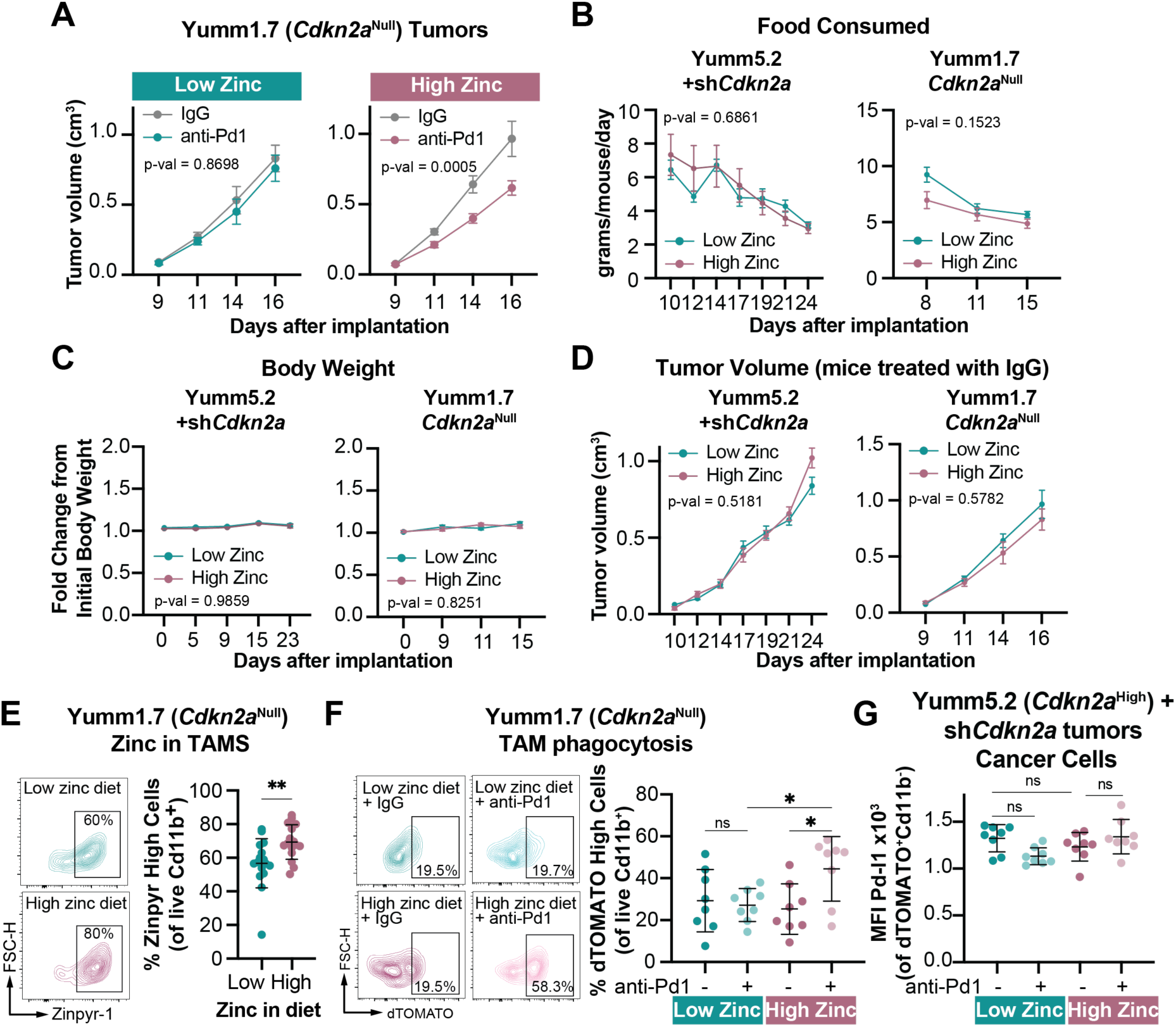
High zinc diet overcomes anti-Pd1 resistance in the Yumm1.7 *Cdkn2a*^Null^ model; Different zinc diets do not alter body weight or food consumed; Different zinc diets do not alter cancer cell Pd-l1 expression *in vivo*. Related to Figure 7. **(A)** Mice were fed a low (55 mg/kg) or high (300 mg/kg) zinc diet and implanted with Yumm1.7 (*Cdkn2a*^Null^) cells expressing empty vector (EV) control or physiological levels of p16 (p16 Exp). Mice were treated with IgG or anti-PD1 (200 µg, i.p., 3×/week). Longitudinal tumor growth in the indicated groups. Mean ± SEM (n=12/group); Mixed-effects model. **(B-D)** Mice were assigned to low (55 mg/kg) or high (300 mg/kg) zinc diets and then implanted with Yumm5.2 (*Cdkn2a*^High^) cells with knockdown of *Cdkn2a* (shCdkn2a, left) or Yumm1.7 (*Cdkn2a*^Null^) cells expressing empty vector (EV) control or physiological levels of p16 (p16 Exp, right). Mice were treated with IgG or anti-Pd1 (200 µg, i.p., 3×/week). **(B)** Food consumed. **(C)** Body weight. **(D)** Longitudinal tumor growth in IgG treated groups. Mean ± SEM (n=12/group); Mixed-effects model. **(E-F)** Mice were assigned to a high (300 mg/kg) zinc diet and implanted with Yumm1.7 (*Cdkn2a*^Null^) cells expressing empty vector (EV) control or physiological levels of p16 (p16 Exp). Once tumor establishment, mice were treated with IgG or anti-Pd1 (200 µg, i.p., 3×/week). **(E)** Representative flow cytometry plots (left) and quantification (left) of labile zinc in TAMs. Mean ± SD; student’s t-test. **(F)** Representative flow cytometry plots (left) and quantification (right) of the percentage of TAMs with dTOMATO signal (phagocytic TAMs). Mean ± SD; one-way ANOVA. **(G)** Mice were fed a high (300 mg/kg) zinc diet and implanted with dTOMATO-labeled Yumm5.2 (*Cdkn2a*^High^) cells with knockdown of GFP (shControl) or *Cdkn2a* (sh*Cdkn2a*). Once tumor establishment, mice were treated with IgG or anti-Pd1 (200 µg, i.p., 3×/week). Dot plot of mean fluorescence intensity (MFI) of Pd-l1 in cancer cells (dTOMATO^+^ and Cd11b^−^). Mean ± SD; one-way ANOVA. * p<0.05, ** p<0.01, *** p<0.001, **** p<0.0001, ns = not significant.

## Acknowledgements

This work was supported by grants from the National Institutes of Health (R37CA240625 to KM. Aird, R01CA259111 to KM. Aird and NW. Snyder, 1R01CA301573-01 to KM. Aird, N. Hempel, and R. Buj, R01DE031729 and P50CA254865 to R. Bao, and T32 CA082084 to RE. Dadey), the Congressionally Directed Medical Research Program (HT9425-23-1-0436 to KM. Aird and OC230324 to NK. Tangudu), the Melanoma Research Foundation (to R. Buj), the HERA Ovarian Cancer Foundation (to A. Amalric, A. Uboveja, and NK. Tangudu), the University of Chicago Cancer Center Support Grant (to A. Muir), the Ludwig Center for Metastasis Research (to A. Muir), the MaryEllen Connelan Award (to JJ. Apiz Saab), the Robert C and Mary Jane Gallo Scholarship Fund (to JJ. Apiz Saab) and the Harper Fellowship at the University of Chicago (to JJ. Apiz Saab). We utilized resources from the Hillman Cancer Center shared facilities, partially supported by award P30CA047904 including the Animal Facility, the Cytometry Facility (RRID:SCR_025361), the Biostatistics Facility (RRID:SCR_025355), and the Cancer Bioinformatics Services (RRID:SCR_025355). The Cancer Bioinformatics Services were additionally supported by the University of Pittsburgh Center for Research Computing, specifically the HTC high-performance computing cluster, which is supported by NIH (S10OD028483). We also used the University of Pittsburgh Single Cell Core Facility (RRID:SCR_025110), whose services and instruments were in part graciously supported by the University of Pittsburgh and the Department of Medicine. Similarly, we also used the Health Sciences Sequencing Core at UPMC Children’s Hospital of Pittsburgh (RRID:SCR_023116), which resources were generously supported by the University of Pittsburgh, the Office of the Senior Vice Chancellor for Health Sciences, the Department of Pediatrics, the Institute for Precision Medicine, and the Richard K. Mellon Foundation for Pediatric Research. Finally, we extend our gratitude to Francisco J. Vazquez Moreno for his exceptional contribution to the graphic illustrations included in this paper.

## Author Contributions

Raquel Buj: Conceptualization, Investigation, Methodology, Visualization, Writing – Original Draft, Review & Editing, Funding acquisition. Aidan R. Cole: Investigation, Methodology, Writing – Review & Editing. Jeff Danielson: Investigation, Methodology. Jimmy Xu: Investigation, Methodology. Akash Kishore: Investigation, Methodology. Katarzyna M. Kedziora: Investigation, Methodology. Jie Chen: Investigation, Methodology. Baixue Yang: Investigation, Methodology, Writing – Review & Editing. David Barras: Investigation, Methodology. Apoorva Uboveja: Investigation, Methodology, Writing – Review & Editing. Amandine Amalric: Investigation, Methodology, Writing – Review & Editing. Juan J. Apiz Saab: Investigation, Methodology. Jayamanna Wickramasinghe: Investigation, Methodology. Naveen Kumar Tangudu: Investigation, Methodology, Writing – Review & Editing. Evan Levasseur: Investigation, Methodology, Writing – Review & Editing, Hui Wang: Investigation, Methodology. Aspram Minasyan: Investigation, Methodology. Rebecca E. Dadey: Investigation, Methodology. Allison C. Sharrow: Investigation, Methodology. Sheryl Kunning: Investigation, Methodology. Frank P. Vendetti: Investigation, Methodology. Dayana B. Rivadeneira: Investigation, Methodology. Christopher J. Bakkenist: Investigation, Methodology, Writing – Review & Editing. Tullia C Bruno: Investigation, Methodology, Writing – Review & Editing. Greg M. Delgoffe: Investigation, Methodology, Writing – Review & Editing. Nadine Hempel: Investigation, Methodology, Writing – Review & Editing, Funding acquisition. Nathaniel W. Snyder: Investigation, Methodology, Writing – Review & Editing. Riyue Bao: Investigation, Methodology, Writing – Review & Editing. Adam C. Soloff: Investigation, Methodology, Writing – Review & Editing. John M. Kirkwood: Investigation, Methodology, Writing – Review & Editing. Denarda Dangaj Laniti: Investigation, Methodology, Writing – Review & Editing, Andrew V. Kossenkov: Investigation, Methodology, Writing – Review & Editing. Alexander Muir: Investigation, Methodology, Writing – Review & Editing. Jishnu Das: Investigation, Methodology, Writing – Review & Editing. Diwakar Davar: Investigation, Methodology, Writing – Review & Editing. Clementina Mesaros: Investigation, Methodology, Writing – Review & Editing. Katherine M. Aird: Conceptualization, Investigation, Visualization, Writing – Original Draft, Review & Editing, Supervision, Project Administration, Funding Acquisition. All authors reviewed the results and approved the final version of the paper.

## Declaration of Interests

The authors declare no competing interests.

## MATERIAL AND METHODS

### Melanoma Patient Cohort

This cohort comprises 248 cutaneous melanoma patients, pooled from five independent, publicly available studies^35,108,189–191^, with publicly available pre-treatment RNA-Seq and overall survival data. Details are provided in **Table S6**. Briefly, eligible patients had resectable stage III melanoma treated with neoadjuvant immunotherapy or unresectable stage III/IV treated with systemic immunotherapy. Treatment regimens included anti-PD1 monotherapy or combination therapy with anti-CTLA-4 or TLR9 agonists. For **Fig. 1A**, patients were stratified according to *CDKN2A* expression. Expression levels of the target genes were categorized as follows: patients in or above the third tertile were considered to have high expression, while those in or below the first tertile were considered to have low expression. For **Fig. 2B**, T cell inflamed signature^192^ and phagocytosis signature scores (this paper, see **Table S1**) were calculated based on the sum of the genes in each signature. Patients were stratified according to median score.

### Kaplan-Meier plotter (KMP) Cancer Patient Cohort

Survival data from cancer patients was downloaded from Kaplan-Meier (KM) plotter ‘Start KM Plotter for immunotherapy’ (kmplot.com)^193–195^ in January 2025. The analysis was performed using the following parameters: the gene symbol was set to *CD8A* with *Filter by mean expression* of patients with high expression of *ITGAM* (**Fig. 1G**). Patients were split based on *mean* expression. *Overall survival* was selected as the survival metric, with no restrictions on follow-up thresholds. The analysis was restricted to patients who had received *all* anti*-*PD1 therapy and included *all* tumor types and sex. The resulting data was exported as text for subsequent analysis in Graph Prism 10.

### Patient scRNA-Seq data sets and processing

The Jereby-Arnon 2018 melanoma dataset^196^ and the Chen 2020 head and neck squamous cell carcinoma (HNSCC) dataset^197^ were downloaded from the Gene Expression Omnibus (GSE115978 and GSE150430, respectively). For sample processing, genes were retained if expressed in at least three cells, cells were kept if they had more than 200 features, and cells with >20% mitochondrial reads were removed. Library size bias was corrected using Seurat’s NormalizeData, scaling expression values to 10,000 counts per cell and log-transforming the data. Data was then scaled using Seurat’s ScaleData with all genes as features. Cancer cells and macrophages were annotated based on publicly available cell type labels. Tumor samples were then stratified into *CDKN2A*^High^ and *CDKN2A*^Low^ groups based on *CDKN2A* median expression in the cancer cell compartment. Signature scores were calculated as the mean expression of genes in the zinc signature and phagocytosis signature annotated in this paper (**Table S1**) for each cell in the macrophage or cancer cell compartment.

### Cell Lines and culture conditions

All cell lines used in this study were purchased from ATCC and were used within 50 passages. ATCC performs cell line authentication by short tandem repeat (STR) profiling. Cells were cultured within 3 months of receipt or resuscitation. The murine melanoma cell lines Yumm5.2 (ATCC, catalog no. CRL-3367, RRID: CVCL_JK43) and Yumm1.7 (ATCC, catalog no. CRL-3362, RRID: CVCL_JK16) were cultured in DMEM and Ham’s F-12 50/50 media (Corning, catalog no. 10-092-CV), supplemented with nonessential amino acids (Corning, catalog no. 25-025-Cl). The murine macrophage cell line RAW 264.7 (ATCC, catalog no. TIB-71 ™, RRID: CVCL_0493), the human melanoma cell line SK-MEL-28 (ATCC, catalog no. HTB-72, RRID: CVCL_0526), and the human embryonic kidney cell line HEK293-FT (ATCC, catalog no. CRL-3216, RRID: CVCL_6911) were cultured in DMEM (Corning, catalog no. 10-013-CV), while the human melanoma cell line RPMI-7951 (ATCC, catalog no. HTB-66, RRID: CVCL_1666) was cultured in MEM (Corning, catalog no. 10-009-CV). All cell lines were supplemented with 10% FBS (BioWest, catalog no. S1620) and 1% Penicillin/Streptomycin (Gibco, catalog no. 15-140-122) and maintained at 37 °C with 5% CO_2_. To ensure the absence of Mycoplasma contamination, all cell lines were tested monthly as described in^198^.

### Plasmid information and cloning

To perform knockdown experiments, short hairpin RNAs (shRNAs) targeting specific genes were used. The shRNA sequences used in this study included: human shp16 (TRCN0000010482), mouse sh*Cdkn2a* #1 (TRCN0000077816) and #2 (TRCN0000362595), and mouse shSlc39a9 #1 (TRCN0000346746) and #2 (TRCN0000346750). The hairpins for shp16 and shSlc39a9 #1 and #2 were purchased pre-cloned into the lentiviral vectors expressing puromycin resistance (Sigma-Aldrich). The sh*Cdkn2a* #1 and #2 hairpins sequences (TRCN0000077816 and TRCN0000362595, respectively) were obtained from Integrated DNA Technologies (IDT) as desalted oligos and subsequently cloned into the lentiviral pLKO.1 vector expressing blasticidin resistance (Addgene, catalog no. 26655). Briefly, 2 µg of pLKO.1 vector was digested with 1 µL of EcoRI-HF (New England Biolabs, catalog no. R3101S), 1 µL of AgeI-HF (New England Biolabs, catalog no. R3552S), and 1 µL of rCutsmart 10X Buffer (New England Biolabs, catalog no. B6004S) in a final reaction volume of 10 µL. The digestion was incubated for 1 hour at 37°C. The linearized vector was separated by electrophoresis on a 1% agarose (IBI Scientific, catalog no. IB70070) gel and purified using the Qiagen Gel Extraction Kit (Qiagen, catalog no. 28704), following the manufacturer’s instructions. Simultaneously, shRNA oligos were reconstituted to a final concentration of 100 µM in molecular-grade water (Corning, catalog no. 46-000-CM) and annealed by incubating 11.25 µL of each oligo with 2.5 µL of 10X annealing buffer (1M NaCl, 100 mM Tris, pH 7.4). The reaction was performed in a thermal cycler (Bio-Rad, T100™ Thermal Cycler, serial no. 621BR59583) starting at 95°C for 5 minutes, followed by a ramp-down to 25°C at 5°C/min. The ligation reaction was prepared by combining 50 ng of purified pLKO.1 vector with 1 µL of annealed shRNA oligos (diluted 1:400 in 0.5X annealing buffer), 1 µL of T4 ligase (New England Biolabs, catalog no. M0202T), and 1 µL of T4 ligase buffer (New England Biolabs, catalog no. B0202S) in a final volume of 10 µL. The reaction was incubated at room temperature for 3 hours, after which 5 µL of the ligation mixture was used to transform One Shot™ Stbl3™ Chemically Competent *E. coli* (Thermo Fisher Scientific, catalog no. C737303) following the manufacturer’s instructions. Control lentiviral vectors, (Addgene, catalog no. 110419, RRID: Addgene_110419 and Addgene, catalog no. 30323, RRID: Addgene_30323), were used for both human and mouse experiments. For overexpression experiments, lentiviral plasmids were synthe-sized by Twist Bioscience. The Twist Bioscience empty vector (EV) was used as the control in overexpressing experiments. The pLX304-dTOMATO lentiviral plasmid was generously provided by the Gomes Lab at the H. Lee Moffitt Cancer Center.

### Lentiviral Packaging and Infection conditions

Lentiviral vectors were packaged as follows: 10 µg of the corresponding lentiviral vector was mixed with 7.5 µg of psPAX2 (Addgene, catalog no. 12260, RRID: Addgene_12260) and 7.5 µg of pMD2.G (Addgene, catalog no. 12259, RRID: Addgene_12259) in 1.5 mL of Opti-MEM™ media (Gibco, catalog no. 31985070) and incubated for 5 minutes at room temperature. In parallel, 50 µL of Lipofectamine™ 2000 Transfection Reagent (Invitrogen, catalog no. 11668019) was diluted in 1.5 mL of Opti-MEM™ media and incubated for 5 minutes at room temperature. The Lipofectamine mixture was added dropwise to the plasmid mixture, gently swirled to mix, and incubated for 20 minutes at room temperature. The Lipofectamine-lentiviral complexes were then added dropwise to a 60-70% confluent 10 cm dish of HEK293-FT cells, seeded 24 hours in advance and containing 7 mL of regular media, and incubated at 37°C with 5% CO2. Six to eight hours later, the media was replaced with 9 mL of fresh regular media and incubated at 37°C with 5% CO_2_. Seventy-two hours later, the viral supernatants were collected, filtered through a 45 µm filter (Fisherbrand™, catalog no. 09-720-005), and used to infect target cells for 16 hours. Infected cells were selected for 3 days with 1 µg/mL puromycin (Thermo Fisher Scientific, catalog no. A1113802) for SK-MEL-28 and RPMI-7951, 2 µg/mL puromycin for RAW 264.7 cells, and 3 µg/mL puromycin and/or 3 µg/mL blasticidin (InvivoGen, catalog no. ant-bl-05) for Yumm5.2 and Yumm1.7 cells.

### Mouse Models

All mice were housed in a HEPA-filtered ventilated rack system at the Animal Facility of the Assembly Building of The Hillman Cancer Center, University of Pittsburgh School of Medicine. Mice were maintained on a 12-hour light/dark cycle at 21 ± 2 °C and were group-housed up to 5 mice per cage. All animal procedures were approved by the Institutional Animal Care and Use Committee (IACUC) of the University of Pittsburgh.

To investigate the role of *CDKN2A* in the response to immunotherapy, we developed two complementary subcutaneous syngeneic melanoma models. In the first model, *Cdkn2a* expression was knocked down in the *Cdkn2a*^High^ murine melanoma cell line Yumm5.2. In the second model, p16 was overexpressed in the *Cdkn2a*^Null^ melanoma cell line Yumm1.7 to physiological levels that did not induce a senescence-associated phenotype. To investigate the role of SLC39A9 in resistance to anti-Pd1 treatment in *Cdkn2a*^Low^ cancer cells, we used Yumm5.2 cells with dual knockdown of *Cdkn2a* and *Slc39a9* (sh*Cdkn2a*n + sh*Slc39a9*), as well as Yumm1.7 cells with *Slc39a9* knockdown (sh*Slc39a9*). In all models described above, 350,000 cells were subcutaneously implanted into the right flank of 6– 8 weeks old male C57BL/6J mice (The Jackson Laboratory, strain #000664, RRID: IMSR_JAX:000664). To assess the role of T cells in anti-Pd1 response, 350,000 Yumm5.2 sh*Cdkn2a*/sh*Slc39a9* double knockdown cells were implanted into B6.129s7-*Rag1*^tm1Mom^/J mice (The Jackson Laboratory, strain #002216, RRID: IMSR_JAX:002216). For co-injection experiments, 350,000 Yumm5.2 sh*Cdkn2a* cells were mixed with 35,000 RAW264.7 cells overexpressing Slc39a1 and subcutaneously implanted into the right flank of 6-8 weeks old male NOD.Cg-Prkdc^scid^ Il2rg^tm1Wjl^/Szj (The Jackson Laboratory, strain #005557, RRID:IMSR_JAX:005557). In all cases, control groups received cells overexpressing corresponding shControl (shGFP) or empty vector (see *Plasmid Information and cloning* section).

In all cases described above, cells were resuspended in 200 µL DPBS (Corning, catalog no. 21-031-CV) and subcutaneously implanted using a 1 mL tuberculin syringe and a 25G needle (Becton Dickinson, catalog no. 309626). Tumor formation was monitored, and all models developed palpable tumors within 8–14 days of implantation. Tumor length (L) and width (W) were measured three times a week once palpable tumors appeared using an electronic caliper set in mm (Adoric, model number# KC24, resolution 0.1mm ± 0.2mm). Tumor volume was calculated using the formula: ½ (L × W²) where L > W. Mouse body weight was measured weekly using an electronic scale. The experimental endpoint for longitudinal tumor growth studies was defined as the point when the first mouse reached IACUC-approved criteria: either a tumor volume exceeding 1 cm³ or a 20% loss of body weight, whichever occurred first. For survival studies, the endpoint for each animal was similarly determined based on IACUC guidelines.

### Mice treatments

For anti-Pd1 experiments, mice were administered 200 µg of anti-mouse Pd1 (Leinco Technologies, catalog no. P362) or placebo IgG2a (Leinco Technologies, catalog no. I-1177). diluted in DPBS to a final volume of 200 µL, by intraperitoneal injection three times per week using insulin syringes (Excel International, catalog no. 26028). For macrophage depletion, mice were treated with 45 mg/kg pexiclartinib (Selleck Chemicals, catalog no. S7818), diluted in a vehicle solution containing 45% PE300 (Sigma-Aldrich, catalog no. 202371), 45% ddH₂O, 5% Tween-80 (Sigma-Aldrich, catalog no. P4780), and 5% DMSO (Fisher BioReagents, catalog no. BP231-100), administered via oral gavage at a final volume of 150 µL, three times per week. Treatment was delivered using a 1 mL syringe (Becton Dickinson, catalog no. 309659) and a reusable animal feeding needle (Cadence Science, catalog no. 7910; 2.5 mm ball diameter, 3.81 cm length, 20G, curved), which were autoclaved between uses. Control mice received the same volume of vehicle solution following the same dosing schedule. All treatments described above were initiated upon detection of palpable tumors. For dietary zinc experiments, standard rodent chow (LabDiet, catalog no. 5P76) containing 110 ppm (110 mg/Kg) zinc was replaced with custom-made diets from LabDiet. Mice were fed either a high-zinc diet (LabDiet, catalog no. 5SNB) containing 300 ppm (300 mg/Kg) zinc or a low-zinc diet (LabDiet, catalog no. 5SNA) containing 55 ppm (55 mg/Kg) zinc. Both experimental diets were identical in composition to the standard chow, except for the zinc content. Mice were acclimated to the new diet for 10 to 15 days prior to tumor cell implantation and continued on the assigned diet throughout the experiment. The amount of food per cage was measured before and after replenishing the feeder one to two times per week to estimate food intake per mouse and day on each diet.

### *In vivo* and *in vitro* phagocytosis assay

For *in vivo* phagocytosis assays, cancer cells were infected with a lentivirus encoding the dTOMATO fluorescent protein (see *Lentiviral Packaging and Infection conditions* section). dTOMATO-positive cancer cells were sorted prior to implantation into mice. The level of dTOMATO expression varied by less than 1% across the different cancer cell groups implanted. Once tumors reached a volume of 250–300 mm³, tumors were harvested, dissociated (see *Tumor Dissociation* section), stained, and analyzed by flow cytometry (see *Immunostaining and Flow Cytometry* section). The percentage of phagocytic TAMs was defined as the proportion of live Cd11b⁺ cells that also expressed dTOMATO (**Fig. 2E**). For *in vitro* phagocytosis assays, cancer cells were labeled with CellTrace™ Violet (Thermo Fisher, Cat. No. C34557) according to the manufacturer’s instructions. One million CellTrace-labeled cancer cells were added to 300,000 IFNγ-activated RAW264.7 macrophages (100 ng/mL IFNγ for 24 hours) previously plated, and co-cultured for 24 hours. After incubation, cells were harvested by scraping, stained, and analyzed by flow cytometry (see *Immunostaining and Flow Cytometry* section). The percentage of phagocytic RAW264.7 cells was defined as the live Cd11b⁺ population that coexpressed CellTrace Violet (**Fig. 2E**). Finally, the in vitro phagocytic activity was also determined with the stablished pHrodo red E.coli particles (**Fig. S6E**) as follows: 75,000 RAW264.7 cells seeded in 12-well plates were washed with PBS and media replaced for 1.5 mL of 200 µg/mL of pHrodo red E.coli particles (Invitrogen, catalog no. P35361) resuspended in media without FBS and incubated 1h at 37°C and 5% CO_2_. Cells were scrapped out and stained with zinpry-1 and Live/Dead Fixable Near-IR Dead Cell Stain (see *In vitro labile Zinc quantification* section).

### Tumor disassociation

To obtain a single-cell suspension, dissected tumors were enzymatically and mechanically disrupted. First, the tumors were mechanically minced using surgical disposable scalpels (Surgical Design, catalog no. SCALPEL 12B) into pieces smaller than 5 mm in diameter on a 6 cm plate. Next, 2 mL of enzymatic cocktail [8 mg of collagenase IV (Gibco, catalog no. 17104-019), 800 µL of dispase (Stem Cell Technologies, catalog no. 07913), and 25 µg of DNase (Sigma Aldrich, catalog no. D4263-5VL) diluted in FBS-free MEM (Corning, catalog no. 10-009-CV)] was added to the minced tumors. The mixture was incubated for 20 minutes at 37°C in a non-CO₂ incubator. Minced tumor pieces were then further disrupted by smashing them between two fully frosted cytology slides (Fisherbrand, catalog no. 12-550-401). The resulting single-cell suspension was filtered through a 70 µm strainer (Fisherbrand, catalog no. 22-363-548) and resuspended in 5 mL of FBS-free media. The cells were centrifuged at 350 g for 5 minutes at 4°C and resuspended in 1x RBC lysis buffer (BD, catalog no. 555899). The RBC lysis reaction was incubated for 2–5 minutes at room temperature, after which the reaction was stopped by adding 5 mL of FBS-free media. The cells were then centrifuged at 350 g for 5 minutes at 4°C, resuspended in DPBS, and were ready for subsequent procedures.

### Single-cell RNA-Sequencing

Yumm5.2 sh*Cdkn2a* hairpins #1 and #2, and control shGFP cells were subcutaneously implanted into five mice per condition (see *Mice Models* section). Once reached a volume of 0.5 cm³, tumors were harvested and processed into a single-cell suspension (see *Tumor cell disassociation* section). Single-cell suspensions were first stained with a 1:4000 dilution of viability dye (ThermoFisher Scientific, catalog no. 65-0863-14) in DPBS for 20 minutes on ice and in the dark. Then, cells were centrifuged at 350 g for 5 minutes at 4°C and stained with 2 µg/mL of anti-Cd45 PE-conjugated antibody (ThermoFisher Scientific, catalog no. 12-0451-83), diluted in EasySep™ buffer (Stemcell Technologies, catalog no. 20144) for 20 minutes on ice and protected from light. After centrifugation at 350 g for 5 minutes at 4°C, cells were resuspended and pooled by sample group in 5 mL of FBS-free MEM media (Corning, catalog no. 10-009-CV). Single live PE-positive (Cd45+) cells were sorted using a Sony MA900 cell sorter (serial no. 0714313) and collected in 0.04% BSA/PBS for downstream processing. cDNA Libraries were prepared at the University of Pittsburgh Single Cell Core Facility. Briefly, viability was confirmed with greater than 80% live cells using a Nexcelom automated cell counter (serial no. Auto2000-201-1019). Samples were then labeled with TotalSeq-C antibody hashtags (Biolegend, catalog no. 155881, 155883, 155885, 155887, 155889, 155891) following manufacturer’s instructions. Equal numbers of cells from each sample were then pooled together and loaded onto the Chromium controller (serial no. CXVG11091) for single-cell capture and barcoding. cDNA library preparation was carried out using the Chromium Next GEM Single Cell 5’ Reagent Kits v2 (10X Genomics, catalog no. PN-1000265) following the manufacturer’s protocol. The sizes of the library pools were analyzed using a Standard Sensitivity NGS Fragment Analyzer Kit (Agilent) followed by fluorescent quantification using the Promega QuantiFluor ONE dsDNA kit on the Infinite F Nano+ (Tecan). The library pools were normalized per manufacturer protocol (Illumina). Sequencing was performed on the NovaSeq 6000 platform (serial no. A00524), with 28bp/90bp paired-end reads. The sequencing data was demultiplexed with 10X Genomics Cell Ranger mkfastq version v6.0.2. The resulting sequence data was then aligned to the mouse reference genome (mm10) using cellranger multi (v.7.1.0) to generate count matrices for each condition. Analysis was conducted using R (v.4.3.0). Seurat’s was used for quality control, data normalization, and data analysis. The data was normalized using the NormalizeData function, following which the top 2000 variable genes were chosen using the ‘vst’ parameter setting of the FindVariableFeatures function. The 6 data objects were merged and used for further analysis. Celltypes were annotated using ImmGen data and the SingleR package. Gene Set Enrichment Analysis (GSEA) was performed using ranked gene lists (log2foldchange * log10pval) using the GOBP_PHAGOCYTOSIS mouse gene set from MSigDB. Pathway signature scores for each cell were computed using the ssGSEA framework as implemented in the escape package. Raw and processed RNA-seq data can be found on Gene Expression Omnibus (GSE291530).

### Immunostaining and flow cytometry

Tumors were harvested and processed into a single-cell suspension (see *Tumor disassociation* section). Single-cell suspensions were first blocked with 5 µg/mL of Cd16/Cd32 (Invitrogen, catalog no. 14016185) diluted in DPBS for 10 minutes on ice. Cells were stained for 20 minutes at 4°C protected from the light after which cells were fixed in 4% paraformaldehyde (Sigma Aldrich, catalog no. 158127-3Kg) for 15 minutes at 4°C, shielded from light, and resuspended in FACS buffer (1x PBS, 10 mM HEPES [Gibco, catalog no. 15630080], 1% heat-inactivated FBS [Sigma Aldrich, catalog no. F4135-100ML], 0.05% sodium azide [Sigma Aldrich, catalog no. S2002-25G]). Stained cells were immediately analyzed on a Fortessa (Beckman Coulter, serial no. H64717700020). FlowJo version 10 was used for further analysis. Antibodies and dyes used in this study include: 0.5 µg/mL of anti-Cd11b Brilliant Violet 421-conjugated antibody (Biolegend, catalog no. 101236), 2 µg/mL of anti-Pd-l1 PE/Cyanine5-conjugated antibody (Biolegend, catalog no. 124344), 0.5 µg/mL of anti-Cd8a FITC-conjugated antibody (Thermo Fisher Scientific, cat no. 11-0081-82), 0.5 µg/mL of anti-Cd4 Brillian Violet 421-conjugated antibody (Biolegend, catalog no. 100437), 2 µg/mL of anti-Cd45 PE-conjugated antibody (ThermoFisher Scientific, catalog no. 12-0451-83), 50 µM Zinpyr-1 (Cayman Chemical, catalog no. 15122). Antibodies were diluted in DPBS, containing a 1:1000 dilution of Live/Dead Fixable Near-IR Dead Cell Stain kit (Invitrogen, catalog no. L34976).

### Isolation of plasma and tumor interstitial fluid (TIF)

Blood from melanoma patients treated with anti-PD1 immunotherapy (**Fig. 3F**) was collected by venipuncture into ACD-A tubes (Becton-Dickinson, catalog no. 364606) as part of an IRB-approved biorepository protocol (HCC 20-2019, MOD20010266-016). Mouse blood was collected via lancet check bleed (Braintree Scientific, catalog no. GR 4MM) into EDTA tubes (SAI Infusion Technologies, catalog no. K3EDTA). Plasma was separated by centrifugation at 1500g for 10 minutes at 4°C and stored at −80°C for later analysis. After bleeding, animals were euthanized by cervical dislocation, and tumors were harvested, weighed, rinsed briefly in saline (150 mM NaCl), and excess saline removed using filter paper (Cytiva, catalog no. 1001-813). TIF was isolated using a previously described centrifugal method^199^ (**Fig. S3A**). Briefly, tumors were placed on a 20 µm strainer (pluriSelect, catalog no. 43-50020-03) within a sterile 50 mL conical tube (FisherbrandTM, catalog no. 06-443-19) and centrifuged at 11,000g for 10 minutes at 4°C. TIF was collected from the conical tube, rapidly frozen in liquid nitrogen, and stored at −80°C. TIF volumes ranged from 5 to 100 µL, <20% of the tumors did not yield any fluid. The interval from euthanasia to centrifugation was <2 minutes per mouse.

### Harvest cells and conditioned media for ICP-MS and labile zinc quantification

For ICP-MS and labile zinc quantification, cells were seeded in 10 cm tissue culture dishes (Cellstar, catalog no. 82050-916) at densities of 500,000 and 300,000 cells per dish, respectively. Cells were seeded in complete media and incubated for 48 hours (see *Cell Lines and Culture Conditions* section for details). After this initial incubation, cells were washed twice with DPBS, and the media was replaced with 6 mL of FBS-free DMEM and Ham’s F-12 50/50 media (Corning, catalog no. 10-092-CV) and incubated for an additional 24 hours. Conditioned media was filtered through a 0.45 µm filter (Fisherbrand™, catalog no. 09-720-005), aliquoted, snap-frozen in liquid nitrogen, and stored at −80°C until analysis. This conditioned media is referred to as “Media” in **Figs. 4C-E** and **Fig. S4C-F**.

For ICP-MS, attached cells were washed twice with ice-cold DPBS, scraped into 1 mL of 80% methanol (Fisher Scientific, catalog no. A456-4) over dry ice, snap-frozen in liquid nitrogen, and stored at −80°C until analysis. For Labile Zinc Quantification, attached cells were washed twice with room-temperature DPBS, scraped into 1 mL of room-temperature DPBS, and immediately processed for staining. Six and three replicates per condition (for both cells and conditioned media) were harvested for ICP-MS and labile zinc quantification, respectively. In both cases, two additional plates were cultured in parallel to estimate the cell number at the end of the experiment, which was used to normalize zinc levels per cell. Cell counts were determined using a hemocytometer, by counting four squares in two independent aliquots taken from the same sample and averaging the results.

### Quantification of metals by mass spectrometry

Human (**Fig. 3F**) and Mouse Plasma and Mouse TIF (**Fig. 3A**) were collected as described in the *Isolation of Plasma and Tumor Interstitial Fluid (TIF)* section. *In vitro* cells and conditioned media were collected as outlined in the *Harvesting Cells and Conditioned Media for ICP-MS and Labile Zinc Quantification* section. For cells collected in 80% methanol, the methanol was evaporated to dryness under nitrogen gas prior to further processing. All samples including methanol-evaporated cells, conditioned media, TIF, and plasma, were spiked with 159Tb and 89Y (High-Purity Standards, catalog nos. 1057-1 and 1067-1, respectively) as ISTDs to final concentrations of 0.2 ppb and 1 ppb, respectively. ISTDs were used to correct variations in sample preparation and to monitor fluctuations during the ICP-MS acquisition. Samples were digested with 250 µL of concentrated nitric acid (Fisher Chemical™, catalog no. A509P500) at 50°C for 16 hours. Following digestion, samples were diluted to a final volume of 5 mL with metal-free water (produced using a Milli-Q system, Millipore Sigma, catalog no. IQ7003). Multi-element metal standard calibration curves were generated at the same time with each batch of samples, using a premixed metal ion stock solution (High-Purity Standards, catalog no. QCS-27-100), using the exact sample procedure as for the samples. The calibration curves used ½ serial dilutions ranging from 250 ppb to 0.12 ppb and were spiked with same amount of ISTD as the samples. Quality controls were prepared by exactly same protocol as the samples and were ran at the beginning and the end of the analysis of each batch, to verify measurement accuracy thorough the batch. The limit of quantification for each metal was determined for each batch, and was usually 0.12 ppb or 0.24 ppb, depending on the batch. All solvents used in the protocol were of LC/MS or TraceMetal grade. All samples, including controls, were prepared in 15 mL polypropylene tubes (Fisher Scientific; catalog no. 14-955-238).

Metal quantification was performed using an iCAP RQ ICP-MS (Thermo Scientific, serial no. ICAPRQ01408) equipped with a Cetac ASX-560 autosampler (Teledyne Cetac Technologies). The ICP-MS system was operated under constant argon plasma gas flow at 110 psi. Elemental detection was performed in kinetic energy discrimination (KED) mode using helium gas at 17 psi. Data for each sample were acquired in triplicate, and averaged values were reported.

### Quantification of metabolite levels

Mouse TIF was collected as described in the *Isolation of Plasma and Tumor Interstitial Fluid (TIF)* section. LC/MS was performed using a QExactive orbitrap mass spectrometer using an Ion Max source and heated electrospray ionization (HESI) probe coupled with a Dionex Ultimate 3000 UPLC system (Thermo Fisher). External mass calibration was set every 7 days and internal mass calibration (lock masses) was not used. The mass spectrometer was operated in full-scan, polarity switching mode with the spray voltage set to 3.0 kV, the heated capillary held at 275 °C, and the HESI probe held at 350 °C. Seath gas flow rate was set to 40 units, the auxiliary gas flow at 15 units, and the sweep gas flow at 1 unit. Data acquisition was performed in a range of 70 to1000 *m/z*, with resolution set to 70,000, the AGC target at 1e6, and the maximum injection time at 20 milliseconds.

A library of 149 chemical standards of plasma polar metabolites was built as we have previously described (see first paragraph of *Quantification of metabolite levels in TIF and plasma* from the methods section in Sullivan MR et al. article^199^). Polar metabolites were extracted from 5 µL of each sample using 45 µL of a solvent mixture consisting of acetonitrile, methanol, and formic acid (75:25:0.1), which included isotopically labeled internal standards (ISTDs) listed in **Table S7**. Samples were vortexed at 4°C for 10 minutes and centrifuged at 15,000g for 10 minutes to separate the supernatant containing soluble polar metabolites. Aliquots of 2 µL of sample were injected onto a ZIC-pHILIC (2.1×150 mm) analytical column equipped with a guard column (2.1 x 20mm), both of 5 µm particle size (EMD Milipore, catalog no. 150454). Column was maintained at 25°C and the autosampler was set at 4°C. The mobile phase consisted of Buffer A (20 mM ammonium carbonate, 0.1% ammonium hydroxide), and Buffer B (acetonitrile). The chromatographic gradient was run at a flow rate of 0.150 mL/minute as follows: 0–20 min: linear gradient from 80% to 20% B; 20–20.5 min: linear gradient from 20% to 80% B; 20.5–28 min: hold at 80%. All solvents used in this protocol were LC/MS gradient.

Metabolite identification was performed using XCalibur 2.2 software (Thermo Fisher Scientific), applying a 5 ppm mass accuracy window and a 0.5-minute retention time window. External standard pools were used for metabolite peak identification based on *m/z* values and retention times. The limit of detection ranged from 100 nM to 3 µM, depending on the metabolite. For metabolite quantification, two methods were used: stable isotope dilution for metabolites with available ISTDs, and external standard calibration for those without Briefly, for stable isotope dilution quantification, peak areas of labeled ISTDs were compared to external standard pools to determine their concentration. The peak area of each unlabeled metabolite in the TIF and plasma samples was then compared to the quantified ISTDs to calculate metabolite concentrations. For metabolites without ISTDs, quantification by external calibration involved normalizing the peak area of each analyte to that of a labeled ISTD to account for sample recovery differences. This normalization was applied to both biological samples and external standard pool dilutions. A standard curve was generated from the normalized peak areas, with linear fits typically above r² = 0.95. These curves were used to convert peak areas in TIF or plasma samples into metabolite concentrations. These results are considered semiquantitative due to matrix differences between external standards (prepared in water) and the samples. Please, refer to *Quantification of metabolite levels in TIF and plasma* from the methods section in Sullivan MR et al. article^199^ for more details.

### CRISPR Drop out Screen

The pooled mouse metabolic-focused CRISPR KO library was a gift from Dr. Kivanc Birsoy (Addgene, catalog no. 160129). Fifteen million Yumm5.2 cells expressing sh*Cdkn2a* or control shGFP were infected with pooled CRISPR KO library to an MOI <0.25 to achieve ∼500-fold library coverage after selection with 3 µg/mL puromycin. The cells were passaged every 2–3 days for two weeks, maintaining the full population, before implantation into mice. A total of 350,000 cells per mouse were injected into 36 mice per group, achieving a 550-fold library coverage. Mice were treated with either anti-Pd1 or IgG as described in the *Mice Treatments* section above. Once tumors reached 0.5 cm³, they were harvested, and DNA was isolated using the Quick-DNA Miniprep Plus kit (Zymo Research, catalog no. D4069) following the manufacturer’s instructions. DNA from each mouse in the same group was pooled and used for gRNA amplification. To ensure adequate gRNA representation, calculations from Wohlhieter et al.^200^ were used to achieve >450-fold coverage. In brief, 64 µg of genomic DNA per condition was divided into 16 PCR reactions, with 4 µg per reaction. The gRNA region was amplified using primers listed in **Table S8** and Ex Taq DNA polymerase (Takara, catalog no. RR001A) following the manufacturer’s protocol. The PCR conditions were as follows: one cycle at 94 °C for 2 minutes, followed by 30 cycles of 94 °C for 20 seconds, 63 °C for 30 seconds, and 72 °C for 45 seconds, concluding with a final extension step at 72 °C for 1 minute. The reaction was performed in a thermal cycler (Bio-Rad, T100™ Thermal Cycler, serial no. 621BR59583). The resulting PCR amplicons were purified using the Wizard® SV Gel and PCR Clean-Up System (Promega, catalog no. A9282) following the manufacturer’s instructions and pooled. The cleaned amplicons were sequenced using a NextSeq high output 75 cycle kit 1×50 in an Illumina NextSeq 500 (Serial no. NB: 501446). Data was analyzed using the MAGeCK bioinformatics pipeline to identify negatively enriched genes (Li et al., 2014). **Fig. 3C** shows the Log_2_ fold change of the negative CRISPR score of solute carrier transporter in YUMM5.2 sh*Cdkn2a* tumors in mice treated with anti-Pd-1 vs. IgG vs. shControl tumors in mice treated with anti-Pd-1 vs. IgG.

### Methyl-β-cyclodextrin (MβCD) treatment

Cells seeded in 12-well plates (Corning, catalog no. 3513) for immunofluorescence (IF) or in 6 cm plates (Falcon, catalog no. 353002) for flow cytometry were treated with 10 mM MβCD (Sigma Aldrich, catalog no. C4555-10G) diluted in FBS-free DMEM and Ham’s F-12 50/50 media (Corning, catalog no. 10-092-CV), supplemented with 10 mM HEPES (Gibco, catalog no. 15630080). The treatment was carried out for 1 hour at 37°C with 5% CO₂. After treatment, cells were collected for downstream analysis (i.e. IF or labile zinc quantification). Note that MβCD-treated cells are extremely loosely adhered, so washes must be done gently. To prevent cell loss during the immunofluorescence staining procedure, fixation time was increased to 20 minutes.

### Statin treatment

Two million cells were seeded in 10 cm dishes and treated with 1 µM Lovastatin (Selleckchem, Cat. No. S2061), Atorvastatin (MedChemExpress, Cat. No. HY-B0589) or vehicle (DMSO). Cells were incubated for 4 days, with media replaced at 48 hours using fresh media containing corresponding treatment. At 72 hours, cells were trypsinized, and 35,000 cells per condition were seeded onto coverslips in 12-well plates for immunofluorescence, using media containing corresponding treatment. In parallel, 1 million cells were seeded in 10 cm dishes in media containing corresponding treatment for intracellular labile zinc quantification. After 24 hours, cells on coverslips were washed, fixed, and stained (see *Immunofluorescence* section). Cells in 10 cm dishes were washed, scraped, and stained for labile zinc quantification (see *In Vitro Labile Zinc Quantification* section).

### *In vitro* labile Zinc quantification

Cells and conditioned media were collected as outlined in the *Harvesting Cells and Conditioned Media for ICP-MS and Labile Zinc Quantification* section. One million scraped cells were pelleted in a round-bottom 96-well plate (Corning, catalog no. 3366) by centrifugation at 350 g for 5 minutes at 4°C. Cells were then stained with 50 µM Zinpyr-1 (Cayman Chemical, catalog no. 15122) diluted in DPBS, containing a 1:1000 dilution of Live/Dead Fixable Near-IR Dead Cell Stain kit (Invitrogen, catalog no. L34976), in a final volume of 50 µL for 20 minutes at 4°C, protected from light. Following staining, cells were fixed in 4% paraformaldehyde (Sigma Aldrich, catalog no. 158127-3Kg) for 15 minutes at 4°C, shielded from light, and resuspended in FACS buffer (1x PBS, 10 mM HEPES [Gibco, catalog no. 15630080], 1% heat-inactivated FBS [Sigma Aldrich, catalog no. F4135-100ML], 0.05% sodium azide [Sigma Aldrich, catalog no. S2002-25G]). Stained cells were immediately analyzed on a CytoFlex Flow Cytometer (Beckman Coulter, serial no. 822338373). Flow cytometry data were analyzed with FlowJo version 10. The median fluorescence intensity (MFI) of single live cells was measured to assess intracellular zinc levels per sample.

Labile zinc concentration was measured as previously described^201^. Briefly, 50 µL of conditioned media was mixed with 50 µL of 50 µM Zinpyr-1 (Cayman Chemical, catalog no. 15122) in a 96-well polystyrene microplate (Greiner Bio-One, catalog no. 655101). The mixture was incubated for 20 minutes at 37°C in a CO₂ incubator. Fluorescence was then measured in a fluorescent plate reader (Promega GloMax explorer, serial no. 9720100141) with excitation at 475 nm and emission at 500-550 nm. This first fluorescence measurement reflects the signal of Zinpyr-1 bound to zinc in the sample (F). To determine the baseline fluorescence, 50 µL of 600 µM of the zinc chelator N,N,N’,N’-tetrakis(2-pyridinylmethyl)-1,2-ethanediamine (TPEN, MilliporeSigma, catalog no. 6163941100MG), was added to each well. After another 20-minute incubation at 37°C in a CO₂ incubator, fluorescence was measured again, representing the minimum fluorescence (F_min_) due to the chelation of zinc. Finally, to obtain the maximum fluorescence (F_max_), 50 µL of 2 mM ZnSO₄ (Sigma Aldrich, catalog no. Z2876-500ML) was added to each well. After a final 20-minute incubation at 37°C in a CO₂ incubator, the fluorescence was measured one last time. The zinc concentration in each sample was then calculated using the formula: Zn²⁺ concentration = K_D_ × ((F – Fmin) / (Fmax – F)), where K_D_ is the dissociation constant of the zinc:Zinpyr-1 complex (0.7 nM)^201^. Finally, zinc concentrations were normalized to the number of cells in each sample at the time of harvesting.

### Immunofluorescence (IF)

Approximately 30,000 to 50,000 cells were seeded onto coverslips in 12-well plates (Corning, catalog no. 3513) and incubated at 37°C with 5% CO₂. Because Yumm1.7 cells have low attachment properties, these cells were seeded onto poly-L-ornithine-coated coverslips. Briefly, coverslips were first washed with 2 mL of 200 proof ethanol (Decon Labs) for 5 minutes, followed by two washes with molecular-grade water (Corning, catalog no. 46000CM). Then, 500 µL of poly-L-ornithine (Sigma Aldrich, catalog no. P4957-50mL) was added to each well containing a coverslip. The plates were sealed with parafilm (VWR, catalog no. 52858-032) to prevent evaporation, and incubated overnight at 37°C. Before seeding the cells, the coverslips were washed three times with 1 mL of molecular-grade water and air-dried for 10 minutes. Approximately 16 hours after seeding, the cells were fixed with 4% paraformaldehyde (Sigma Aldrich, catalog no. 158127-3Kg) for 10 minutes followed by permeabilized with 0.2% Triton X-100 (VWR, catalog no. 97062-208) for 5 minutes. For plasma membrane staining of SLC39A9, however, permeabilization was omitted. The cells were then blocked with 5 µg/mL of Cd16/Cd32 (Invitrogen, catalog no. 14016185) diluted in 3% BSA/PBS (Fisher BioReagents, catalog no. BP1600) for 10 minutes. Both human and mouse cells were subsequently incubated with 10 µg/mL of anti-Slc39a9 (Thermo Scientific, catalog no. PA521074), diluted in 3% BSA/PBS for 1 hour. After incubation, the cells were washed three times with PBS and then incubated with 0.75 µg/mL of secondary anti-rabbit Cy3-conjugated antibody (Jackson ImmunoResearch, catalog no. 111-165-003), diluted in 3% BSA/PBS for 1 hour in the dark. Cells were then washed 10 times with PBS and permeabilized with 0.2% Triton X-100 for 5 minutes. Cells were stained with 0.15 μg/mL DAPI (diluted in PBS) for 1 minute. Finally, coverslips were mounted (Sigma Aldrich, catalog no. F6182-20ML) and sealed with transparent nail polish. Controls included staining with 10 µg/mL normal rabbit IgG (Cell Signaling, catalog no. 2729) and secondary-only staining.

For filipin staining cells were cultured on coverslips and fixed with 4% paraformaldehyde for 10 minutes. Cells were incubated with 0.1 mg/mL filipin III (Cayman Chemicals, catalog no. 70440) for 1h at room temperature. Coverslips were mounted and sealed with transparent nail polish. Of note that Yumm1.7 cells were seeded in media containing lipiddepleted FBS (Omega Scientific, catalog no. FB-50).

Five to ten images were captured per condition using an inverted microscope (Nikon ECLIPSE Ti2, serial no. 542740) equipped with a 20x/0.17 objective (Nikon DIC N2 Plan Apo), a digital camera (ORCA-Fusion, catalog no. C1440-20UP, serial no. 000670) and the NIS-Elements AR software. Image analysis was conducted using Python (v. 3.10).

SLC39A9 quantification was performed as follows: Nuclear staining images (DAPI) were normalized to the 3rd and 97th percentiles before segmentation, which was carried out using a pre-trained CellPose model (“cyto”)^202^. Nuclei touching the image borders were excluded from further analysis. The mean Cy3 fluorescence signal (anti-SLC39A9) was quantified within the nuclear regions. Background signal was corrected by subtracting the median intensity of the entire image. Finally, the number of cells per condition was randomized, and 200 cells per condition were plotted as fold change relative to the control condition. Filipin quantification was performed as follows: Individual cells were defined as regions of interest (ROIs) and captured using image thresholding. Analysis was conducted using the ‘Analyze Particles’ tool in ImageJ (Fiji), excluding cells at the image periphery. Integrated Density values were determined for each ROI. A total of at least 100 cells per condition were analyzed, and data are presented as fold change relative to the control condition.

### Western Blotting

Cell lysates were collected in 1X sample buffer [2% SDS (Millipore Sigma, catalog no. 75746-1Kg), 10% glycerol (Sigma Aldrich, catalog no. G5516), 0.01% bromophenol blue (Millipore Sigma, catalog no. BX1410), 62.5 mM Tris (Fisher Scientific, catalog no. AAJ65594A1), pH=6.8 and 0.1 M DTT (Thermo Fisher, catalog no. 15508-013)] and boiled at 95° C for 10 minutes. Protein concentration was determined using Bradford assay (Bio-Rad, catalog no. 5000006) following manufacturer’s instructions. Proteins were resolved using SDS-PAGE gels and transferred to nitrocellulose membranes (GE Healthcare Life Sciences, catalog no. 10600001) as previously described^54^. Antibodies used include: anti human p16 (Abcam, catalog no. ab108349, working concentration 1.5 µg/mL), mouse anti-p16 (Cell Signaling Technology, catalog no. 29271S, working concentration 1 µg/mL), anti-p19 (Abcam, catalog no. ab80, working concentration 1 µg/mL), anti-β-Actin (Sigma-Aldrich, catalog no. A1978, working concentration 1 µg/mL), anti-mouse HRP (Cell Signaling Technology, catalog no. 7076, working concentration 0.1 µg/mL), and anti-rabbit HRP (Cell Signaling Technology, catalog no. 7074,, working concentration 0.2 µg/mL).

### Bulk RNA-Seq and RT-qPCR

Total RNA was extracted from cells with Trizol (Ambion, catalog no. 15596018) and DNase treated, cleaned, and concentrated using Zymo columns (Zymo Research, catalog no. R1013) following the manufacturer’s instructions. Optical density values for RNA were measured using NanoDrop One (Thermo Scientific) to confirm an A260 and A280 ratios above 1.9. For bulk RNA-Seq, RNA integrity number (RIN) was measured using BioAnalyzer (Agilent Technologies; RRID:SCR_019715) RNA 6000 Nano Kit to confirm RIN above 7 for each sample. The cDNA libraries, next-generation sequencing, and bio-informatics analysis was performed by Novogene. Differential expression analysis was done using DESeq (v2_1.6.3). Raw and processed RNA-seq data can be found on Gene Expression Omnibus (GSE295787).

For RT-qPCR, a total of 2 µg of RNA was reverse-transcribed to cDNA (Applied Biosystems, catalog no. A25742) following manufacturer’s instructions in a thermal cycler (Bio-Rad, T100™ Thermal Cycler, serial no. 621BR59583). Relative expression of target genes was analyzed using the Real-Time PCR System (Bio-Rad, CFX Connect, serial no. 788BR09362) with clear 96-well plates (Greiner Bio-One, catalog no. 652240). Primers were designed using the Integrated DNA Technologies (IDT) web tool (**Table S8**). Conditions for amplification were: 5 min at 95° C, 40 cycles of 10 sec at 95° C and 7 sec at 62° C. The assay ended with a melting curve program: 15 sec at 95° C, 1 min at 70° C, then ramping to 95° C while continuously monitoring fluorescence. Each sample was assessed in triplicate. Relative quantification was determined to multiple reference genes (Rplp0, Gusb, and Rp132) to ensure reproducibility using the delta-delta CT method.

### Senescence and proliferation assays

SA-β-Gal staining was performed as previously described^203^. Cells were fixed in a mixture of 2% formaldehyde (Fisher Scientific, catalog no. BP531-500) and 0.2% glutaraldehyde (Fisher Scientific, catalog no. 01909-100) diluted in PBS for 5 minutes, followed by staining with a staining cocktail containing: 40 mM Na_2_HPO_4_ (Sigma Aldrich, catalog no. 567547), 150 mM NaCl (Sigma Aldrich, catalog no. S5886), 2 mM MgCl_2_ (Sigma Aldrich, catalog no. M1028), 5Mm K_3_Fe(CN)_6_ (Sigma Aldrich, catalog no. 244023), 5 mM K_4_Fe(CN)_6_ (Sigma Aldrich, catalog no. 455989), and 1 mg/ml 5-Bromo-4-chloro-3-indolyl β-D-galactopyranoside (Sigma Aldrich, catalog no. B4252). Staining was incubated overnight at 37° C in a non-CO2 incubator after which reaction was stopped with tap water. Images were acquired immediately upon reaction stopped using an inverted microscope (Nikon Eclipse Ts2, serial no. 139045) with a 20X/0.40 objective (Nikon LWD) equipped with a camera (Nikon DS-Fi3). Each sample was assessed in triplicate and at least 100 cells per well were counted (>300 cells per experiment). For colony formation, an equal number of cells were seeded in 12-well plates and cultured for an additional 1-2 weeks. Colony formation was visualized by fixing cells in 1% paraformaldehyde for 5 minutes followed by staining with 0.05% crystal violet for 20 minutes on rocking. Wells were destained in 500mL 10% acetic acid for 10 minutes on rocking and absorbance (590 nm) measured using a spectrophotometer (Promega GloMax explorer, serial no. 9720100141). Each sample was assessed in triplicate.

### Qualification and Statistical analysis

GraphPad Prism version 10.0 was used to perform statistical analysis. The level of significance between two groups was assessed with pared or unpaired Student’s *t* test as necessary and indicated in figure caption. For data sets with more than two groups, one-way ANOVA followed by Šidák correction for multiple comparisons was applied. For longitudinal tumor growth studies mixed-effects model was used and Šidák correction for multiple comparisons was applied. For survival analysis log-rank (Mantel-Cox) test was used. Pearson correlation was used to assess correlation. P-values <0.05 were considered significant. Specific statistical details used in scRNA-Seq and bulk RNA-Seq data are included in the corresponding section.

## Notes

### Competing Interest Statement

The authors have declared no competing interest.

### Summary of Updates

1) The title has been updated to better reflect the main findings of the study. 2) The manuscript has been reorganized to improve the logical flow of the text and data presentation. 3) A new abstract has been written to align with the revised structure and to clearly highlight the key results. 4) All figures have been redesigned to make the data easier to interpret. 5) A new *in vitro* experiment show that anti-Pd1 treatment enhances phagocytosis by RAW264.7 macrophages when co-cultured with *Cdkn2a* ^High^ cancer cells. This phagocytic response is impaired when macrophages are co-cultured with *Cdkn2a* ^Low^ cancer cells. 6) New *in vitro* data demonstrating that inhibition of *de novo* cholesterol synthesis with statins decreases plasma membrane Slc39a9 and reduces intracellular labile zinc levels in *Cdkn2a* ^Low^ cancer cells. 7) New *in vivo* experiments using a second mouse model demonstrate that *Cdkn2a* ^Low^ cancer cells outcompete macrophages for zinc via plasma membrane Slc39a9. This zinc depletion leads to reduced macrophage phagocytosis. 8) A new *in vivo* experiment in which zinc-uptake proficient RAW264.7 macrophages (overexpressing Slc39a1) were co-injected with *Cdkn2a* ^Low^ cancer cells into NSG mice and treated with either anti-Pd1 or IgG. The results show that zinc-replenished macrophages restore phagocytic function in response to anti-Pd1 and lead to a reduction in tumor volume.

## REFERENCES

1 Han, G. et al. 9p21 loss confers a cold tumor immune microenvironment and primary resistance to immune checkpoint therapy. Nature communications 12, 5606 (2021). 10.1038/s41467-021-25894-9

2 Tu, Z. et al. The cell senescence regulator p16 is a promising cancer prognostic and immune check-point inhibitor (ICI) therapy biomarker. Aging (Albany NY) 15, 2136–2157 (2023). 10.18632/aging.204601

3 Deng, C. et al. Pan-cancer analysis of CDKN2A alterations identifies a subset of gastric cancer with a cold tumor immune microenvironment. Hum Genomics 18, 55 (2024). 10.1186/s40246-024-00615-7

4 Fan, K., Dong, Y., Li, T. & Li, Y. Cuproptosis-associated CDKN2A is targeted by plicamycin to regulate the microenvironment in patients with head and neck squamous cell carcinoma. Front Genet 13, 1036408 (2022). 10.3389/fgene.2022.1036408

5 Luo, J. P., Wang, J. & Huang, J. H. CDKN2A is a prognostic biomarker and correlated with immune infiltrates in hepatocellular carcinoma. Biosci Rep 41 (2021). 10.1042/BSR20211103

6 Chen, Z. et al. Comprehensive Analysis Revealed that CDKN2A is a Biomarker for Immune Infiltrates in Multiple Cancers. Front Cell Dev Biol 9, 808208 (2021). 10.3389/fcell.2021.808208

7 Gutiontov, S. I. et al. CDKN2A loss-of-function predicts immunotherapy resistance in non-small cell lung cancer. Scientific reports 11, 20059 (2021). 10.1038/s41598-021-99524-1

8 Yu, J. et al. Genetic Aberrations in the CDK4 Pathway Are Associated with Innate Resistance to PD-1 Blockade in Chinese Patients with Non-Cutaneous Melanoma. Clin Cancer Res 25, 6511–6523 (2019). 10.1158/1078-0432.CCR-19-0475

9 Adib, E. et al. CDKN2A Alterations and Response to Immunotherapy in Solid Tumors. Clin Cancer Res 27, 4025–4035 (2021). 10.1158/1078-0432.CCR-21-0575

10 Liu, S. et al. Response and recurrence correlates in individuals treated with neoadjuvant anti-PD-1 therapy for resectable oral cavity squamous cell carcinoma. Cell Rep Med 2, 100411 (2021). 10.1016/j.xcrm.2021.100411

11 Peng, Y. et al. Co-occurrence of CDKN2A/B and IFN-I homozygous deletions correlates with an immunosuppressive phenotype and poor prognosis in lung adenocarcinoma. Mol Oncol 16, 1746–1760 (2022). 10.1002/1878-0261.13206

12 Zhao, L., Zhou, X., Li, H., Yin, T. & Jiang, Y. Prognosis of immunotherapy for non-small cell lung cancer with CDKN2A loss of function. J Thorac Dis 16, 507–515 (2024). 10.21037/jtd-23-1017

13 Leon, K. E., Tangudu, N. K., Aird, K. M. & Buj, R. Loss of p16: A Bouncer of the Immunological Surveillance? Life (Basel) 11 (2021). 10.3390/life11040309

14 Banchereau, R. et al. Molecular determinants of response to PD-L1 blockade across tumor types. Nat Commun 12, 3969 (2021). 10.1038/s41467-021-24112-w

15 Wang, S. et al. Loss of CDKN2A Enhances the Efficacy of Immunotherapy in EGFR Mutant Non-Small Cell Lung Cancer. Cancer Res (2024). 10.1158/0008-5472.CAN-24-1817

16 Xue, L. et al. Next-generation sequencing identifies CDKN2A alterations as prognostic biomarkers in recurrent or metastatic head and neck squamous cell carcinoma predominantly receiving immune checkpoint inhibitors. Front Oncol 13, 1276009 (2023). 10.3389/fonc.2023.1276009

17 Jang, H. J. et al. Inhibition of Cyclin Dependent Kinase 4/6 Overcomes Primary Resistance to Programmed Cell Death 1 Blockade in Malignant Mesothelioma. Ann Thorac Surg 114, 1842–1852 (2022). 10.1016/j.athoracsur.2021.08.054

18 Buj, R., Leon, K. E., Anguelov, M. A. & Aird, K. M. Suppression of p16 alleviates the senescence-associated secretory phenotype. Aging (Albany NY) 13, 3290–3312 (2021). 10.18632/aging.202640

19 Brahmer, J. R. et al. Safety and activity of anti-PD-L1 antibody in patients with advanced cancer. N Engl J Med 366, 2455–2465 (2012). 10.1056/NEJMoa1200694

20 Topalian, S. L. et al. Safety, activity, and immune correlates of anti-PD-1 antibody in cancer. N Engl J Med 366, 2443–2454 (2012). 10.1056/NEJMoa1200690

21 Wolchok, J. D. et al. Final, 10-Year Outcomes with Nivolumab plus Ipilimumab in Advanced Melanoma. N Engl J Med (2024). 10.1056/NEJMoa2407417

22 Finn, R. S. et al. Atezolizumab plus Bevacizumab in Unresectable Hepatocellular Carcinoma. N Engl J Med 382, 1894–1905 (2020). 10.1056/NEJMoa1915745

23 Herbst, R. S. et al. Atezolizumab for First-Line Treatment of PD-L1-Selected Patients with NSCLC. N Engl J Med 383, 1328–1339 (2020). 10.1056/NEJMoa1917346

24 Pardoll, D. M. The blockade of immune checkpoints in cancer immunotherapy. Nat Rev Cancer 12, 252–264 (2012). 10.1038/nrc3239

25 Moore, J. et al. Identification of a conserved subset of cold tumors responsive to immune checkpoint blockade. J Immunother Cancer 13 (2025). 10.1136/jitc-2024-010528

26 Rappold, P. M. et al. A Targetable Myeloid Inflammatory State Governs Disease Recurrence in Clear-Cell Renal Cell Carcinoma. Cancer Discov 12, 2308–2329 (2022). 10.1158/2159-8290.CD-21-0925

27 Rashidian, M. et al. Immuno-PET identifies the myeloid compartment as a key contributor to the outcome of the antitumor response under PD-1 blockade. Proc Natl Acad Sci U S A 116, 16971–16980 (2019). 10.1073/pnas.1905005116

28 Bader, J. E. et al. Obesity induces PD-1 on macrophages to suppress anti-tumour immunity. Nature 630, 968–975 (2024). 10.1038/s41586-024-07529-3

29 Gordon, S. R. et al. PD-1 expression by tumour-associated macrophages inhibits phagocytosis and tumour immunity. Nature 545, 495–499 (2017). 10.1038/nature22396

30 Coulton, A. et al. Using a pan-cancer atlas to investigate tumour associated macrophages as regulators of immunotherapy response. Nat Commun 15, 5665 (2024). 10.1038/s41467-024-49885-8

31 Strauss, L. et al. Targeted deletion of PD-1 in myeloid cells induces antitumor immunity. Sci Immunol 5 (2020). 10.1126/sciimmunol.aay1863

32 Kim, I. S. et al. Immuno-subtyping of breast cancer reveals distinct myeloid cell profiles and immunotherapy resistance mechanisms. Nature cell biology 21, 1113–1126 (2019). 10.1038/s41556-019-0373-7

33 Yu, J. et al. Liver metastasis restrains immunotherapy efficacy via macrophage-mediated T cell elimination. Nature medicine 27, 152–164 (2021). 10.1038/s41591-020-1131-x

34 Zhang, Y. et al. Myeloid cells are required for PD-1/PD-L1 checkpoint activation and the establishment of an immunosuppressive environment in pancreatic cancer. Gut 66, 124–136 (2017). 10.1136/gutjnl-2016-312078

35 Davar, D. et al. Neoadjuvant vidutolimod and nivolumab in high-risk resectable melanoma: A prospective phase II trial. Cancer Cell 42, 1898–1918 e1812 (2024). 10.1016/j.ccell.2024.10.007

36 Shen, L. et al. PD-1/PD-L pathway inhibits M.tb-specific CD4(+) T-cell functions and phagocytosis of macrophages in active tuberculosis. Sci Rep 6, 38362 (2016). 10.1038/srep38362

37 Wang, L. et al. PD-L1-expressing tumor-associated macrophages are immunostimulatory and associate with good clinical outcome in human breast cancer. Cell Rep Med 5, 101420 (2024). 10.1016/j.xcrm.2024.101420

38 Li, C. W., Lai, Y. J., Hsu, J. L. & Hung, M. C. Activation of phagocytosis by immune checkpoint blockade. Front Med 12, 473–480 (2018). 10.1007/s11684-018-0657-5

39 Liu, M. et al. Metabolic rewiring of macrophages by CpG potentiates clearance of cancer cells and overcomes tumor-expressed CD47-mediated ‘don’t-eat-me’ signal. Nat Immunol 20, 265–275 (2019). 10.1038/s41590-018-0292-y

40 Ring, N. G. et al. Anti-SIRPalpha antibody immunotherapy enhances neutrophil and macrophage antitumor activity. Proc Natl Acad Sci U S A 114, E10578–E10585 (2017). 10.1073/pnas.1710877114

41 Gonzalez, M. A. et al. Phagocytosis increases an oxidative metabolic and immune suppressive signature in tumor macrophages. J Exp Med 220 (2023). 10.1084/jem.20221472

42 Kamb, A. et al. A cell cycle regulator potentially involved in genesis of many tumor types. Science 264, 436–440 (1994). 10.1126/science.8153634

43 Nobori, T. et al. Deletions of the cyclin-dependent kinase-4 inhibitor gene in multiple human cancers. Nature 368, 753–756 (1994). 10.1038/368753a0

44 Sherr, C. J. Cancer cell cycles. Science 274, 1672–1677 (1996). 10.1126/science.274.5293.1672

45 Hall, M. & Peters, G. Genetic alterations of cyclins, cyclin-dependent kinases, and Cdk inhibitors in human cancer. Adv Cancer Res 68, 67–108 (1996). 10.1016/s0065-230x(08)60352-8

46 Cairns, P. et al. Rates of p16 (MTS1) mutations in primary tumors with 9p loss. Science 265, 415–417 (1994). 10.1126/science.8023167

47 Herman, J. G. et al. Inactivation of the CDKN2/p16/MTS1 gene is frequently associated with aberrant DNA methylation in all common human cancers. Cancer Res 55, 4525–4530 (1995).

48 Ranade, K. et al. Mutations associated with familial melanoma impair p16INK4 function. Nat Genet 10, 114–116 (1995). 10.1038/ng0595-114

49 Zhao, R., Choi, B. Y., Lee, M. H., Bode, A. M. & Dong, Z. Implications of Genetic and Epigenetic Alterations of CDKN2A (p16(INK4a)) in Cancer. EBioMedicine 8, 30–39 (2016). 10.1016/j.ebiom.2016.04.017

50 Sherr, C. J. The INK4a/ARF network in tumour suppression. Nat Rev Mol Cell Biol 2, 731–737 (2001). 10.1038/35096061

51 Serrano, M., Hannon, G. J. & Beach, D. A new regulatory motif in cell-cycle control causing specific inhibition of cyclin D/CDK4. Nature 366, 704–707 (1993). 10.1038/366704a0 [doi]

52 Serrano, M., Lin, A. W., McCurrach, M. E., Beach, D. & Lowe, S. W. Oncogenic ras provokes premature cell senescence associated with accumulation of p53 and p16INK4a. Cell 88, 593–602 (1997). 10.1016/s0092-8674(00)81902-9

53 Buj, R. & Aird, K. M. p16: cycling off the beaten path. Mol Cell Oncol 6, e1677140 (2019). 10.1080/23723556.2019.1677140

54 Buj, R. et al. Suppression of p16 Induces mTORC1-Mediated Nucleotide Metabolic Reprogramming. Cell Rep 28, 1971–1980 e1978 (2019). 10.1016/j.celrep.2019.07.084

55 Tangudu, N. K. et al. De Novo Purine Metabolism is a Metabolic Vulnerability of Cancers with Low p16 Expression. Cancer Res Commun 4, 1174–1188 (2024). 10.1158/2767-9764.Crc-23-0450

56 Tangudu, N. K. et al. ATR promotes mTORC1 activity via de novo cholesterol synthesis. EMBO Rep (2025). 10.1038/s44319-025-00451-3

57 Wolff, B. & Naumann, M. INK4 cell cycle inhibitors direct transcriptional inactivation of NF-kappaB. Oncogene 18, 2663–2666 (1999). 10.1038/sj.onc.1202617

58 Tyagi, E. et al. Loss of p16(INK4A) stimulates aberrant mitochondrial biogenesis through a CDK4/Rb-independent pathway. Oncotarget 8, 55848–55862 (2017). 10.18632/oncotarget.19862

59 Jenkins, N. C. et al. The p16(INK4A) tumor suppressor regulates cellular oxidative stress. Oncogene 30, 265–274 (2011). 10.1038/onc.2010.419

60 Choi, B. Y. et al. The tumor suppressor p16(INK4a) prevents cell transformation through inhibition of c-Jun phosphorylation and AP-1 activity. Nat Struct Mol Biol 12, 699–707 (2005). 10.1038/nsmb960

61 Lee, M. H. et al. Tumor suppressor p16(INK4a) inhibits cancer cell growth by downregulating eEF1A2 through a direct interaction. J Cell Sci 126, 1744–1752 (2013). 10.1242/jcs.113613

62 Al-Khalaf, H. H., Mohideen, P., Nallar, S. C., Kalvakolanu, D. V. & Aboussekhra, A. The cyclin-dependent kinase inhibitor p16INK4a physically interacts with transcription factor Sp1 and cyclin-dependent kinase 4 to transactivate microRNA-141 and microRNA-146b-5p spontaneously and in response to ultraviolet light-induced DNA damage. J Biol Chem 288, 35511–35525 (2013). 10.1074/jbc.M113.512640

63 Murakami, Y., Mizoguchi, F., Saito, T., Miyasaka, N. & Kohsaka, H. p16(INK4a) exerts an anti-inflammatory effect through accelerated IRAK1 degradation in macrophages. J Immunol 189, 5066–5072 (2012). 10.4049/jimmunol.1103156

64 Hu, Y. et al. Dual roles of hexokinase 2 in shaping microglial function by gating glycolytic flux and mitochondrial activity. Nat Metab 4, 1756–1774 (2022). 10.1038/s42255-022-00707-5

65 Davies, L. C. et al. Peritoneal tissue-resident macrophages are metabolically poised to engage microbes using tissue-niche fuels. Nat Commun 8, 2074 (2017). 10.1038/s41467-017-02092-0

66 Sugimoto, R. et al. Zinc Deficiency as a General Feature of Cancer: a Review of the Literature. Biol Trace Elem Res 202, 1937–1947 (2024). 10.1007/s12011-023-03818-6

67 Kiouri, D. P. et al. Multifunctional role of zinc in human health: an update. EXCLI J 22, 809–827 (2023). 10.17179/excli2023-6335

68 Mehta, A. J., Yeligar, S. M., Elon, L., Brown, L. A. & Guidot, D. M. Alcoholism causes alveolar macrophage zinc deficiency and immune dysfunction. Am J Respir Crit Care Med 188, 716–723 (2013). 10.1164/rccm.201301-0061OC

69 Wirth, J. J., Fraker, P. J. & Kierszenbaum, F. Zinc requirement for macrophage function: effect of zinc deficiency on uptake and killing of a protozoan parasite. Immunology 68, 114–119 (1989).

70 Sheikh, A. et al. Zinc influences innate immune responses in children with enterotoxigenic Escherichia coli-induced diarrhea. J Nutr 140, 1049–1056 (2010). 10.3945/jn.109.111492

71 Hall, S. C. et al. Critical Role of Zinc Transporter (ZIP8) in Myeloid Innate Immune Cell Function and the Host Response against Bacterial Pneumonia. J Immunol 207, 1357–1370 (2021). 10.4049/jimmunol.2001395

72 Kitamura, H. et al. Toll-like receptor-mediated regulation of zinc homeostasis influences dendritic cell function. Nat Immunol 7, 971–977 (2006). 10.1038/ni1373

73 Mocchegiani, E. et al. Metallothioneins/PARP-1/IL-6 interplay on natural killer cell activity in elderly: parallelism with nonagenarians and old infected humans. Effect of zinc supply. Mech Ageing Dev 124, 459–468 (2003). 10.1016/s0047-6374(03)00023-x

74 Aydemir, T. B., Blanchard, R. K. & Cousins, R. J. Zinc supplementation of young men alters metallothionein, zinc transporter, and cytokine gene expression in leukocyte populations. Proceedings of the National Academy of Sciences of the United States of America 103, 1699–1704 (2006). 10.1073/pnas.0510407103

75 Shi, H. N., Scott, M. E., Stevenson, M. M. & Koski, K. G. Energy restriction and zinc deficiency impair the functions of murine T cells and antigen-presenting cells during gastrointestinal nematode infection. J Nutr 128, 20–27 (1998). 10.1093/jn/128.1.20

76 Jeong, J. & Eide, D. J. The SLC39 family of zinc transporters. Mol Aspects Med 34, 612–619 (2013). 10.1016/j.mam.2012.05.011

77 Huang, L. & Tepaamorndech, S. The SLC30 family of zinc transporters - a review of current understanding of their biological and pathophysiological roles. Mol Aspects Med 34, 548–560 (2013). 10.1016/j.mam.2012.05.008

78 Ren, X. et al. SLC39A10 promotes malignant phenotypes of gastric cancer cells by activating the CK2-mediated MAPK/ERK and PI3K/AKT pathways. Exp Mol Med 55, 1757–1769 (2023). 10.1038/s12276-023-01062-5

79 Li, M. et al. Aberrant expression of zinc transporter ZIP4 (SLC39A4) significantly contributes to human pancreatic cancer pathogenesis and progression. Proc Natl Acad Sci U S A 104, 18636–18641 (2007). 10.1073/pnas.0709307104

80 Cheng, X. et al. Zinc transporter SLC39A13/ZIP13 facilitates the metastasis of human ovarian cancer cells via activating Src/FAK signaling pathway. J Exp Clin Cancer Res 40, 199 (2021). 10.1186/s13046-021-01999-3

81 Hogstrand, C., Kille, P., Ackland, M. L., Hiscox, S. & Taylor, K. M. A mechanism for epithelial-mesenchymal transition and anoikis resistance in breast cancer triggered by zinc channel ZIP6 and STAT3 (signal transducer and activator of transcription 3). Biochem J 455, 229–237 (2013). 10.1042/BJ20130483

82 Cheng, X. et al. Solute Carrier Family 39 Member 6 Gene Promotes Aggressiveness of Esophageal Carcinoma Cells by Increasing Intracellular Levels of Zinc, Activating Phosphatidylinositol 3-Kinase Signaling, and Up-regulating Genes That Regulate Metastasis. Gastroenterology 152, 1985–1997 e1912 (2017). 10.1053/j.gastro.2017.02.006

83 Liu, M. et al. ZIP4 Promotes Pancreatic Cancer Progression by Repressing ZO-1 and Claudin-1 through a ZEB1-Dependent Transcriptional Mechanism. Clin Cancer Res 24, 3186–3196 (2018). 10.1158/1078-0432.CCR-18-0263

84 Thomas, P., Pang, Y. & Dong, J. Ligand-independent signaling and migration of breast cancer cells expressing membrane androgen receptor, ZIP9 (SLC39A9). Mol Cell Endocrinol 578, 112060 (2023). 10.1016/j.mce.2023.112060

85 Prasad, A. S. et al. Nutritional and zinc status of head and neck cancer patients: an interpretive review. J Am Coll Nutr 17, 409–418 (1998). 10.1080/07315724.1998.10718787

86 Federico, A. et al. Effects of selenium and zinc supplementation on nutritional status in patients with cancer of digestive tract. Eur J Clin Nutr 55, 293–297 (2001). 10.1038/sj.ejcn.1601157

87 Xie, Y. et al. Higher serum zinc levels may reduce the risk of cervical cancer in Asian women: A meta-analysis. J Int Med Res 46, 4898–4906 (2018). 10.1177/0300060518805600

88 Lin, S. & Yang, H. Ovarian cancer risk according to circulating zinc and copper concentrations: A meta-analysis and Mendelian randomization study. Clin Nutr 40, 2464–2468 (2021). 10.1016/j.clnu.2020.10.011

89 Wu, X., Wu, H., Liu, L., Qiang, G. & Zhu, J. Serum zinc level and tissue ZIP4 expression are related to the prognosis of patients with stages I-III colon cancer. Transl Cancer Res 9, 5585–5594 (2020). 10.21037/tcr-20-2571

90 Zhao, J. et al. Comparative study of serum zinc concentrations in benign and malignant prostate disease: A Systematic Review and Meta-Analysis. Sci Rep 6, 25778 (2016). 10.1038/srep25778

91 Singh, K. P. et al. Effect of zinc on immune functions and host resistance against infection and tumor challenge. Immunopharmacol Immunotoxicol 14, 813–840 (1992). 10.3109/08923979209009237

92 Satoh, M. et al. Prevention of carcinogenicity of anticancer drugs by metallothionein induction. Cancer Res 53, 4767–4768 (1993).

93 Sun, J. et al. Reduction in squamous cell carcinomas in mouse skin by dietary zinc supplementation. Cancer Med 5, 2032–2042 (2016). 10.1002/cam4.768

94 Fong, L. Y., Mancini, R., Nakagawa, H., Rustgi, A. K. & Huebner, K. Combined cyclin D1 overexpression and zinc deficiency disrupts cell cycle and accelerates mouse forestomach carcinogenesis. Cancer Res 63, 4244–4252 (2003).

95 Wang, J. et al. Serum zinc as a biomarker to predict the efficacy of immune checkpoint inhibitors in cancers. PloS one 20, e0326057 (2025). 10.1371/journal.pone.0326057

96 Tang, W. et al. Efficacy of zinc carnosine in the treatment of colorectal cancer and its potential in combination with immunotherapy in vivo. Aging (Albany NY) 14, 8688–8699 (2022). 10.18632/aging.204380

97 Ying Yang, H. F., Xinyu Xu, Shankun Yao, Wenhao Yu and Zijian Guo. Zinc Ion-Induced Immune Responses in Antitumor Immunotherapy. CCS Chemistry 0, 1–20 (2024). 10.31635/ccschem.023.202303514

98 Sugandha Narayan, R. D., Zaigham Abbas Rizvi, Amit Awasthi. Zinc dampens anti-tumor immunity by promoting Foxp3+ regulatory T cells. Front Immunol 15 (2024). 10.3389/fimmu.2024.1389387

99 Chang, W., Li, H., Ou, W. & Wang, S. Y. A novel zinc metabolism-related gene signature to predict prognosis and immunotherapy response in lung adenocarcinoma. Front Immunol 14, 1147528 (2023). 10.3389/fimmu.2023.1147528

100 Lim, B., Kim, K. S. & Na, K. pH-Responsive Zinc Ion Regulating Immunomodulatory Nanoparticles for Effective Cancer Immunotherapy. Biomacromolecules 24, 4263–4273 (2023). 10.1021/acs.biomac.3c00557

101 Kreuger, I. Z. M., Slieker, R. C., van Groningen, T. & van Doorn, R. Therapeutic Strategies for Targeting CDKN2A Loss in Melanoma. J Invest Dermatol 143, 18–25 e11 (2023). 10.1016/j.jid.2022.07.016

102 Young, R. J. et al. Loss of CDKN2A expression is a frequent event in primary invasive melanoma and correlates with sensitivity to the CDK4/6 inhibitor PD0332991 in melanoma cell lines. Pigment Cell Melanoma Res 27, 590–600 (2014). 10.1111/pcmr.12228

103 Grard, M. et al. Homozygous Co-Deletion of Type I Interferons and CDKN2A Genes in Thoracic Cancers: Potential Consequences for Therapy. Front Oncol 11, 695770 (2021). 10.3389/fonc.2021.695770

104 Barriga, F. M. et al. MACHETE identifies interferon-encompassing chromosome 9p21.3 deletions as mediators of immune evasion and metastasis. Nat Cancer 3, 1367–1385 (2022). 10.1038/s43018-022-00443-5

105 Tiwari, A., Trivedi, R. & Lin, S. Y. Tumor microenvironment: barrier or opportunity towards effective cancer therapy. J Biomed Sci 29, 83 (2022). 10.1186/s12929-022-00866-3

106 Kather, J. N. et al. Topography of cancer-associated immune cells in human solid tumors. Elife 7 (2018). 10.7554/eLife.36967

107 Meyer, C. et al. Frequencies of circulating MDSC correlate with clinical outcome of melanoma patients treated with ipilimumab. Cancer Immunol Immunother 63, 247–257 (2014). 10.1007/s00262-013-1508-5

108 Riaz, N. et al. Tumor and Microenvironment Evolution during Immunotherapy with Nivolumab. Cell 171, 934–949 e916 (2017). 10.1016/j.cell.2017.09.028

109 Zhu, Y. et al. Disruption of tumour-associated macrophage trafficking by the osteopontin-induced colony-stimulating factor-1 signalling sensitises hepatocellular carcinoma to anti-PD-L1 blockade. Gut 68, 1653–1666 (2019). 10.1136/gutjnl-2019-318419

110 Xu, J. et al. CSF1R signaling blockade stanches tumor-infiltrating myeloid cells and improves the efficacy of radiotherapy in prostate cancer. Cancer Res 73, 2782–2794 (2013). 10.1158/0008-5472.CAN-12-3981

111 Stafford, J. H. et al. Colony stimulating factor 1 receptor inhibition delays recurrence of glioblastoma after radiation by altering myeloid cell recruitment and polarization. Neuro Oncol 18, 797–806 (2016). 10.1093/neuonc/nov272

112 Freeman, G. J. et al. Engagement of the PD-1 immunoinhibitory receptor by a novel B7 family member leads to negative regulation of lymphocyte activation. J Exp Med 192, 1027–1034 (2000). 10.1084/jem.192.7.1027

113 Hirano, F. et al. Blockade of B7-H1 and PD-1 by monoclonal antibodies potentiates cancer therapeutic immunity. Cancer Res 65, 1089–1096 (2005).

114 Dhupkar, P., Gordon, N., Stewart, J. & Kleinerman, E. S. Anti-PD-1 therapy redirects macrophages from an M2 to an M1 phenotype inducing regression of OS lung metastases. Cancer Med 7, 2654–2664 (2018). 10.1002/cam4.1518

115 Xu, C. et al. Protocol for detecting macrophage-mediated cancer cell phagocytosis in vitro and in vivo. STAR Protoc 4, 101940 (2023). 10.1016/j.xpro.2022.101940

116 Vitale, I., Manic, G., Coussens, L. M., Kroemer, G. & Galluzzi, L. Macrophages and Metabolism in the Tumor Microenvironment. Cell Metab 30, 36–50 (2019). 10.1016/j.cmet.2019.06.001

117 Tangudu, N. K. et al. ATR promotes mTORC1 activation via de novo cholesterol synthesis in p16-low cancer cells. bioRxiv (2023). 10.1101/2023.10.27.564195

118 Deleye, Y. et al. CDKN2A/p16INK4a suppresses hepatic fatty acid oxidation through the AMPKalpha2-SIRT1-PPARalpha signaling pathway. J Biol Chem 295, 17310–17322 (2020). 10.1074/jbc.RA120.012543

119 Bantubungi, K. et al. Cdkn2a/p16Ink4a regulates fasting-induced hepatic gluconeogenesis through the PKA-CREB-PGC1alpha pathway. Diabetes 63, 3199–3209 (2014). 10.2337/db13-1921

120 Khan, A., Liu, Y., Gad, M., Kenny, T. C. & Birsoy, K. Solute carriers: The gatekeepers of metabolism. Cell 188, 869–884 (2025). 10.1016/j.cell.2025.01.015

121 Sage, M. A. G. & Duffy, D. M. Novel Plasma Membrane Androgen Receptor SLC39A9 Mediates Ovulatory Changes in Cells of the Monkey Ovarian Follicle. Endocrinology 165 (2024). 10.1210/endocr/bqae071

122 Converse, A. & Thomas, P. The zinc transporter ZIP9 (Slc39a9) regulates zinc dynamics essential to egg activation in zebrafish. Sci Rep 10, 15673 (2020). 10.1038/s41598-020-72515-4

123 Thomas, P. & Dong, J. (-)-Epicatechin acts as a potent agonist of the membrane androgen receptor, ZIP9 (SLC39A9), to promote apoptosis of breast and prostate cancer cells. The Journal of steroid biochemistry and molecular biology 211, 105906 (2021). 10.1016/j.jsbmb.2021.105906

124 Thomas, P., Pang, Y. & Dong, J. Membrane androgen receptor characteristics of human ZIP9 (SLC39A) zinc transporter in prostate cancer cells: Androgen-specific activation and involvement of an inhibitory G protein in zinc and MAP kinase signaling. Mol Cell Endocrinol 447, 23–34 (2017). 10.1016/j.mce.2017.02.025

125 Matsuura, W. et al. SLC39A9 (ZIP9) regulates zinc homeostasis in the secretory pathway: characterization of the ZIP subfamily I protein in vertebrate cells. Biosci Biotechnol Biochem 73, 1142–1148 (2009). 10.1271/bbb.80910

126 Yamaji, T. et al. A CRISPR Screen Using Subtilase Cytotoxin Identifies SLC39A9 as a Glycan-Regulating Factor. iScience 15, 407–420 (2019). 10.1016/j.isci.2019.05.005

127 Zhang, T. et al. Crystal structures of a ZIP zinc transporter reveal a binuclear metal center in the transport pathway. Sci Adv 3, e1700344 (2017). 10.1126/sciadv.1700344

128 Bulldan, A., Malviya, V. N., Upmanyu, N., Konrad, L. & Scheiner-Bobis, G. Testosterone/bicalutamide antagonism at the predicted extracellular androgen binding site of ZIP9. Biochim Biophys Acta Mol Cell Res 1864, 2402–2414 (2017). 10.1016/j.bbamcr.2017.09.012

129 Bukiya, A. N. & Dopico, A. M. Common structural features of cholesterol binding sites in crystallized soluble proteins. J Lipid Res 58, 1044–1054 (2017). 10.1194/jlr.R073452

130 Atger, V. M. et al. Cyclodextrins as catalysts for the removal of cholesterol from macrophage foam cells. J Clin Invest 99, 773–780 (1997). 10.1172/JCI119223

131 Zidovetzki, R. & Levitan, I. Use of cyclodextrins to manipulate plasma membrane cholesterol content: evidence, misconceptions and control strategies. Biochim Biophys Acta 1768, 1311–1324 (2007). 10.1016/j.bbamem.2007.03.026

132 Xiong, Y., Zhang, H. & Beach, D. Subunit rearrangement of the cyclin-dependent kinases is associated with cellular transformation. Genes Dev 7, 1572–1583 (1993). 10.1101/gad.7.8.1572

133. 133 (!!! INVALID CITATION !!! 12,13,31,92-96,107-111).

134 Bian, Y. et al. Cancer SLC43A2 alters T cell methionine metabolism and histone methylation. Nature 585, 277–282 (2020). 10.1038/s41586-020-2682-1

135 Ho, P. C. et al. Phosphoenolpyruvate Is a Metabolic Checkpoint of Anti-tumor T Cell Responses. Cell 162, 1217–1228 (2015). 10.1016/j.cell.2015.08.012

136 Chang, C. H. et al. Metabolic Competition in the Tumor Microenvironment Is a Driver of Cancer Progression. Cell 162, 1229–1241 (2015). 10.1016/j.cell.2015.08.016

137 Li, Z. et al. SLC3A2 promotes tumor-associated macrophage polarization through metabolic reprogramming in lung cancer. Cancer Sci 114, 2306–2317 (2023). 10.1111/cas.15760

138 Kyritsi, K. et al. Downregulation of p16 Decreases Biliary Damage and Liver Fibrosis in the Mdr2(/) Mouse Model of Primary Sclerosing Cholangitis. Gene Expr 20, 89–103 (2020). 10.3727/105221620X15889714507961

139 Cudejko, C. et al. p16INK4a deficiency promotes IL-4-induced polarization and inhibits proinflammatory signaling in macrophages. Blood 118, 2556–2566 (2011). 10.1182/blood-2010-10-313106

140 Fuentes, L. et al. Downregulation of the tumour suppressor p16INK4A contributes to the polarisation of human macrophages toward an adipose tissue macrophage (ATM)-like phenotype. Diabetologia 54, 3150–3156 (2011). 10.1007/s00125-011-2324-0

141 Liang, Y. et al. p16(INK4a) Plays Critical Role in Exacerbating Inflammaging in High Fat Diet Induced Skin. Oxid Med Cell Longev 2022, 3415528 (2022). 10.1155/2022/3415528

142 Che, H. et al. p16 deficiency attenuates intervertebral disc degeneration by adjusting oxidative stress and nucleus pulposus cell cycle. eLife 9 (2020). 10.7554/eLife.52570

143 Zhu, Z., Song, H. & Xu, J. CDKN2A Deletion in Melanoma Excludes T Cell Infiltration by Repressing Chemokine Expression in a Cell Cycle-Dependent Manner. Front Oncol 11, 641077 (2021). 10.3389/fonc.2021.641077

144 Balli, D., Rech, A. J., Stanger, B. Z. & Vonderheide, R. H. Immune Cytolytic Activity Stratifies Molecular Subsets of Human Pancreatic Cancer. Clin Cancer Res 23, 3129–3138 (2017). 10.1158/1078-0432.CCR-16-2128

145 Wartenberg, M. et al. Integrated Genomic and Immunophenotypic Classification of Pancreatic Cancer Reveals Three Distinct Subtypes with Prognostic/Predictive Significance. Clin Cancer Res 24, 4444–4454 (2018). 10.1158/1078-0432.CCR-17-3401

146 Morrison, C. et al. Predicting response to checkpoint inhibitors in melanoma beyond PD-L1 and mutational burden. J Immunother Cancer 6, 32 (2018). 10.1186/s40425-018-0344-8

147 Nowak, J. E., Harmon, K., Caldwell, C. C. & Wong, H. R. Prophylactic zinc supplementation reduces bacterial load and improves survival in a murine model of sepsis. Pediatr Crit Care Med 13, e323–329 (2012). 10.1097/PCC.0b013e31824fbd90

148 Brazao, V. et al. Zinc supplementation increases resistance to experimental infection by Trypanosoma cruzi. Vet Parasitol 154, 32–37 (2008). 10.1016/j.vetpar.2008.02.015

149 Humphrey, P. A., Ashraf, M. & Lee, C. M. Hepatic cells’ mitotic and peritoneal macrophage phagocytic activities during Trypanosoma musculi infection in zinc-deficient mice. J Natl Med Assoc 89, 259–267 (1997).

150 Salvin, S. B., Horecker, B. L., Pan, L. X. & Rabin, B. S. The effect of dietary zinc and prothymosin alpha on cellular immune responses of RF/J mice. Clin Immunol Immunopathol 43, 281–288 (1987). 10.1016/0090-1229(87)90137-1

151 Nabeel-Shah, S. et al. C2H2-zinc-finger transcription factors bind RNA and function in diverse post-transcriptional regulatory processes. Mol Cell 84, 3810–3825 e3810 (2024). 10.1016/j.molcel.2024.08.037

152 Jackson, K. A., Valentine, R. A., Coneyworth, L. J., Mathers, J. C. & Ford, D. Mechanisms of mammalian zinc-regulated gene expression. Biochem Soc Trans 36, 1262–1266 (2008). 10.1042/BST0361262

153 Newcomer, K. et al. Malignant melanoma: evolving practice management in an era of increasingly effective systemic therapies. Curr Probl Surg 59, 101030 (2022). 10.1016/j.cpsurg.2021.101030

154 Clinton, L. K. et al. The LAST guidelines in clinical practice: implementing recommendations for p16 use. Am J Clin Pathol 144, 844–849 (2015). 10.1309/AJCPUXLP7XD8OQYY

155 Chan, S. H., Chiang, J. & Ngeow, J. CDKN2A germline alterations and the relevance of genotype-phenotype associations in cancer predisposition. Hered Cancer Clin Pract 19, 21 (2021). 10.1186/s13053-021-00178-x

156 Fraker, P. J. & King, L. E. Reprogramming of the immune system during zinc deficiency. Annu Rev Nutr 24, 277–298 (2004). 10.1146/annurev.nutr.24.012003.132454

157 Costello, L. C. & Franklin, R. B. Cytotoxic/tumor suppressor role of zinc for the treatment of cancer: an enigma and an opportunity. Expert Rev Anticancer Ther 12, 121–128 (2012). 10.1586/era.11.190

158 Mao, W. et al. Statin shapes inflamed tumor microenvironment and enhances immune checkpoint blockade in non-small cell lung cancer. JCI Insight 7 (2022). 10.1172/jci.insight.161940

159 Cantini, L. et al. High-intensity statins are associated with improved clinical activity of PD-1 inhibitors in malignant pleural mesothelioma and advanced non-small cell lung cancer patients. Eur J Cancer 144, 41–48 (2021). 10.1016/j.ejca.2020.10.031

160 Cortellini, A. et al. Integrated analysis of concomitant medications and oncological outcomes from PD-1/PD-L1 checkpoint inhibitors in clinical practice. J Immunother Cancer 8 (2020). 10.1136/jitc-2020-001361

161 Kansal, V. et al. Statin drugs enhance responses to immune checkpoint blockade in head and neck cancer models. J Immunother Cancer 11 (2023). 10.1136/jitc-2022-005940

162 Takada, K. et al. A propensity score-matched analysis of the impact of statin therapy on the outcomes of patients with non-small-cell lung cancer receiving anti-PD-1 monotherapy: a multicenter retrospective study. BMC Cancer 22, 503 (2022). 10.1186/s12885-022-09385-8

163 Liao, Y., Lin, Y., Ye, X. & Shen, J. Concomitant Statin Use and Survival in Patients With Cancer on Immune Checkpoint Inhibitors: A Meta-Analysis. JCO Oncol Pract, OP2400583 (2025). 10.1200/OP-24-00583

164 Wu, J. Y. et al. Cancer-Derived Succinate Promotes Macrophage Polarization and Cancer Metastasis via Succinate Receptor. Mol Cell 77, 213–227 e215 (2020). 10.1016/j.molcel.2019.10.023

165 Reinfeld, B. I. et al. Cell-programmed nutrient partitioning in the tumour microenvironment. Nature 593, 282–288 (2021). 10.1038/s41586-021-03442-1

166 Epstein, M. M. et al. Dietary zinc and prostate cancer survival in a Swedish cohort. Am J Clin Nutr 93, 586–593 (2011). 10.3945/ajcn.110.004804

167 Gonzalez, A., Peters, U., Lampe, J. W. & White, E. Zinc intake from supplements and diet and prostate cancer. Nutr Cancer 61, 206–215 (2009). 10.1080/01635580802419749

168 Li, L. & Gai, X. The association between dietary zinc intake and risk of pancreatic cancer: a meta-analysis. Biosci Rep 37 (2017). 10.1042/BSR20170155

169 Pan, S. Y. et al. Antioxidants and breast cancer risk-a population-based case-control study in Canada. BMC Cancer 11, 372 (2011). 10.1186/1471-2407-11-372

170 Adzersen, K. H., Jess, P., Freivogel, K. W., Gerhard, I. & Bastert, G. Raw and cooked vegetables, fruits, selected micronutrients, and breast cancer risk: a case-control study in Germany. Nutr Cancer 46, 131–137 (2003). 10.1207/S15327914NC4602_05

171 Lin, L. C. et al. Effects of zinc supplementation on clinical outcomes in patients receiving radiotherapy for head and neck cancers: a double-blinded randomized study. Int J Radiat Oncol Biol Phys 70, 368–373 (2008). 10.1016/j.ijrobp.2007.06.073

172 Shi, Z. et al. Association between dietary zinc intake and mortality among Chinese adults: findings from 10-year follow-up in the Jiangsu Nutrition Study. Eur J Nutr 57, 2839–2846 (2018). 10.1007/s00394-017-1551-7

173 Gutierrez-Gonzalez, E. et al. Dietary Zinc and Risk of Prostate Cancer in Spain: MCC-Spain Study. Nutrients 11 (2018). 10.3390/nu11010018

174 Zhang, Y., Song, M., Mucci, L. A. & Giovannucci, E. L. Zinc supplement use and risk of aggressive prostate cancer: a 30-year follow-up study. Eur J Epidemiol 37, 1251–1260 (2022). 10.1007/s10654-022-00922-0

175 Gallus, S. et al. Dietary zinc and prostate cancer risk: a case-control study from Italy. Eur Urol 52, 1052–1056 (2007). 10.1016/j.eururo.2007.01.094

176 Fernandez-Lazaro, C. I. et al. Dietary Antioxidant Vitamins and Minerals and Breast Cancer Risk: Prospective Results from the SUN Cohort. Antioxidants (Basel) 10 (2021). 10.3390/antiox10030340

177 Luo, H. et al. Association between Dietary Zinc and Selenium Intake, Oxidative Stress-Related Gene Polymorphism, and Colorectal Cancer Risk in Chinese Population - A Case-Control Study. Nutr Cancer 73, 1621–1630 (2021). 10.1080/01635581.2020.1804950

178 Bravi, F. et al. Dietary intake of selected micronutrients and the risk of pancreatic cancer: an Italian case-control study. Ann Oncol 22, 202–206 (2011). 10.1093/annonc/mdq302

179 Hoppe, C. et al. Zinc as a complementary treatment for cancer patients: a systematic review. Clin Exp Med 21, 297–313 (2021). 10.1007/s10238-020-00677-6

180 Trumbo, P., Yates, A. A., Schlicker, S. & Poos, M. Dietary reference intakes: vitamin A, vitamin K, arsenic, boron, chromium, copper, iodine, iron, manganese, molybdenum, nickel, silicon, vanadium, and zinc. J Am Diet Assoc 101, 294–301 (2001). 10.1016/S0002-8223(01)00078-5

181 Wang, K. et al. Zinc ions activate AKT and promote prostate cancer cell proliferation via disrupting AKT intramolecular interaction. Oncogene 44, 8–18 (2025). 10.1038/s41388-024-03195-x

182 Taylor, K. M. et al. ZIP7-mediated intracellular zinc transport contributes to aberrant growth factor signaling in antihormone-resistant breast cancer Cells. Endocrinology 149, 4912–4920 (2008). 10.1210/en.2008-0351

183 Lelliott, E. J. et al. Intracellular zinc protects tumours from T cell-mediated cytotoxicity. Cell Death Differ 31, 1707–1716 (2024). 10.1038/s41418-024-01369-4

184 Provinciali, M., Pierpaoli, E., Bartozzi, B. & Bernardini, G. Zinc Induces Apoptosis of Human Melanoma Cells, Increasing Reactive Oxygen Species, p53 and FAS Ligand. Anticancer Res 35, 5309–5316 (2015).

185 Xu, M. et al. Zinc-based radioenhancers to activate tumor radioimmunotherapy by PD-L1 and cGAS-STING pathway. J Nanobiotechnology 22, 782 (2024). 10.1186/s12951-024-02999-z

186 Du, M. & Chen, Z. J. DNA-induced liquid phase condensation of cGAS activates innate immune signaling. Science 361, 704–709 (2018). 10.1126/science.aat1022

187 Zhang, L. et al. A Peritumorally Injected Immunomodulating Adjuvant Elicits Robust and Safe Metalloimmunotherapy against Solid Tumors. Adv Mater 34, e2206915 (2022). 10.1002/adma.202206915

188 Sies, H. & Jones, D. P. Reactive oxygen species (ROS) as pleiotropic physiological signalling agents. Nat Rev Mol Cell Biol 21, 363–383 (2020). 10.1038/s41580-020-0230-3

189 Gide, T. N. et al. Distinct Immune Cell Populations Define Response to Anti-PD-1 Monotherapy and Anti-PD-1/Anti-CTLA-4 Combined Therapy. Cancer Cell 35, 238–255 e236 (2019). 10.1016/j.ccell.2019.01.003

190 Lee, J. H. et al. Transcriptional downregulation of MHC class I and melanoma de-differentiation in resistance to PD-1 inhibition. Nat Commun 11, 1897 (2020). 10.1038/s41467-020-15726-7

191 Hugo, W. et al. Genomic and Transcriptomic Features of Response to Anti-PD-1 Therapy in Metastatic Melanoma. Cell 165, 35–44 (2016). 10.1016/j.cell.2016.02.065

192 Ayers, M. et al. IFN-γ-related mRNA profile predicts clinical response to PD-1 blockade. The Journal of clinical investigation 127, 2930–2940 (2017). 10.1172/jci91190

193 Gyorffy, B. Integrated analysis of public datasets for the discovery and validation of survival-associated genes in solid tumors. Innovation (Camb) 5, 100625 (2024). 10.1016/j.xinn.2024.100625

194 Gyorffy, B. Transcriptome-level discovery of survival-associated biomarkers and therapy targets in non-small-cell lung cancer. Br J Pharmacol 181, 362–374 (2024). 10.1111/bph.16257

195 Kovacs, S. A., Fekete, J. T. & Gyorffy, B. Predictive biomarkers of immunotherapy response with pharmacological applications in solid tumors. Acta Pharmacol Sin 44, 1879–1889 (2023). 10.1038/s41401-023-01079-6

196 Jerby-Arnon, L. et al. A Cancer Cell Program Promotes T Cell Exclusion and Resistance to Checkpoint Blockade. Cell 175, 984–997 e924 (2018). 10.1016/j.cell.2018.09.006

197 Chen, Y. P. et al. Single-cell transcriptomics reveals regulators underlying immune cell diversity and immune subtypes associated with prognosis in nasopharyngeal carcinoma. Cell Res 30, 1024–1042 (2020). 10.1038/s41422-020-0374-x

198 Uphoff, C. C. & Drexler, H. G. Detection of mycoplasma contaminations. Methods Mol Biol 290, 13–23 (2005). 10.1385/1-59259-838-2:013

199 Sullivan, M. R. et al. Quantification of microenvironmental metabolites in murine cancers reveals determinants of tumor nutrient availability. Elife 8 (2019). 10.7554/eLife.44235

200 Corrin A. Wohlhieter et al. An optimized NGS sample preparation protocol for in vitro CRISPR screens. STAR Protocols 2, 100390 (2021). 10.1016/j.xpro.2021.100390

201 Alker, W., Schwerdtle, T., Schomburg, L. & Haase, H. A Zinpyr-1-based Fluorimetric Microassay for Free Zinc in Human Serum. Int J Mol Sci 20 (2019). 10.3390/ijms20164006

202 Stringer, C., Wang, T., Michaelos, M. & Pachitariu, M. Cellpose: a generalist algorithm for cellular segmentation. Nat Methods 18, 100–106 (2021). 10.1038/s41592-020-01018-x

203 Dimri, G. P. et al. A biomarker that identifies senescent human cells in culture and in aging skin in vivo. Proc Natl Acad Sci U S A 92, 9363–9367 (1995). 10.1073/pnas.92.20.9363

